# Accurately positioning functional residues with robotics-inspired computational protein design

**DOI:** 10.1101/2021.07.02.450934

**Authors:** Cody Krivacic, Kale Kundert, Xingjie Pan, Roland A. Pache, Lin Liu, Shane O Conchúir, Jeliazko R. Jeliazkov, Jeffrey J. Gray, Michael C. Thompson, James S. Fraser, Tanja Kortemme

## Abstract

Accurate positioning of functional residues is critical for the design of new protein functions, but has remained difficult because of the prevalence of irregular local geometries in active sites. Here we introduce two computational methods that build local protein geometries from sequence with atomic accuracy: fragment kinematic closure (FKIC) and loophash kinematic closure (LHKIC). FKIC and LHKIC integrate two approaches: robotics-inspired kinematics of protein backbones and insertion of peptide fragments, and show up to 140-fold improvements in native-like predictions over either approach alone. We then integrate these methods into a new design protocol, pull-into-place (PIP), to position functionally important sidechains via design of new structured loop conformations. We validate PIP by remodeling a sizeable active site region in an enzyme and confirming the engineered new conformations of two designs with crystal structures. The described methods can be applied broadly to the design of many new protein geometries and functions.

## INTRODUCTION

The ability to design proteins with new and useful functions has many applications in biotechnology and medicine. Computational methods have been successfully applied to the design of many new proteins with “idealized” stable structures^1^, but engineering new functions has remained a substantial challenge^2^. This discrepancy in design success can, at least partially, be attributed to several features that distinguish functional sites from idealized protein structures. First, functional sites are often highly sensitive to small errors in functional group positioning, and nonproductive conformations must be avoided for efficient catalysis. Second, cavities and charged functional surfaces in active sites can destabilize proteins considerably. Finally, many activities require multiple functional conformations separated by finely tuned energy barriers.

Here we focus specifically on the problem of positioning functionally important sidechains via design of new structured loop conformations. Loops make up 50% of residues in enzyme active sites, compared to just 30% of residues overall^3^. This prevalence of loops in active sites suggests that structured loops are well suited to supporting the bespoke geometries of sidechain and backbone functional groups that are often necessary for function. However, the large number of degrees of freedom, which enable loops to adopt geometries optimized for function, also makes loops difficult to design since the space of backbone conformations to consider is vast.

A handful of methods have been developed to address these challenges as reviewed previously^4^, including strategies utilizing the protein design program Rosetta^5^. Eiben et al. tasked players of the computer game Foldit with designing a loop region to better desolvate the active site of a computationally designed Diels-Alderase^6^. The players were successful, ultimately designing a 13-residue insertion that improved catalysis by 20-fold, in part via a *de novo* helix-turn-helix motif. This strategy is not generalizable, however, as it required human intervention to solve the loop design problem. Murphy et al. developed an algorithm to redesign an active site loop in human guanine deaminase^7^, and Borgo et al. developed a similar algorithm to redesign a substrate-binding loop in *E. coli* methionine aminopeptidase^8^. Both algorithms place a sidechain in the ideal position for function, enumerate backbone coordinates capable of supporting the sidechain and then remodel the backbone to intersect one or more of those coordinates. In particular, Murphy et al. automatically determined the ideal length of the redesigned loop, while Borgo et al. derived ideal sidechain positions from the Protein Data Bank (PDB) and simultaneously accommodated multiple sidechains. Unlike the Foldit players^6^, however, neither of these two computational methods made large rearrangements to the loop backbone or incorporated regions of secondary structure into the designed loop, placing a practical limit on the extent to which loops can be rearranged to adopt new functions.

In this study we introduce two robotics-inspired loop modeling methods, fragment-inspired kinematic closure (FKIC) and loophash-sampled kinematic closure (LHKIC), that can generate considerable structural changes and can include secondary structure elements. Each of these methods combines the robotics-inspired inverse kinematic closure algorithm (KIC) with a fragment-based search of conformational space, and are geared towards the design of new backbone geometries (LHKIC) and the prediction of backbone geometries from sequence (FKIC). We incorporate these methods into a new loop design protocol called Pull Into Place (PIP). The PIP protocol has three steps: (i) generation of new backbone conformations, where functional groups of interest are gently pulled towards their desired positions using harmonic restraints, (ii) sequence design using fixed-backbone side-chain optimizations with the same restraints, and (iii) structure prediction using unrestrained flexible-backbone simulations to identify designs predicted to adopt the desired new backbone conformation. We demonstrate that PIP is capable of accurately designing large segments of the protein backbone by remodeling an enzyme active site loop and confirming the conformations of two designs with crystal structures. Detailed characterization of one successful design reveals a robustness to mutation, suggesting that multiple interactions contribute to the conformation of the remodeled loop. Together, the techniques described in this paper will advance the engineering of functional proteins by allowing the design of user-defined new active site geometries.

## RESULTS

We set out to develop a method (PIP) to design new local backbone geometries in Rosetta to accurately position amino acid functional groups in a functional site. The PIP algorithm required 3 components: (1) a method to generate suitable and designable backbone conformations for positioning defined functional groups, (2) a way to stabilize these new backbone and side chain conformations by finding sequences optimal for the desired structure, and (3) a method to predict the new conformation given a sequence, to assess whether the desired structure is also optimal for the designed sequence.

We reasoned that robotics-inspired methods developed in our lab and incorporated into Rosetta^9,10^ would be suitable to generate conformations for (1), and design methods for (2) existed in Rosetta^11^. However, modeling structures given designed sequences represents a major bottleneck that becomes limiting especially when many designed sequences are to be evaluated and target structures do not exclusively adopt regular secondary structure geometries. We first describe two new methods that address this bottleneck and increase the efficiency of robotics-inspired sampling methods to predict and design new backbone conformations: fragment-kinematic closure (FKIC) and loophash-kinematic closure (LHKIC). We then describe the application of the entire PIP protocol to a design problem in which we reshape the backbone geometry of the active site in ketosteroid isomerase (KSI) to enable catalysis with a non-native residue.

### FKIC and LHKIC algorithms

FKIC and LHKIC integrate two concepts that have separately led to considerable advances in protein modeling: sampling preferred combinations of backbone torsions from fragments of proteins in the protein structure databank (PDB)^12^, and improved sampling of loop regions with an inverse kinematic closure algorithm termed ‘KIC’^9^ borrowed from the field of robotics^13^.

KIC determines ‘mechanically accessible’ conformations for internal protein segments of given lengths by sampling the phi/psi torsion degrees of freedom in the segment. Three Cα atoms of an N-residue segment are designated as pivots, leaving N-3 non-pivot Cα atoms. In the standard implementation of KIC in Rosetta^9^, non-pivot torsions are sampled from a residue type-specific Ramachandran map. A major bottleneck for KIC is that the number of possible conformations (combinations of torsion angles) increases dramatically with increasing numbers of residues to be modeled. This problem becomes especially limiting when predicting larger contiguous segments or entire functional sites which often contain both loop and secondary structure regions in several interacting segments; sampling of neighboring torsions from Ramachandran space is particularly non-ideal for secondary structure regions, where coupled torsions are unlikely to be sampled independently.

To enable these more challenging but also more realistic problems, we integrated KIC with fragment insertion^14^ in the Rosetta protein structure prediction and design program^5^. In the new FKIC structure prediction method, non-pivot degrees of freedom are taken from peptide fragments that are picked from the PDB using the sequence of the target loop^15^; KIC is then used to determine the values of the pivot torsions (**Fig. 1a**). We reasoned that FKIC would combine the improvements of KIC demonstrated previously^9^ with the reduction of degrees of freedom by using coupled torsion angles from fragments (in contrast to sampling all non-pivot torsions independently from Ramachandran space as in KIC), making FKIC particularly suitable for sampling local active site geometries.

**Figure 1.**
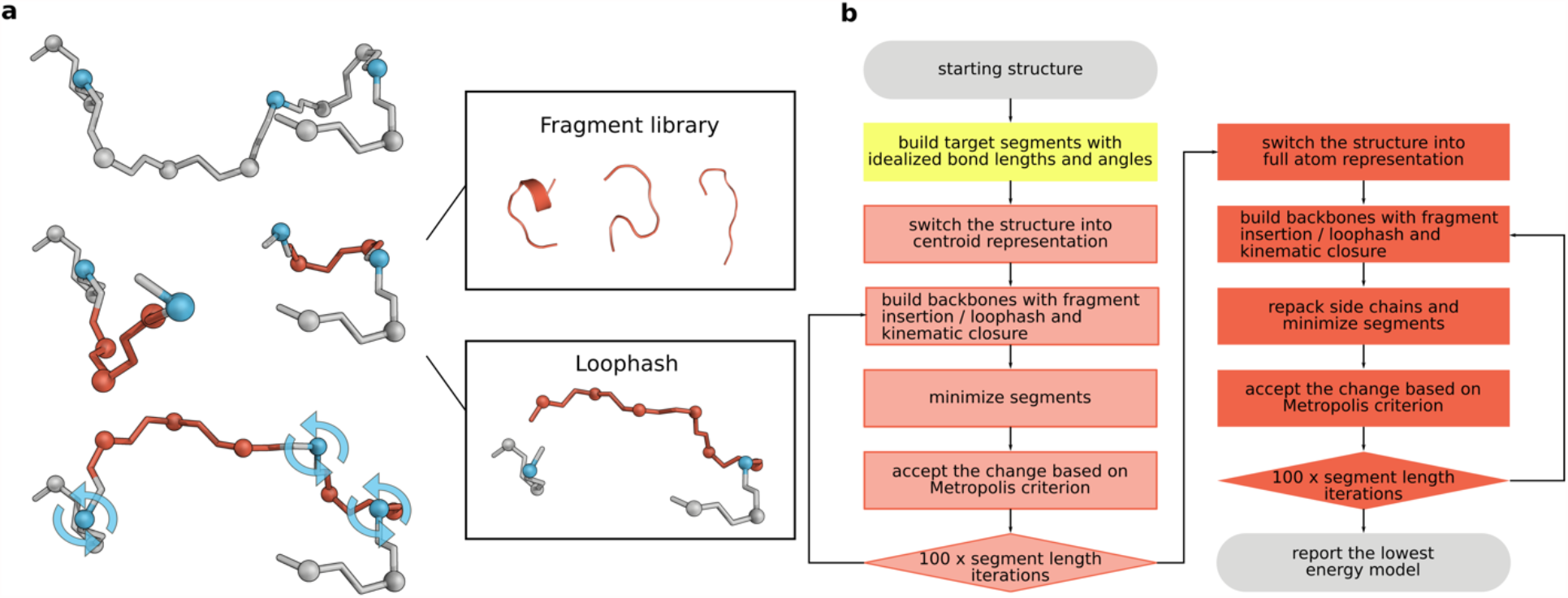
FKIC and LHKIC protocols for sampling protein backbone conformations. **(a)** Individual FKIC/LHKIC move. Three Cα atoms (blue) on the target segment to be modeled (grey) are picked randomly as pivots. Fragment insertion (FKIC) or loop hash (LHKIC) is applied to sample torsion degrees of freedom at non-pivot atoms (red), which breaks the chain. The KIC algorithm is then used to close the chain by determining appropriate values for the pivot torsions. **(b)** Integration of FKIC/LHKIC moves into a structure prediction protocol in Rosetta. Two stages of Monte Carlo simulated annealing - a beginning low resolution stage (centroid, light red) followed by a high-resolution stage (all-atom, red) - are used to find the lowest Rosetta energy conformation of target segments. For benchmarking, the native conformations of target segments and surrounding side chains are removed at the beginning of the simulation (**Methods**).

In design problems, the sequences of the target segments are variable during the sampling, which might limit the utility of FKIC for designing new segment conformations since the fragments in FKIC are picked based on sequence similarity to the starting structure. We therefore developed an additional method, LHKIC, which uses the loophash protocol^16^ to pick fragments that simultaneously sample structures and sequences of the target loops. The loophash protocol uses the 6D transformation between the residue before the first pivot and the residue after the last pivot as a query key to find peptide fragments from the PDB that approximately close the gap between these two residues (**Fig. 1a**). After insertion of a fragment, KIC determines the pivot torsions that close the gap and LHKIC mutates remodeled residues to the amino acids from the inserted fragment to improve local sequence-structure compatibility. Individual FKIC or LHKIC sampling moves (**Fig. 1a**) are then followed by optimization of side chain conformations in and around the altered backbone region and integrated into a Monte Carlo minimization protocol (**Fig. 1b, Supplementary Fig. 1**); sampled conformations are evaluated with Rosetta’s all-atom energy function^17,18^s.

### Loop structure prediction benchmarking

We tested the ability of FKIC to recapitulate the local conformations of protein segments, given their sequences, on a benchmark of 45 12-residue loops^19^, which were used previously to evaluate KIC^9^ (“Standard” set). We also constructed two new datasets designed to address the challenges of modeling active site regions highlighted above. The first new set contained 30 16-residue-segments where each segment contains both regular secondary structure elements and loop regions (“Mixed Segment” set). We reasoned that FKIC should improve the sampling of regular secondary structure elements by building them from fragments, whereas these elements might be difficult to model with KIC alone, since KIC samples each non-pivot torsion angle independently. The second new set contained 30 pairs of interacting 10-residue segments (“Multiple Segments” set). For all three benchmark sets, we discarded information on the native geometry of all target segments by building starting conformations with extended phi/psi torsion angles and idealized bond lengths and angles. All side chains within 10Å from the target segments were deleted and replaced with side chain conformations from a rotamer library^20^ during FKIC simulations. In all benchmark simulations, we also excluded fragments from structures homologous to the benchmark cases (Methods) to test FKIC’s ability to predict novel structures. For each test protein in each set, we generated 500 models with FKIC and calculated the backbone heavy atom root mean square deviation (RMSD) of each target segment after aligning the protein without the modeled segment to its crystal structure. We also applied control methods that use KIC and fragment insertion (CCD)^21^ alone to the same datasets using an otherwise identical protocol in Rosetta. For most comparisons, we replaced KIC with an improved KIC protocol called next-generation KIC (NGK)^10^. We used two performance metrics: The first quantifies prediction accuracy by determining the RMSD of the model with the lowest (best) Rosetta energy to each native structure and then taking the median RMSD value across each dataset. The second quantifies sampling performance by measuring the fraction of native-like (correct) models generated for each protein case, where native-like is defined as <1Å (“sub-Å”) RMSD to the native structure, and again taking the median for each dataset. We also measured the median run time to determine whether any increased sampling performance increases computational cost (**Methods** and **Supplementary Table 1a**).

The Rosetta KIC method had previously been shown^9^ to be comparable to a state-of-the-art molecular mechanics method^19^. The NGK update^10^ led to improved performance over KIC, and had comparable performance to GalaxyLoop-PS2^22^, RCD+^23^, Sphinx^24^, LEAP^25^ and FREAD^26,27^ when tested on identical datasets. Here we show that FKIC improves structure prediction accuracy over CCD, KIC and NGK, with the largest changes for the two new datasets (**Fig. 2a, top** and **Supplementary Table 1a**). On the 16-residue Mixed Segment dataset, which tests the ability of FKIC to predict conformations of protein segments with arbitrary secondary structure composition, the median accuracy improved to 0.53Å RMSD with FKIC compared to 1.29Å and 1.07Å with CCD and NGK alone, respectively. In 23/30 cases FKIC was able to identify conformations very close to the crystallographic structure (<1Å RMSD) as the lowest scoring model, compared to only 13 cases with CCD and 15 cases with NGK (**Supplementary Table 2**). For the Multiple Segments dataset, which tests the ability of FKIC to predict conformations of discontinuous interacting segments (a common feature of protein active sites), FKIC was the only method that yielded atomic (1Å) median accuracy, compared to 1.97Å and 1.29Å with CCD and NGK alone, respectively (**Fig. 2a, top** and **Supplementary Table 1a**). Representative examples where FKIC correctly predicted protein conformations while NGK failed are shown for the Mixed Segment and Multiple Segments datasets in **Figs. 2b,c** and details are given in **Supplementary Tables 2** and **3**. The improvements on the Standard dataset were smaller (median RMSD was 0.62Å with FKIC compared to 0.64Å for NGK, **Supplementary Table 1a**), but for 35/45 proteins FKIC finds lower energy structures than NGK (**Supplementary Table 4**). Cases where FKIC predictions did not lead to the identification of sub-Å accuracy lowest-scoring models can be attributed to both sampling and energy function limitations (**Supplementary Table 5, Supplementary Fig. 2**, and **Supplementary Note 1**). LHKIC performed similarly to FKIC when tested on the standard dataset (**Fig, 2a, Supplementary Table 1a**, and **Supplementary Note 2**).

**Figure 2.**
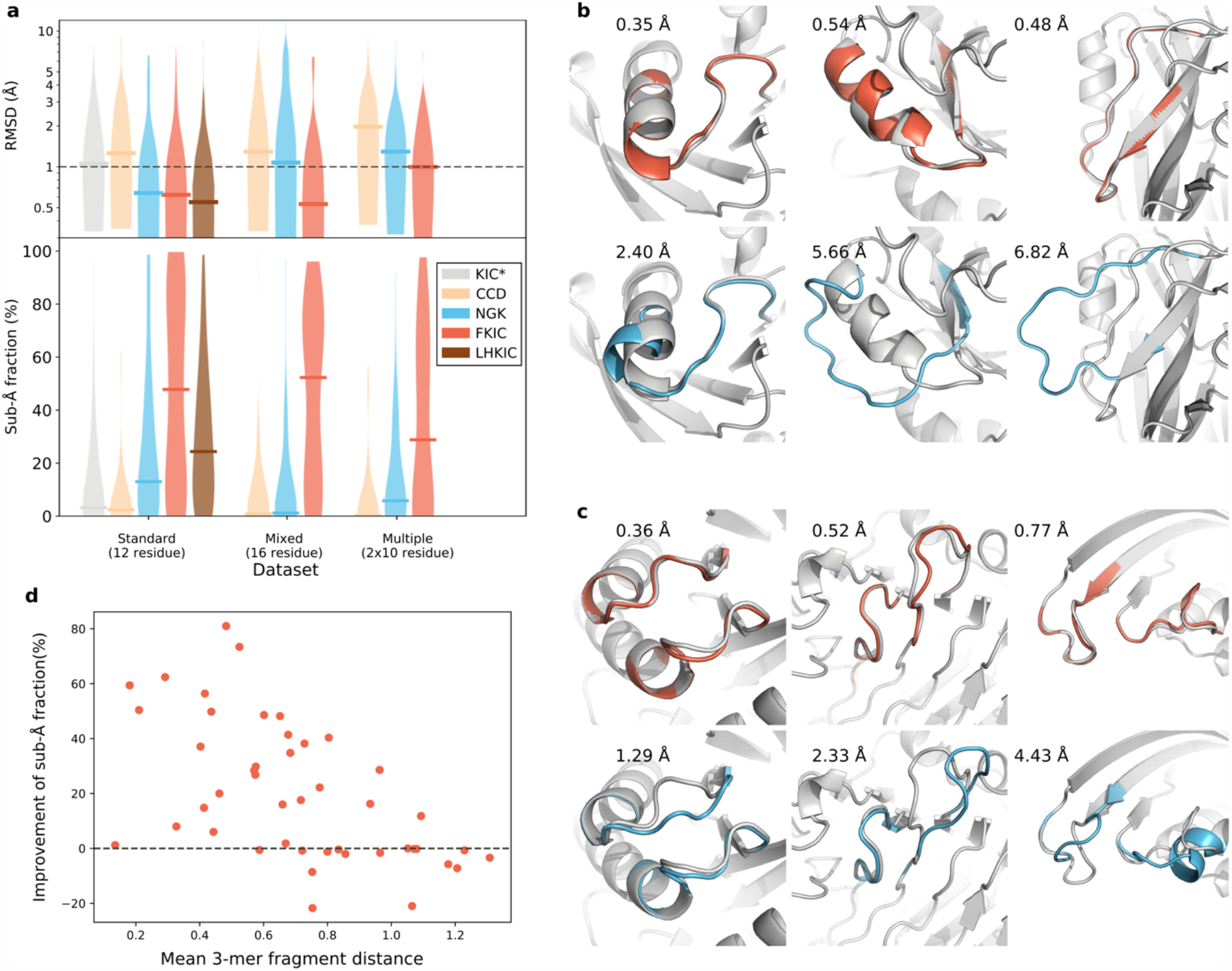
FKIC improves structure prediction accuracy. **(a)** Comparison of performance of different methods for three datasets: (i) Standard dataset described in ref.^9^, and 2 new sets: (ii) A “Mixed Segment” dataset with 30 16-residue regions that contain both loops and segments of regular secondary structure and (iii) a “Multiple Segments’’ dataset of 30 cases with 2 separate 10-residue regions that are interacting. KIC^9^: grey; CCD^21^: orange; NGK^10^: blue; FKIC: red, LHKIC: brown. Upper panel: violin plot of RMSD of lowest energy (best) model across each dataset. Horizontal bars indicate the median lowest-energy RMSD. FKIC is the only method that provides predictions with atomic accuracy (≤ 1Å median RMSD) for all datasets. Lower panel: violin plot of fraction of predicted models in each dataset that have sub-Å accuracy. FKIC leads to considerable improvements over previous methods. Asterisk indicates data from ref.^36^; all other simulations were run with the ref2015 Rosetta energy function^17^; methods using fragments (CCD and FKIC) used identical fragment libraries that excluded fragments from structural homologs to the target proteins. **(b**,**c)** FKIC correctly reconstructs geometries where the previous state-of-the-art method, NGK, fails. Shown are examples from the Mixed Segment **(b)** and Multiple Segments dataset **(c)**. Experimentally determined structures: grey; predictions from FKIC: red, top; predictions from NGK: blue, bottom. RMSDs to the experimentally determined structures are given in each panel in Å. **(d)** The fraction of sub-Å predictions is negatively correlated with the mean 3-mer fragment distance (**Methods**). Each data point represents a protein from the standard 12-residue dataset.

The second notable advance with FKIC, in addition to the improvements in accuracy described above, is how frequently FKIC generated conformations that are <1Å RMSD from the crystallographic conformation (**Fig. 2a, bottom**). For the Mixed Segment set, the median fraction of sub-Å predictions for FKIC was 52.3%, which was 45- and 105-fold higher than for NGK and CCD, respectively. For the Multiple Segments dataset, the median fraction of sub-Å predictions was 28.5% with FKIC, which was 5-fold higher than with NGK (5.5%) and 143-fold higher (0.2%) than with CCD (**Supplementary Table 1a**). In several cases, FKIC was able to find correct solutions for even larger conformational sampling problems such as a set with 2 interacting 12-residue segments (**Supplementary Note 3** and **Supplementary Tables 6-7**).

These improvements in sampling efficiency are important in particular for design, since they reduce the computational time needed to predict the conformation of a designed segment, allowing for more designs to be evaluated. Overall, the improvement of the fraction of sub-Å predictions is negatively correlated with the mean 3-mer fragment distance from the native structure (**Fig. 2d, Methods**). This observation shows that high quality fragments focus the sampling on native-like conformations.

In all benchmark simulations described above, we excluded fragments from structures homologous to the benchmark cases. As expected, both prediction accuracy and the median fraction of sub-Å predictions improved further with fragments from homologs present in the database (**Supplementary Table 1b**). FKIC might therefore be able to accurately sample even larger regions in cases where homologous structures are available. Taken together, we conclude that FKIC drastically increases sampling efficiency in the prediction of protein segments from sequence.

### Application of the PIP protocol

With improved methods for sampling and prediction of backbone conformations in hand, we set out to test the entire PIP protocol in a design application. We chose *Pseudomonas testosteroni* ketosteroid isomerase (KSI) as a model system. In the KSI active site, a catalytic aspartate at position 38 abstracts a proton from a steroid substrate to catalyze an energetically favorable double-bond rearrangement. When using 5(10)-estrene-3,17-dione as a substrate (selected for the absorbance change caused by the KSI-catalyzed double-bond rearrangement), we observe that mutating aspartate 38 in KSI to a glutamate reduces the protein’s k_cat_ by approximately 104-fold (**Table 1**), similar to previous work that reported a reduction of 240-fold in the D38E mutant compared wild-type^28^. This reduction in k_cat_ was attributed to the misplacement of the side chain carboxyl group that is common to glutamate and aspartate due to the additional carbon-carbon bond in the glutamate sidechain. Because of this sensitivity to perturbations on the length scale of a carbon-carbon bond, we reasoned that KSI was a good model system to test the ability of PIP to accurately position functional groups. In particular, we set out to replace aspartate with glutamate while maintaining the precise placement of the side chain carboxyl group (**Fig. 3a**) by reshaping a sizable region of the protein backbone (11-12 residues, **Fig. 3b**). No known homologs of KSI contain a glutamate at the catalytic position^29^. Thus, any designed solutions would be novel and a fragment-based design protocol would not be able to rely on naturally occurring homologs that have already solved this particular problem.

**Table 1.** Kinetic parameters of wild-type (WT) KSI, WT D38E, and designs. Ranges are based on the standard deviation of three independent experiments.

| Enzyme | $k_{\text{cat}}$ ($\text{min}^{-1}$) | $K_{\text{M}}$ ($\mu\text{M}$) | $k_{\text{cat}}/K_{\text{M}}$ ($\mu\text{M}^{-1} \text{min}^{-1}$) |
| --- | --- | --- | --- |
| WT | $350 \pm 18$ | $120 \pm 32$ | $2.9 \pm 0.79$ |
| WT D38E | $3.4 \pm 0.50$ | $37 \pm 4.9$ | $0.092 \pm 0.018$ |
| V1D8r | $1.7 \pm 0.41$ | $67 \pm 15$ | $0.025 \pm 0.0084$ |
| V2D9r | $0.29 \pm 0.0040$ | $9.0 \pm 2.0$ | $0.032 \pm 0.0084$ |

**Figure 3.**
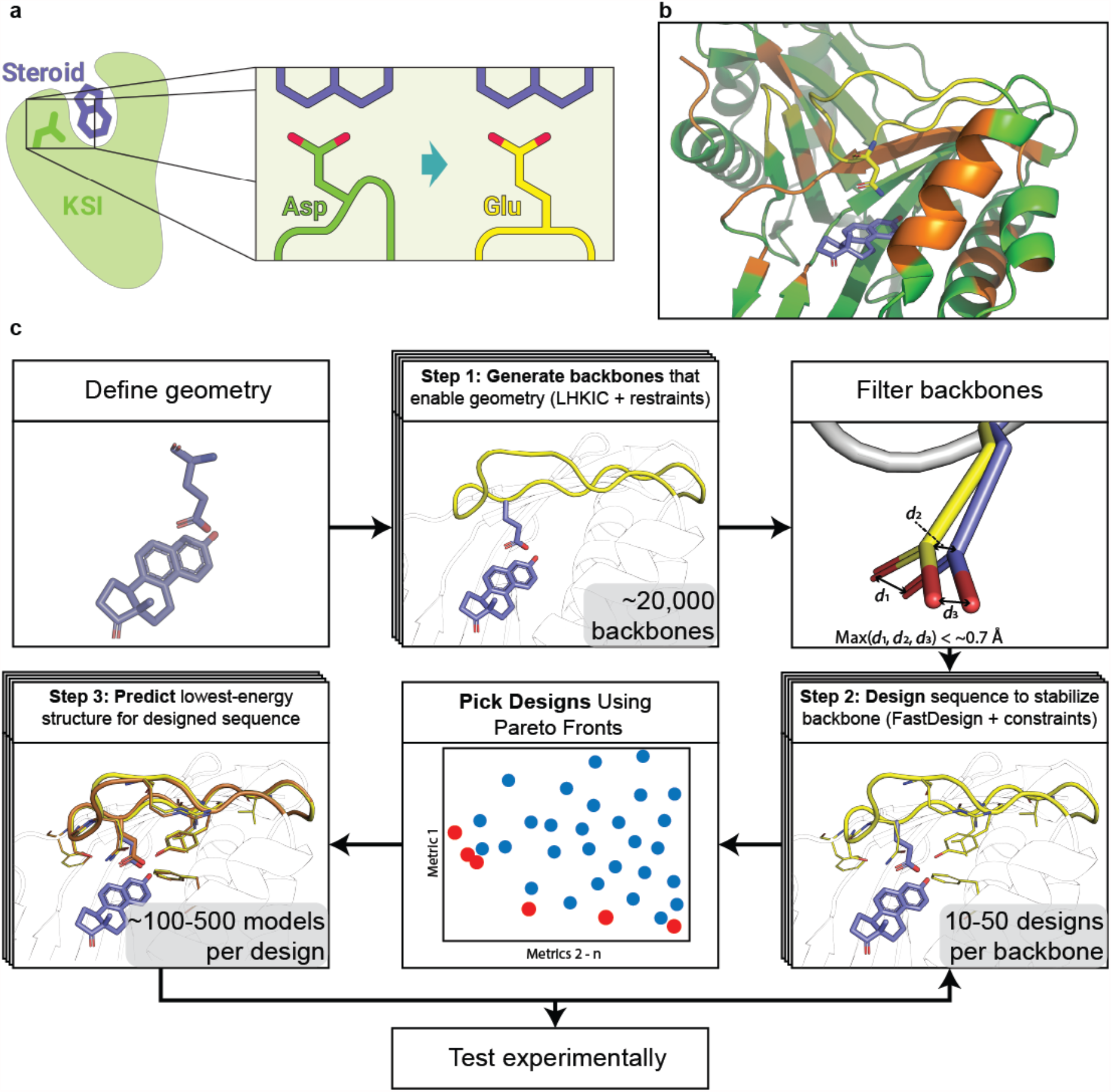
PIP design protocol applied to ketosteroid isomerase (KSI). **(a)** Schematic of design goal for KSI. Green: wild-type KSI with catalytic aspartate. Yellow: Designed KSI variant with reshaped active site to position the glutamate carboxyl group in place of the wild-type aspartate carboxyl group. **(b)** KSI wild-type structure (PDB 1QJG), showing the active site regions to be remodeled in yellow. Residues allowed to change identity (design) or conformation (repack) during the design process (PIP version 2) are shown in yellow or orange, respectively, and static positions are shown in green. **(c)** Steps of the PIP protocol applied to KSI. Top left: functional geometry is defined. Top middle: new backbone conformations (yellow) are built to satisfy the geometric restraints. Top right: Backbones are filtered based on their ability to satisfy the geometric restraints. d1, d2, and d3 refer to the distances of the atoms in the carboxyl group to their defined ideal positions. Bottom right: Sequences are designed to stabilize the *de novo* backbone. Bottom middle: Pareto fronts are used to select designs for structure prediction. Blue: all designs, red: Pareto-efficient designs. Bottom left: The lowest-energy structure (orange) is predicted using loop modeling methods in Rosetta.

Our PIP design protocol for KSI (**Fig. 3c**) proceeded in three steps: In step 1, we built 20,000 *de novo* backbone conformations that positioned the functional carboxyl group using harmonic coordinate restraints defined by the amide atoms of asparagine 38 (an inactivating mutation for the catalytic D38 that enables a transition state mimic to be crystallized) in PDB file 1QJG in place of the catalytic D38) (**Fig. 3c**). We selected the 1,600-4,000 conformations that best matched the desired geometry based on their restraint satisfaction, which we defined as the maximum distance of any restrained atom in the model to the atom’s ideal position (**Fig.3c**, top right panel).

In the second step, these new backbone conformations were stabilized by redesigning the local environment, where all residues of the new backbone segments as well as residues in the environment were redesigned using design methods in Rosetta (see **Methods**). This process resulted in 10-50 designs per input structure. We then selected 200-422 design models for structure prediction in step 3. These designs were selected based on how close the catalytic residue carboxyl group atoms were to their desired positions, and several computational design quality metrics (see below and **Methods**).

While step 2 (design) aims to find sequences that are optimal for the targeted new conformations, step 3 (structure prediction) aims to assess whether these sequences indeed fold into the targeted conformation (i.e. is the conformation also optimal given the sequence). Steps 2 and 3 were iterated to further optimize sequence-structure combinations. In particular, designed sequences that produced structure prediction models that correctly placed the functional carboxyl group but are not the lowest-scoring model generated by the structure prediction protocol were fed back to step 2 for further sequence optimization.

We created designs using two versions of the PIP protocol, denoted versions 1 and 2, which differed in several details. Version 1 was developed before FKIC and LHKIC, and therefore used NGK for both model generation (step 1) and structure prediction (step 3). In step 1, we varied the length of the active site loop from 0 to -6 residues (relative to its native length). In subsequent steps, we made comparisons only between loops of the same length, to avoid biases towards longer loops that can make more favorable interactions at the expense of loss of conformational entropy not considered in Rosetta. Sequence design (step 2) was done using fixed-backbone rotamer sampling. Residues within 4Å of the active site loop were designed (i.e. allowed to change amino acid identity, see **Methods**) excluding residues Y14, F54, D99, A114 and F116 that are important for catalysis. In total, 33-39 residues were allowed to design, depending on loop length. Designs from step 2 to be evaluated in step 3 were selected with a probability proportional to their Boltzmann-weighted Rosetta scores (see **Methods**). This approach was intended to improve the diversity of the selected designs, while still selecting more favorable (low-scoring) designs. The designs selected for experimental testing either retained the native loop length or shortened the loop by one residue.

In version 2, we made several changes: We used LHKIC for model generation (step 1) and FKIC for structure prediction (step 3). This strategy takes advantage of the ability of LHKIC to sample both sequence and structure simultaneously in step 1 (as fragment picking in LHKIC is independent of the starting sequence). Conversely, FKIC is better suited to predicting conformations given a sequence in step 3, since FKIC picks fragments based on the input sequence. In step 2, we incorporated a small degree of backbone flexibility into the design process by using the Rosetta FastDesign method, which iterates fixed-backbone sequence design and fixed-sequence structure minimization. Because this design algorithm is more computationally expensive than that from version 1, we made fewer designs per backbone model (10 instead of 50). Based on the results from version 1, we only considered two loop lengths: the native length and a one-residue deletion. We also allowed a different (and smaller) set of residues to design: 25 – 26 residues in the active site loop and 4 residues in a small β-hairpin (residues 74 – 77 in the dimer partner) that make inter-chain contacts with the active site loop. To select designs for step 3, we incorporated knowledge from additional metrics besides Rosetta score and functional group positioning. We used metrics including the number of buried unsatisfied and oversaturated hydrogen bonds, a fragment quality filter, total solvent-accessible surface area, and Rosetta’s foldability metric (**Methods**). Because it is unclear *a priori* how to prioritize these metrics, we used Pareto fronts consisting of the above metrics to choose designs for computational structure prediction (**Fig. 3c, Supplementary Fig. 3**). We also selected more designs than in version 1 (up to 422 instead of 200) for structure prediction in early iterations of step 3, taking advantage of the fact that FKIC requires fewer simulations than NGK to make sub-Ångstrom predictions.

### Selection of designed KSI variants

We selected 32 designs for experimental testing, 14 from version 1 and 22 from version 2. Designs were named according to the version of PIP used to create them (V1 and V2) and a design number (D1, D2, …) (**Supplementary Tables 8-9, Supplementary Figs. 4-5**). Designs appended with “r” indicate that mutations were reverted to the wild-type residue based on visual inspection; 8/14 designs from version 1 contained reversion mutations (**Supplementary Tables 8-9)**, and 11/22 designs from version 2 contained reversion mutations (for version 2, **Supplementary Table 8 and Supplementary Figs. 4-5** show the 11 original computational designs, with the mutations for 7 additional reversion designs listed in **Supplementary Table 9**). Design selection was based on a number of factors: We chose designs that maximized the gap in Rosetta total (version 1) or fa_attr (version 2) score between models that correctly place the catalytic residue (<1 Å restraint satisfaction, defined as the maximum distance between a restrained atom and its defined position) and models which do not correctly position the catalytic residue (>2 Å restraint satisfaction). We also chose designs that were predicted to have few buried unsatisfied hydrogen bonds, and that did not have significant sequence and structural similarity to previously selected designs. Selected designs contained between 5 (V2D9r) and 32 (V1D7) mutations. For PIP version 1, all selected designs expressed in the insoluble fraction after cell lysis and had to be purified from inclusion bodies. We selected one design to characterize in further detail based on an initial screen of catalytic activity (**Supplementary Table 10**), V1D8r. For version 2 we obtained one design that expressed in the soluble fraction, V2D9r.

### Structural characterization of designed KSI variants

We first compared the active site conformations of the computational models for the two selected designs, V1D8r and V2D9r, to that of the active site conformation of the wild-type protein. We defined two regions: the entire reshaped region, which consists of residues 34-45 for V1D8r and 34-46 for V2D9r, and a highly variable segment consisting of residues 37-42 for V1D8r and 37-43 for V2D9r. Both computational design models had conformations considerably different from the wild-type backbone (**Fig.4a**): V1D8r differed from wild-type by 2.41 Å backbone RMSD in the whole reshaped region and 3.49 in the highly variable region. V2D9r differed from wild-type by 2.50 Å backbone RMSD for the whole reshaped region and 3.34 Å RMSD for the highly variable region.

**Figure 4.**
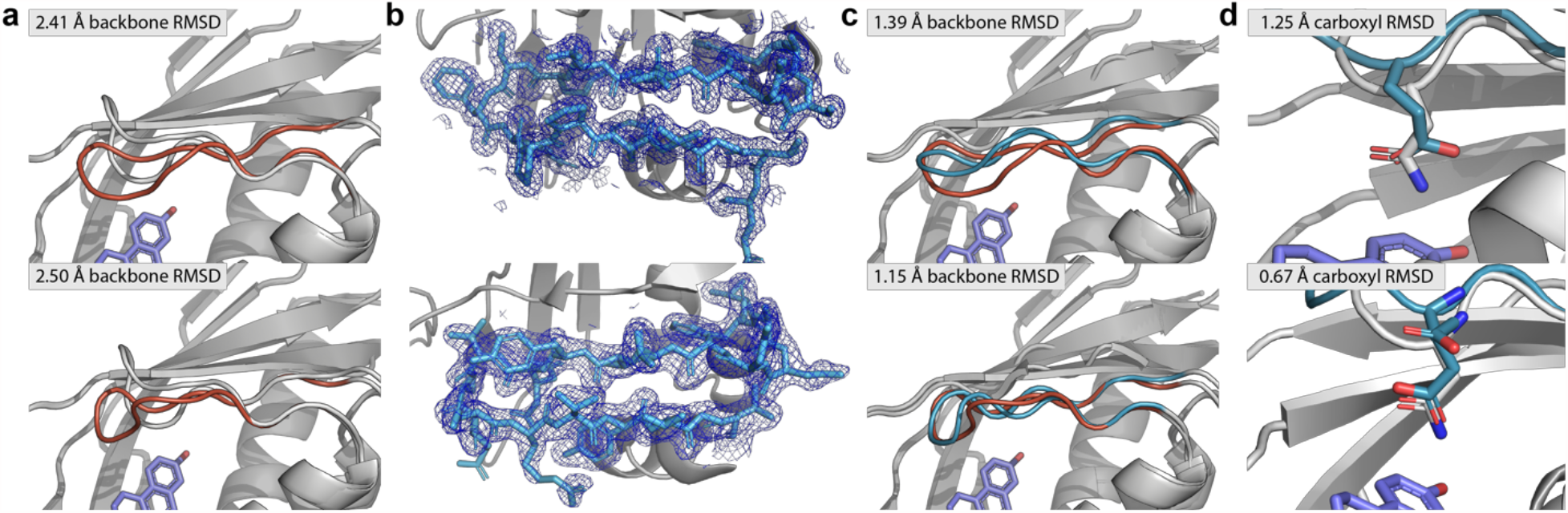
Structural characterization of designs V1D8r and V2D9r. **(a)** Overlay of wildtype KSI crystal structure (grey) and V1D8r (orange, top) and V2D9r (orange, bottom) lowest-energy design models. **(b)** Electron density of catalytic loop for V1D8r (top) and V2D9r (bottom) at 1.0 sigma in mesh representation. **(c)** Overlay of crystal structure (blue) for V1D8r (top) and V2D9r (bottom) with their lowest-energy design models (orange). **(d)** Crystal structure (blue) of V1D8r (top) and V2D9r (bottom) showing the catalytic glutamate’s placement relative to the amide in the KSI starting structure (PDB 1QJG) used to define the catalytic position (grey). RMSD values between compared structures are indicated in different panels.

To assess whether V1D8r and V2D9r indeed adopted the designed new backbone conformations, we determined crystal structures of the two designs. Both structures contained a ligand in the active site. For V1D8r, we observed density from deoxycholate retained from the purification process. V2D9r was co-crystallized with equilenin (which was present in the structure that was used as a basis for design) but also contained some residual density for deoxycholate (see further below). For both designs the electron density of the reshaped backbone region was well-resolved (**Fig. 4b**). Importantly, the backbone geometries of the reshaped backbone region in V1D8r and V2D9r were within 1.39 and 1.15 Å RMSD (N, C, Cɑ, and O backbone atoms) of the corresponding lowest-energy design models (**Fig.4c**).

We next examined the positioning of the catalytic glutamate carboxyl group. Each design placed the catalytic carboxyl within 1.3 Å RMSD of the wild-type aspartate carboxyl (**Fig. 4d**). V2D9r, which was co-crystallized with equilenin, placed the catalytic oxygen with particularly good accuracy (0.7 Å). We note that the crystal structures of both designs showed at least partial occupancy of deoxycholate in the ligand-binding site, and that it is conceivable that the bulky carboxyl moiety of the ligand changed the placement of the catalytic carboxyl group. This hypothesis is supported by the observation that the V2D9r crystal had partial occupancy of deoxycholate in 3 out of 4 asymmetric units, and the positioning of the catalytic residue in those monomers was significantly worse than in the asymmetric unit that contained only equilenin. In the asymmetric unit which contained only equilenin, we observed two distinct possibilities for the placement of the carboxyl of E38, which we modeled as alternate conformations (**Supplementary Fig. 6)**. Despite the apparent flexibility of E38, the crystallographic data support the conclusion that the designed backbone is indeed capable of supporting the desired functional site geometry, as one of the alternate conformations is close to the wild-type carboxyl placement (0.75 Å restraint satisfaction, 0.67 Å carboxyl heavy-atom RMSD compared to the amide group of 38N of 1QJG, **Fig. 4d**). Taken together, the structural analysis shows that PIP is capable of designing novel backbone conformations that differ by over 3 Å from their native counterparts with high accuracy.

### Functional characterization of designed KSI variants

Both designs V1D8r and V2D9r showed robustly measurable enzymatic activity when using 5(10)-estrene-3,17-dione as a substrate (**Fig. 5a,b**). Even though our structural analysis confirmed successful design of backbone and side chain placements, both designs were considerably less active than both wild-type and D38E KSI (**Table 1**): V1D8r and V2D9r had k_cat_ values of 1.7 and 0.29 min^-1^, respectively, compared to a k_cat_ value of 3.4 min^-1^ for D38E in our assay. This result might be expected for several reasons: First, while we took steps to avoid mutations in residues known to be important for catalysis, we still made extensive changes (19 and 12 mutations in design V1D8r and V2D9r, respectively) in and around the active site, which could change the electrostatic environment as well as affect functional or non-productive dynamics that impact catalysis. Second, while wild-type KSI is a dimer, our designs (while modeled as a dimer) are monomeric at the concentrations of the enzyme assay (**Supplementary Fig. 7**) although dimeric in the crystal. These differences could affect functional group positioning in solution^30^. Third, even though the glutamate side chain placement in V2D9r was close to ideal, it was not perfect; even small perturbations towards nonproductive conformations can be significant to catalysis and the designed glutamate may only sample catalytic conformations a fraction of the time. Finally, there is evidence that the catalytic residue in the homologous *Pseudomonas putida* KSI accesses multiple specific productive conformations to enable its participation throughout the catalytic cycle^31^, a property which was not considered by our protocol. Despite these difficulties, our designed enzymes still enhanced the catalysis of their substrate by 4 to 5 orders of magnitude when compared to the water-catalyzed isomerization of the similar 5-androstene-3,17-dione^32^.

**Figure 5.**
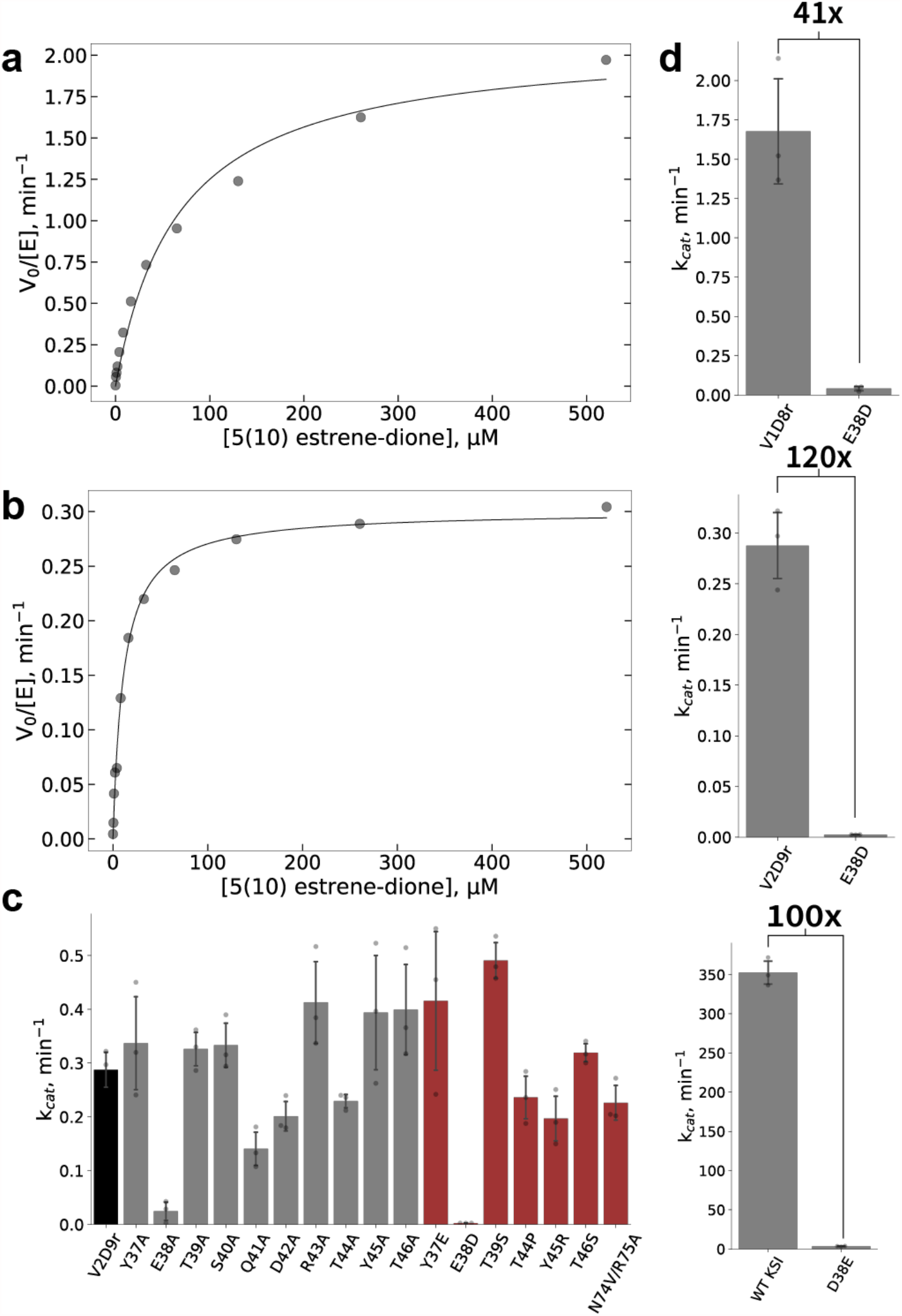
Functional characterization of designs V1D8r and V2D9r. Representative Michaelis-Menten curves for design V1D8r **(a)** or V2D9r **(b). (c)** Bar graph of *k*_cat_ values for design V2D9r (black), alanine scan mutants (grey), and reversion mutants (red). Standard deviation of independent triplicate experiments are shown as error bars with individual measurements shown as points. **(d)** Bar plots showing the k_cat_ values of V1D8r (top), V2D9r (middle), or wild-type KSI (bottom) and their E38D or D38E active site mutations. Values show the fold-change in k_cat_ between the respective D/E active-site residue pairs. Error bars and points are as in (c).

We chose the KSI model system because of its sensitivity to functional group positioning, specifically the considerable drop in catalytic activity upon addition of a carbon-carbon bond when mutating the catalytic aspartate to glutamate. To position the catalytic glutamate carboxylate in the designs close to the aspartate carboxylate in the wild-type, the crystal structures of both designs (**Fig. 4**) showed considerable remodeling of the KSI active site. Accordingly, we wondered whether subtracting the carbon-bond again, i.e. reverting the glutamate in the design back to the original wild-type aspartate, would have much reduced activity. Therefore, to determine whether the designs successfully switched KSI’s preference for its catalytic residue, we tested the catalytic activity of E38D reversion mutations in the context of the V1D8r and V2D9r designed backbones (both E38D reversion mutants in the design background were folded, **Supplementary Fig. 6b-d**). For both V1D8r and V2D9r, we found a substantial reduction in k_cat_ in the E38D reversion mutant; the activities of both mutants were near the detection limit of the assay, and were at least 41-fold and 119-fold for V1D8r and V2D9r, respectively. These fold changes are similar to the fold-change in k_cat_ between wild-type KSI and the D38E mutations (**Fig. 5c, Table 1**). This result suggests that the positioning of the catalytic residue’s carboxyl moiety is still an important determinant of catalytic activity despite the designs’ low k_cat_ compared to wild-type, and that the designed backbone geometries successfully altered the enzyme’s preference for its catalytic residue.

Finally, we sought to test the robustness of V2D9r’s redesigned backbone segment to mutation. To determine whether the activity of V2D9r was dependent on any particular residue in the redesigned region, we performed an experimental alanine scan along all mutated residues. We also made reversion mutants for residues whose backbone atoms did not move significantly between the wild-type and the design conformations (**Table 1, Fig. 5d**). No alanine or reversion mutant affected the k_cat_ more than two-fold except for the catalytic glutamate, suggesting that the designed loop depends on several interactions to adopt a catalytically competent conformation, as well as the glutamate as a general base.

## DISCUSSION

We introduced and validated methods to accurately position functional groups in protein active sites by computational design. We developed and benchmarked two new robotics-inspired sampling methods, FKIC and LHKIC (**Fig. 1**), which model the conformations of backbone segments with high accuracy and efficiency (**Fig. 2**). We then integrated these methods into a new design protocol, PIP (**Fig. 3**), which we validated experimentally by solving crystal structures of designs with reshaped active site regions (**Fig. 4**).

FKIC leads to considerable improvements over the two approaches it combines, the fragment-independent loop modeling method NGK^10^ and the fragment-insertion based prediction approach CCD^21^ (**Fig. 2a**). In addition to sub-Å structure prediction accuracy, our results demonstrate that FKIC provides up to ∼140-fold improvement in sampling native-like conformations on the challenging problems of modeling local protein conformations with multiple segments and arbitrary secondary structure composition. This key advance in sampling performance paves the way to integrate FKIC with other methods in a variety of applications. We expect that FKIC will be most useful in applications that predict or design the conformations of local regions in proteins where the remainder of the protein stays relatively fixed or undergoes only slight adjustments. As such, applications to homology modeling may require integration of FKIC with more aggressive remodeling in the entire protein, not just a local region (**Supplementary Note 4, Supplementary Table 11**). In contrast, FKIC should be suited to accurately predict the conformations of local protein regions from limited or low-resolution experimental data, and, in combination with LHKIC, to design new backbone geometries not seen in nature. Our results provide a first proof-of-concept for the latter. LHKIC, while performing similarly to FKIC overall when predicting conformations given their sequences (slightly worse in the median sub-A fraction on the standard dataset, **Supplementary Table 1a**), is well-suited to design because of the ability of LHKIC to sample both new sequences and new structures simultaneously.

Both versions of PIP used similar robotics-inspired approaches to conformational sampling, but PIP version 2 placed an additional emphasis on fragment-based sampling using FKIC / LHKIC, and analysis of fragment quality using Pareto fronts (**Fig. 3**). Fragment quality measures how well designs conform to local sequence / structure relationships observed in naturally occurring proteins, and designs with a better fragment quality might be expected to be more stable^33^.

Indeed, FKIC sampling improvements and fragment quality are correlated (**Fig. 2d**) and attention to fragment quality as a design metric may have resulted in several beneficial characteristics in design V2D9r, which was both more soluble when expressed in *E. coli*, and had a higher T_M_ (**Supplementary Fig. 6**) than design V1D8r. However, our design sample is small and further exploration of the impact of fragment-based design on design success would be interesting.

Despite the success with positioning a functional group that required reshaping of an active site backbone region, our results also highlight the considerable challenges faced when designing functional proteins. There are several main problems. First, the irregular geometries of naturally occurring proteins are more difficult to engineer than entirely *de novo* protein structures that largely consist of regular elements of secondary structure connected by very short loops^34^. One solution would be to start with entirely *de novo* designed proteins into which to build new functions, but for certain applications and functions, redesign of existing proteins that already possess desired properties may be the preferred strategy. In those cases, the methods presented here provide a new approach. Moreover, FKIC/LHKIC and related methods could also provide a new way to systematically reshape local regions to endow *de novo* designed proteins with new functions. The second problem concerns the difficulty of predicting the conformations and energetics of polar interactions making up protein functional sites with sufficient accuracy. Third, in the case of enzyme engineering, current computational design methods may not correctly represent and optimize for important determinants of specific catalytic functions, such as the role of dynamics or active site electrostatics. Moreover, these determinants may be incompletely understood. Fourth, certain functions may require switching between two or more approximately isoenergetic conformations. Such a scenario is much more challenging to engineer than optimizing for one deep energy minimum, which is sufficient for successful *de novo* design of protein structures. Nevertheless, the ability to sample both conformational and sequence space afforded by the robotics-inspired approaches and protocols presented here should help address these problems, and be useful in both the design and modeling of novel loop conformations that enable specific functional geometries that do not yet exist in nature.

## METHODS

### Structure prediction simulations

#### Fragment-sampled KIC (FKIC) Overview

We developed FKIC as a new protocol for modeling local protein geometries in Rosetta. FKIC is based on the KIC protocol^9^, but, instead of sampling non-pivot φ/ψ torsions probabilistically from Ramachandran space, FKIC uses coupled φ/ψ/ω degrees of freedom from consecutive residues of protein fragments of size nine, three or one to sample conformational space. During the low- and high-resolution sampling stages (**Fig. 1b**), each KIC move in the original KIC protocol is replaced by an FKIC move (**Fig. 1a**). An FKIC move consists of the following sequence of steps: (i) a fragment library (see “Generation of fragment libraries” section below) is chosen at random from all available libraries (i.e. 9mers, 3mers and 1mers), (ii) the chosen fragment library is searched for fragment alignment frames that (at least partially) overlap with the given target sub-segment, (iii) one of the alignment frames is chosen at random, (iv) one of the 200 fragments contained in the given alignment frame is chosen at random, (v) the φ/ψ/ω torsions of that fragment are applied to the respective overlapping region of the given target sub-segment, and (vi) the segment is closed using kinematic closure. We ran the FKIC protocol using the same fragment libraries as for the CCD protocol.

#### Loophash-sampled KIC (LHKIC) Overview

We developed LHKIC for designing local protein geometries in Rosetta. LHKIC and FKIC share the same simulation protocol (**Fig. 1, Supplementary Fig. 1**). In LHKIC, the non-pivot φ/ψ/ω degrees of freedom are sampled from fragments picked by the loophash algorithm^16^. At each KIC sampling step, we calculate the 6D transformation from the residue before the first pivot to the residue after the last pivot. We use the 6D transformation to query a pre-compiled loophash database (see “Generation of loophash databases” in **Supplementary Methods**). One 6D transformation query can return multiple loops. Torsions of a random loop from the returned loops are applied to the residues between the pivot residues. By default, LHKIC does not mutate the sequence of the loop. When the loophash_perturb_sequence option is set to true, LHKIC applies the sequence of the returned loop to the pivot residues and the residues between the pivots.

#### Rosetta Simulations

FKIC and NGK benchmarking simulations were performed using the Rosetta macromolecular modeling and design suite (https://www.rosettacommons.org/software), revision 59052. The LHKIC method was developed later and used Rosetta revision 60022. KIC simulation results reported in **Fig. 2a** in the main text were taken from ref.^10^. Since the publication of the original KIC method, the Rosetta energy function has undergone several revisions, including the changes described in the “talaris2013” and “talaris2014” versions^35^ and the latest improvements made in the “ref2015” version^17^. The ref2015 energy function^17^ was used for all benchmarks unless otherwise noted. Compared to the other energy functions, ref2015 showed a consistent performance improvement (**Supplementary Table 12**).

#### Benchmark Datasets

##### 12-Residue “*Standard*” benchmark dataset

We first tested FKIC on the 12-residue loop benchmark dataset used in previous work^9,10,36^. This benchmark dataset consists of 45 protein structures from the Protein Data Bank (PDB) containing non-redundant 12-residue target segments without regular secondary structure, curated from two previously described datasets^19,37^. We used this dataset even though it is not ideal (for example, the conformation of several segments might be influenced by crystal contacts, see **Supplementary Note 1** and **Supplementary Table 5**) to facilitate comparison of FKIC with previous protocols. For each loop, we retained the N and Cα atoms of the N-terminal residue, as well as the Cα, C and O atoms of the C-terminal residue, which serve as loop anchor points for kinematic closure (as in ref.^9^).

##### 16-Residue “Mixed Segment” benchmark dataset

The *Mixed Segment* benchmark dataset consists of 30 structures from the PDB containing 16-residue target segments. The target segments were derived from structures in the *Standard* benchmark dataset above using the following criteria:

- The crystallographic resolution of the experimentally determined structure is equal to or better than 2Å.
- Each segment has 5 to 11 residues that contain alpha helices or beta strands defined using DSSP^38^ and the remainder of the segment is designated as loop.
- Residues in the segment are at least 4Å away from any chains or copies of the molecule in other asymmetric units, to avoid crystal contacts.
- The segment is at least 5 residues away from the chain termini.

The segment that satisfied all criteria with the lowest distance from the protein surface was selected.

##### 10-Residue “*Multiple Segments*” benchmark dataset

The *Multiple Segments* benchmark dataset consists of 30 structures from the PDB each containing a pair of 10-residue interacting target segments. Structures were derived from the top8000 dataset^39^ using the following criteria:

- The crystallographic resolution of the experimentally determined structure is equal to or better than 2Å.
- Each segment has less than 3 residues that have regular secondary structure, defined as above.
- Residues in the segment are at least 4Å away from any chains or copies of the molecule in other asymmetric units, to avoid crystal contacts.
- Each segment is at least 5 residues away from the chain termini.
- Segments in each pair are within 4Å and are separated by at least 5 residues in primary sequence.

The pair of segments that satisfied all criteria with the lowest distance from the protein surface was selected. We also constructed two analogous sets that contain either two 8-residue segments or two 12-residue segments (**Supplementary Note 3; Supplementary Table 6** and **Supplementary Table 7**).

#### Preparation of benchmark input structures

To exclude information on the native conformation of the target segment(s) for all benchmark datasets, all side chains in the segment(s) as well as side chains within 10Å of the segment(s) (based on all-atom pairwise distance measurements) were removed. The backbone information was removed by changing the segment into an extended conformation with idealized bond lengths and angles. The datasets were constructed with an openly available script: https://github.com/Kortemme-Lab/benchmark_set_construct.

#### Generation of fragment libraries

We generated libraries of 9-mer and 3-mer fragments for all benchmark cases using the fragment picking method described in ref.^15^. The method selects fragments from a representative database of 16,801 protein chains extracted from the PDB and culled such that any two chains have at most 60% sequence identity. The fragment database is part of the Rosetta software and located in rosetta/tools/fragment_tools/vall.jul19.2011.gz. The fragments selected for each segment sequence position span a 3- or 9-residue frame, which overlaps with neighboring frames. Moreover, we allow sampling of 1-mer fragments, which consist of a single triplet of φ/ψ/ω torsions that are generated on the fly by the respective modeling protocol (see below) based on the 3-mer fragment library for the given position. The make_fragments.pl script in the rosetta/tools/fragment_tools/ directory integrates several data sources to maximize fragment quality, including sequence similarity and the detection of homologs using PsiBLAST^40^, predicted secondary structure similarity using PsiPred^41^ and prediction of preferred φ/ψ torsions and solvent accessibility using SPARKS-X^42^. Importantly, for benchmarking purposes, we ran simulations using fragment libraries that excluded homologs to the given query sequence, by providing the –nohoms flag to the make_fragments.pl script, which excluded all protein chains with a PsiBLAST E-value < 0.05^40^ from the fragment picking process.

#### Simulation protocols

The Rosetta ‘CCD’ loop modeling method using fragment insertion and the cyclic coordinate descent (CCD) closure technique^21^ is described in ref.^35^. The NGK loop modeling method is described in ref.^10^. For FKIC simulations, NGK was modified to sample torsions from the generated fragment libraries described above. Similarly, for LHK, NGK was modified to sample torsions from loops picked using loophash^43^ (**Supplementary Methods**). For control simulations that use native bond lengths and angles as input, we replaced the input structure with the native structure and disabled the randomization of torsions at the beginning of the simulation. Full descriptions of RosettaScripts code and command lines can be found in **Supplementary Methods**.

#### Fragment distance calculation

The chord distance^44^ was calculated between pairs of fragments. The chord distance between two angles is defined as: *D*^2^(*θ*_1_,*θ*_2_)=2−2cos(*θ* _2_−*θ*_1_)In our case, this value was calculated for backbone dihedral angles and summed over paired residues between fragments and target loops 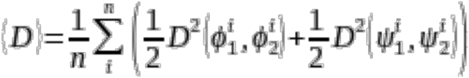 with n=3 defining 3mer fragments for example. (*D*) will have a minimum of 0 if the angles match exactly and a maximum of 4 if the angles differ by 180 degrees.

### Pull Into Place (PIP) design protocol

#### Rosetta version

PIP was run using Rosetta commit 10b6f2f8e20d70757e6b510def2ddcbeef172538 (PIP version 1) or revision 60048 (PIP version 2). We used the latest available scorefunction for each PIP version, which were talaris2013 for PIP version 1 or ref2015 for PIP version 2.

#### Input files

KSI designs were based on PDB structure 1QJG^45^. KSI is an obligate dimer, so we included both monomers in our initial structure. To design different loop lengths, we created several versions of the initial structure: one for each deletion of up to 6 residues, and one with wild-type length. We replaced N38 with a glutamate and relaxed the resulting models 100 times in the talaris2013 (version 1) or ref2015 (version 2) score function using FastRelax, with all atom coordinates restrained to their starting positions. The best (lowest-scoring) relaxed structure was then repacked 100 times, and the lowest-scoring model was used as a template for design. The desired position of the E38 sidechain was defined in a restraint file (**Supplementary Methods**). Each atom in the E38 carboxylate group was restrained to the position of the corresponding atom in the N38 amide group in the starting structure (1QJG). We visually confirmed that N38 in 1QJG had the same rotameric conformation as D38 in wild-type KSI (PDB structure 8CHO). When working with the version 1 designs, we noticed that F54 sometimes changed its rotamer conformation, causing subsequent designed mutations to stabilize the altered conformation. For version 2, we addressed this issue by placing restraints on the zeta-carbon of F54 in a manner similar to the catalytic residue (**Supplementary Methods**). The residues being remodeled were defined in loop files (**Supplementary Methods**). We chose which residues to remodel based on proximity to secondary structure elements and intuition. Our goals were (i) to allow sufficient remodeling on either side of E38 to stabilize its new conformation, (ii) to anchor the loop in secondary structural elements, and (iii) to minimize loop length. With these considerations in mind, we remodeled loops that were 7-13 (version 1) or 12-13 (version 2) residues long. For version 2, we also remodeled a second 4-residue loop on the dimer interface (residues 199-202 using Rosetta numbering, residues 74-77 on chain B using PDB numbering), hoping to maintain favorable contacts between those residues and the catalytic loop.

The residues that were allowed to design (change amino acid identity) and repack (only change rotamer conformation) were specified in a resfile (**Supplementary Methods**). For version 1, any residue that had a sidechain atom within 4Å or 6Å of any loop atom in any model generated in PIP step 1 was allowed to design or repack, respectively. For version 2, we only allowed Rosetta to design residues on the catalytic loop, as well as four residues on the short dimerization loop which directly interacts with the catalytic loop (described above) (**Supplementary Methods**). Repackable residues were selected using the Rosetta clash-based repack shell selector. F54, A114, and F116 were not allowed to design in either version because they are known to be important for positioning the catalytic residue^46^. For version 1, each designed residue was allowed to change to any of the 20 canonical amino acids except cysteine (due to the potential for disulfide bonds) and histidine (due to the potential for pH-dependent behavior). For version 2, we used the LayerDesign task operation in Rosetta to determine which residue identities were allowed at each position. Since version 2 introduced backbone degrees of freedom during the design step, we specified a fold tree to keep conformational changes as local as possible (**Supplementary Methods**).

#### PIP Step 1: Build Models

We created models positioning the E38 carboxylate group by running 20,000 NGK^10^ (version 1) or LHKIC (version 2) simulations with restraints as described above. Backbone remodeling was limited to the loop defined in the appropriate loop file and design was allowed according to the appropriate resfile (**Supplementary Methods**). In version 1 of PIP, the initial coordinates of the loop being remodeled were discarded and rebuilt from scratch. This step was skipped for version 2. Only models that put all three restrained atoms within 0.6 Å (version 1) or 0.7 Å (version 2) of their intended positions were carried on to the next step.

#### PIP Step 2: Design Models

To stabilize models that correctly positioned E38, we ran 50 fixed-backbone (version 1) or 10 FastDesign (version 2) simulations per model, or more if there were relatively few models. Design was allowed according to the appropriate resfile (**Supplementary Methods**). For version 1, we picked 200 designs for PIP step 3 with probability proportional to their Boltzmann-weighted talaris2013 scores (in Rosetta energy units, REU). For version 2, we used Pareto fronts to pick designs so that we could supplement information from the Rosetta score function with additional metrics. These metrics consisted of the total solvent-accessible surface area of the model, two “foldability” metrics that perform 60 brief forward-folding simulations on pieces of the loop and report the fraction of results that placed the segment’s N-terminus within 4 Å of the concomitant residue in the design structure, the maximum distance of the restrained atoms to their ideal position, a fragment quality metric, and the Rosetta fa_attr score. The Foldability metrics remove a portion of the design’s backbone, then rebuild it starting from the N-terminus of the deleted segment using fragment-based assembly. This is repeated 100 times, and the average distance of the C-terminus of the rebuilt segment to its position in the design is reported. The fragment quality metric assesses the geometric similarity between 9-residue fragments in the designed loops and fragments of natural proteins in the PDB, as described in ref.^33^. Specifically, we picked protein fragments from the PDB based on their similarity in sequence, predicted solvent exposure, and predicted secondary structure to the post-simulation design sequence and determined the RMSD of the backbone atoms in each fragment to the final structure. We then determined the lowest RMSD at each position being evaluated, and reported the highest of these.

#### PIP Step 3: Structure Prediction

We computationally assessed our designs by running between 100 and 500 NGK (version 1) or FKIC (version 2) structure prediction simulations for each design; for version 2, we opted to perform fewer structure prediction simulations on a larger number of designs during the early rounds of design. Backbone movement was limited to the loop defined in the appropriate loop file (**Supplementary Methods**). The initial coordinates for that loop were discarded and rebuilt from scratch. Any design for which the lowest scoring decoy put all three carboxylate atoms within 1.2 Å of their intended positions was carried on to the design selection step. Furthermore, any decoy (regardless of score) that put all three carboxylate atoms within 0.6 Å of their intended positions was used as input for a second round of design simulations.

#### PIP Shared Parameters

Filter, scorefunction, residue selector, and certain Rosetta mover definitions were used during every step of the PIP protocol. These were stored as separate RosettaScripts files and imported into each main step template. The fragment quality filter additionally required a weights file describing which scores to use when picking fragments^15^ (**Supplementary Methods**).

#### Design Selection

We picked designs to experimentally test by comparing quality metrics and visually inspecting models. The quality metrics are described in **Supplementary Table 8**. We paid particular attention to the score gap, which measures the difference in the score between the lowest-scoring model with under 1 Å restraint satisfaction and the lowest-scoring model with over 2 Å restraint satisfaction. Several designs were selected despite having a score gap of 0 REU, as they had multiple low-energy conformations. We also made an effort to pick designs from different sequence and structure clusters. Design sequences were clustered hierarchically such that inter-cluster distance was no greater than the mean sequence distance (calculated according to the BLOSUM80 substitution matrix) across all designs. Structure clusters were formed hierarchically such that the RMSD between any two designs in the same cluster was no greater than 1.2 Å. We visually inspected the lowest scoring model for each design to eliminate those with irregular backbone or strained sidechain conformations.

#### Wildtype Reversions

For each design from PIP version 1 selected for experimental validation, we reran the structure prediction simulations (PIP Step 3) for each single wildtype reversion mutation. We then combined any reversions that had no apparent detrimental effect on our quality metrics and again ran the structure prediction simulations. In cases where the combination of all the individually acceptable reversions had a deleterious effect, we selected more conservative combinations of reversions for additional structure prediction simulations. If no acceptable combination of reversions could be found, no reversions were made (**Supplementary Table 8**). For PIP version 2, positions where the backbone was in a similar position to wildtype were reverted, or in some cases mutated to a residue picked by visual inspection (**Supplementary Table 9**), and designs with and without those reversions were ordered for experimental validation.

### Experimental characterization

#### Cloning and purification

The 14 designs chosen for experimental tests from PIP version 1 were ordered from GenScript pre-cloned into the pET-21a expression vector. For PIP version 2, and for characterization of V1D8r and the wild-type protein, we used an expression vector using parts from the modular yeast cloning toolkit^47^ which was similar to pET-21a, except that the cloning resulted in a glycine-serine genetic scar at the C-terminus. Full sequences of ordered designs and vectors can be found in **Supplementary Data 1**. Proteins were expressed in *E. coli* BL21(DE3) cells. Wild-type KSI and design V2D9r were purified essentially as described previously^48,49^ with minor differences. Briefly, cells were lysed in 40 mM potassium phosphate, 2 mM DTT, 1 mM EDTA, and 6 U/mL DNAse I, pH 7.2 using a Microfluidics M-110L microfluidizer. Clarified lysate was then passed through a 10 mL sodium deoxycholate gravity affinity column, prepared as described in reference^48^. The column was washed with 400 mM phosphate, 2 mM DTT, 1 mM EDTA, pH 7.2 followed by lysis buffer (minus DNAse), then eluted with 40 mM phosphate, 2 mM DTT, 1 mM EDTA, and 50% ethanol, pH 7.2. Proteins were then either further purified using a HiLoad 16/600 Superdex 75 pg gel filtration column or dialyzed twice in 1L lysis buffer to remove the ethanol. Most other designed proteins expressed in the insoluble fraction, so inclusion bodies were first purified from the cell lysate: Cells were grown in 1 L LB broth to an optical density of 0.6 at 37 C, followed by overnight expression at 18 C. Cells were then harvested by centrifugation at 3500 rpm for 20 minutes at 4 C, then resuspended in lysis buffer (40 mM Tris-HCl, 1 mM EDTA, 25% sucrose w/v, pH 8.5). Suspensions were lysed in a M-110L microfluidizer and centrifuged at 20,000 rpm for 20 minutes at 4 C. The resulting inclusion body pellet was washed once in 25 mL of 20 mM Tris-HCl, 1% sodium deoxycholate, 200 mM NaCl, and 2 mM EGTA, followed by at least 3 washes of 25 mL 10 mM Tris-HCl, 0.25% sodium doxycholate, pH 8.5, followed by at least 3 washes of 25 mL 20 mM Na-HEPES, 500 mM NaCl, 1 mM EDTA, pH 8.5. Inclusion bodies were centrifuged at 8,000 xg for 10 minutes at 4 C between washes. Proteins in inclusion bodies were solubilized by shaking for 30 minutes with 10 mL 8 M urea, 20 mM Na-HEPES, 500 mM NaCl, 10 mM DTT, and 1 mM EDTA at pH 8.5, then centrifuged at 20,000 rpm for 20 minutes at 4 C to remove cell debris. Solubilized protein was then refolded by stirring for 2 hours at 4 C in 200 mL of 40 mM KPi, 1 mM EDTA, 2 mM DTT. Proteins were then sterile-filtered using 0.4 μm filter paper and further purified via deoxycholate column as described above.

#### Activity assay

Purified KSI variants were tested for catalytic activity using an absorbance assay. 5(10)-Estrene-3,17-dione was solubilized at 2.1 mM and serial-diluted two-fold down to 0.51 μM in 100% DMSO. 115 μL enzyme, prepared in 40 mM potassium phosphate, 2 mM DTT, and 1 mM EDTA at pH 7.2, was then added to 5 μL of substrate for final substrate concentrations between 520 and 0.51 μM. K_M_ and k_cat_ values for the WT enzyme and designs V1D8r and V2D9r were measured at enzyme concentrations between 0.5 and 18 μM. For reversion and alanine scan mutations, k_cat_ values were measured in triplicate at 512 µM substrate. Absorbance at 248 nm was tracked for 5 minutes in a Varian Cary 50 Bio UV-Visible spectrophotometer using a 1cm path length. The first 30-60s of each reaction were excluded to allow the reaction to reach steady-state.

#### X-ray crystallography

Designed proteins were crystallized in 1 M ammonium sulfate (design V1D8r) or 1.6 M ammonium sulfate, 50 mM potassium phosphate, pH 7.2 (design V2D9r) using the hanging drop method. For design V2D9r, an equal volume of 2 mM equilenin (CAS 517-09-9 from Steraloids Inc., catalog ID E0400-000) was added to each drop.

#### X-ray data collection and processing

Prior to X-ray data collection, crystals were cryoprotected and flash-cooled by rapid plunging into liquid nitrogen. Crystals that yielded the V1D8r structure were cryoprotected using a mixture of 50% glycerol and 50% crystallization mother liquor, and crystals that yielded the V2D9r structure were cryoprotected using a mixture of 25% glycerol and 75% crystallization mother liquor. We collected single-crystal X-ray diffraction data on beamline 8.3.1 at the Advanced Light Source. Data collection for V1D8r was performed while the beamline was equipped with a Quantum 315r CCD detector (ADSC), while data collection for the V2D9r structure utilized a newer Pilatus3 S 6M photon-counting detector (Dectris). Both data sets were collected using an X-ray energy of 11111 keV, and the crystals were maintained at a cryogenic temperature (100 K) throughout the course of data collection.

We processed the X-ray data using the Xia2 system^50^, which performed indexing, integration, and scaling with XDS and XSCALE^51^, followed by merging with Pointless^52^. For the [6UAE] structure, a resolution cutoff (1.93 Å) was taken where the signal-to-noise ratio (<I/σI>) of the data fell to a value of approximately 1.0. In the case of the V1D8r structure, the data were collected on an older, smaller detector, and the resolution was limited by the detector edge and the geometric requirements of the experiment. Although other metrics of data quality (such as CC1/2 and <I/σI>) suggest that a more aggressive resolution cutoff would be acceptable, we were limited by the data completeness that could be obtained with the minimum accessible sample-to-detector distance. Further information regarding data collection and processing is presented in **Supplementary Table 12**. The reduced diffraction data were analyzed with phenix.xtriage to check for common crystal pathologies, none of which were identified.

#### Structure determination

We obtained initial phase information for calculation of electron density maps by molecular replacement using the program Phaser^53^, as implemented in the PHENIX suite^54^. For the V1D8r structure, we identified a single copy of the protein in the asymmetric unit using the coordinates from a previous KSI model, and for the V2D9r structure we identified four copies of the protein in the asymmetric unit. Both solutions were consistent with an analysis of Matthews probabilities for the observed unit cell and molecular weight of the protein^55,56^.

We manually rebuilt the molecular replacement solutions using the resulting electron-density maps, followed by iterative refinement of atomic positions, individual atomic displacement parameters (B-factors) with a TLS model, and occupancies, using riding hydrogen atoms and automatic weight optimization, until the model reached convergence. Throughout the course of manual model building, electron density corresponding to several ligand molecules became apparent, which we were able to model. In the V1D8r structure, we observed electron density for two steroid-like molecules, one occupying the KSI active site, and a second nestled at a crystal contact. These densities were modeled using deoxycholate, which was present in one of the purification buffers used to prepare the crystallization samples. Additionally, we identified two phosphate ions in this structure. In the V2D9r structure, we also saw density for steroid ligands in the active sites of each of the four copies of the enzyme. In this case, the modeling was challenging, because the samples were exposed to both deoxycholate (during purification) and equilenin (post-purification), and electron density features suggested that there could be a mixture of both ligands represented in the electron density. We attempted to model various combinations of the ligands into the active site densities, and found that the electron density features could best be described by modeling equilenin in one active site (chain B), deoxycholate in one active site (chain D), and a mixture of both ligands in the other two active sites (chains A and C). Our choice to model the ligand densities in this way is based on both reduction of refinement R-factors, as well as on overall flatness of residual Fo-Fc difference density maps around the modeled ligands. In the V2D9r structure, we also modeled 12 sulfate ions. All model building was performed using Coot^57^ and refinement steps were performed with phenix.refine within the PHENIX suite^54,57^. Restraints for the ligands were calculated using phenix.elbow^58^. The final model coordinates were deposited in the Protein Data Bank (PDB^59^) under accession codes 6UAD and 6UAE. Further information regarding model building and refinement is presented in **Supplementary Table 12**.

#### Size exclusion chromatography

Six uM of purified wild-type KSI, V1D8r, or V2D9r were loaded onto a Superdex 75 10/300 GL column from Cytiva which was pre-equilibrated with running buffer (40 mM phosphate, 2 mM DTT, and 1 mM EDTA). Samples were run isocratically in an Agilent Technologies 1200 Series HPLC for 150 minutes and absorbance was monitored at 280 nm.

#### CD spectroscopy

Samples for CD analysis were prepared at approximately 6 μM enzyme in 40 mM phosphate pH 8.5, 2 mM DTT, and 1 mM EDTA. CD spectra were recorded at 25°C using 2 mm cuvettes (Starna, 21-Q-2) in a JASCO J-710 CD spectrometer (Serial #9079119). The bandwidth was 2 nm, rate of scanning 20 nm/min, data pitch 0.2 nm, and response time 8 s. Each spectrum represents the average of 5 scans. Buffer spectra were subtracted from the sample spectra using the Spectra Manager software Version 1.53.01 from JASCO Corporation. Melting temperatures were assessed by measuring molar ellipticity at 222 nm and increasing the temperature from 25 C to 95 C at 1°C per minute, using a data pitch of 0.5°C.

#### RMSD Calculations

For all RMSD calculations, structures were aligned to all residues except those involved in the RMSD calculation. In order to calculate backbone RMSDs that involved comparing the shorter V1D8r loop to the full-length WT protein, we had to exclude one residue in WT structure. We chose to exclude residue 38, as this resulted in the lowest RMSD between the design and the WT protein.

## Supporting information

Supplementary Sequences

## Data Availability Statement

Coordinates and structure files have been deposited to the Protein Data Bank (PDB) with accession codes 6UAD (V1D8r) and 6UAE (V2D9r). All other relevant data are available in the main text or the supplementary materials.

## Code Availability Statement

Rosetta source code is available from rosettacommons.org. Pull Into Place is available at https://github.com/Kortemme-Lab/pull_into_place. The parameter files used to design KSI are available at https://github.com/Kortemme-Lab/ksi_inputs.

## ACKNOWLEDGEMENTS

The authors would like to thank Dan Herschlag for discussions on KSI that in part inspired the project and for comments on the manuscript, Marco Mravic, Gabe Reder, and Yessica Gómez for contributions to testing robotics-inspired sampling methods, Shivani Mathur for experimental help characterizing expression of designs, the Dueber lab for providing a custom expression vector, and the Kortemme group for discussions. The work was supported by National Science Foundation (NSF) grant DBI-1564692 and National Institutes of Health (NIH) grant R01-GM110089 to TK; and NSF grant MCB 1714915 and NIH grant GM123159 to JSF. We additionally acknowledge the following fellowships: UCSF Discovery Fellowship (XP) and NIH F32 Postdoctoral Fellowship (MCT). TK is a Chan Zuckerberg Biohub Investigator.

## AUTHOR CONTRIBUTIONS

CK, KK, XP, RAP, and TK developed the concepts for the project. KK and CK developed and applied the PIP approach and performed the majority of the experiments, with contributions from LL. RAP and XP developed and benchmarked the FKIC and LHKIC methods, with contributions from SOC, JJ, and JJG. CK, MT, LL, and JSF determined the crystal structures. JJG, JSF, and TK provided guidance, mentorship and resources. CK, KK, XP, RAP, and TK wrote the manuscript with contributions from the other authors.

