## Supplementary Materials for "Accurately positioning functional residues with robotics-inspired computational protein design"

### Title:

**Supplementary Methods**
**Supplementary Notes 1-4**
**Supplementary Tables 1 – 13**
**Supplementary Figures 1 – 7**
**References Cited in the Supplementary Materials**

### SUPPLEMENTARY METHODS

#### Rosetta scripts & command line for CCD

We used the following Rosetta scripts to run the CCD benchmark simulations:

```
<ROSETTASCRIPTS>
   <TASKOPERATIONS>
      <RestrictToLoops name="loop" loops_file="%%loop_file%%"/>
   </TASKOPERATIONS>
   <MOVERS>
      <LoopmodelWrapper name="modeler" loops_file="%%loop_file%%" fast="%%fast%%"/>
   </MOVERS>
   <PROTOCOLS>
      <Add mover_name=”modeler”/>
   </PROTOCOLS>
</ROSETTASCRIPTS>
```

Let the home directory of Rosetta be */path/to/rosetta/main*; the input structure be *my_structure.pdb*; the loop file be *my_structure.loop*; the fragment files be *my_structure.200.9mers.gz* and *my_structure.200.3mers.gz*; and the Rosetta script XML file be *loopmodel.xml*. Then the command to run one simulation is:

*/path/to/rosetta/main/source/bin/rosetta_scripts.linuxgccrelease -database /path/to/rosetta/main/database -in:file:s my_structure.pdb -parser:protocol loopmodel.xml -parser:script_vars loop_file=my_structure.loop fast=no -out:prefix prefix -overwrite -loops:remodel quick_ccd -loops:refine refine_ccd -ex1 -ex2 -loops:frag_sizes 9 3 1 -loops:frag_files my_structure.200.9mers.gz my_structure.200.3mers.gz*

#### Rosetta scripts & command line for NGK

We used the following Rosetta script to run the NGK benchmark simulations:

```
<ROSETTASCRIPTS>
   <TASKOPERATIONS>
      <RestrictToLoops name="loop" loops_file="%%loop_file%%"/>
   </TASKOPERATIONS>
   <MOVERS>
      <LoopModeler name="modeler" config="kic" loops_file="%%loop_file%%" fast="%%fast%%" />
   </MOVERS>
   <PROTOCOLS>
      <Add mover_name=”modeler”/>
   </PROTOCOLS>
```

```
</ROSETTASCRIPTS>
```

Let the home directory of Rosetta be */path/to/rosetta/main*; the input structure be *my_structure.pdb*; the loop file be *my_structure.loop*; and the Rosetta script XML file be *loopmodel.xml.* Then the command line to run one simulation is:

*/path/to/rosetta/main/source/bin/rosetta_scripts.mysql.linuxgccrelease -database /path/to/rosetta/main/main/database -in:file:s my_structure.pdb -parser:protocol loopmodel.xml -parser:script_vars loop_file=my_structure.loop fast=no -out:prefix prefix -overwrite*

**Rosetta scripts & command line for FKIC**

We used the following Rosetta script to run FKIC benchmark simulations:

```
<ROSETTASCRIPTS>
  <TASKOPERATIONS>
    <RestrictToLoops name="loop" loops_file="%%loop_file%%"/>
  </TASKOPERATIONS>
  <MOVERS>
    <LoopModeler
      name="modeler"
      config="kic_with_frags"
      loops_file="%%loop_file%%"
      fast="%%fast%%">
    </LoopModeler>
  </MOVERS>
  <PROTOCOLS>
    <Add mover_name=”modeler”/>
  </PROTOCOLS>
</ROSETTASCRIPTS>
```

Let the home directory of Rosetta be */path/to/rosetta/main*; the input structure be *my_structure.pdb*; the loop file be *my_structure.loop*; the fragment files be *my_structure.200.9mers.gz* and *my_structure.200.3mers.gz*; and the Rosetta script XML file be *loopmodel.xml.* Then the command line for one simulation is:

*/path/to/rosetta/main/source/bin/rosetta_scripts.mysql.linuxgccrelease -database /path/to/rosetta/main/database -in:file:s my_structure.pdb -parser:protocol loopmodel.xml -parser:script_vars loop_file=my_structure.loop fast=no -out:prefix prefix -overwrite  -loops:frag_sizes 9 3 1 -loops:frag_files my_structure.200.9mers.gz my_structure.200.3mers.gz*

*Generation of loophash databases*

We generated the loophash databases using the *loophash_createfiltereddb* application and the VALL database distributed with Rosetta. The command line is:

*mpirun -np 32 loophash_createfiltereddb.mpi.linuxgccrelease -lh:db_path loophash_db/ -in:file:vall path_to_rosetta_tools_repository/tools/fragment_tools/vall.jul19.2011.gz -lh:loopsizes 3 4 5 6 7 8 9 10 11 12 13 14 -lh:num_partitions 32 -lh:createdb_rms_cutoff 4.5 6 7.5 9 10.5 12 13.5 15 16.5 18 19.5 21*

where *loophash_db* is the output path.

### Rosetta scripts & command line for LHKIC

We used the following Rosetta script to run the LHKIC benchmark simulations:

```
<ROSETTASCRIPTS>
   <MOVERS>
      <LoopModeler name="modeler" config="loophash_kic" loops_file="%%loop_file%%"
fast="%%fast%%" />
   </MOVERS>
   <PROTOCOLS>
      <Add mover_name="modeler"/>
   </PROTOCOLS>
</ROSETTASCRIPTS>
```

Let the home directory of Rosetta be */path/to/rosetta/main*; the input structure be my_structure.pdb; the loop file be *my_structure.loop*; the Rosetta script XML file be loopmodel.xml; and the path to the loophash database be *path_to_loophash_db*. Then the command line to run one simulation is:

*/path/to/rosetta/main/source/bin/rosetta_scripts.mysql.linuxgccrelease -database /path/to/rosetta/main/main/database -in:file:s my_structure.pdb -parser:protocol loopmodel.xml -parser:script_vars loop_file=my_structure.loop fast=no -out:prefix prefix -overwrite -lh:loopsizes 6 8 10 -lh:db_path path_to_loophash_db*

### Rosetta scripts & command line for simulations with native bond lengths and angles

For control simulations that use native bond lengths and angles as input, we replaced the Rosetta script XML file with the following:

```
<ROSETTASCRIPTS>
   <TASKOPERATIONS>
```

```
      <RestrictToLoops name="loop" loops_file="%%loop_file%%"/>
   </TASKOPERATIONS>
   <MOVERS>
      <LoopModeler
         name="modeler"
         config="kic_with_frags"
         loops_file="%%loop_file%%"
         fast="%%fast%%">
         <Build skip=”True” />
      </LoopModeler>
   </MOVERS>
   <PROTOCOLS>
      <Add mover_name=”modeler”/>
   </PROTOCOLS>
</ROSETTASCRIPTS>
```

**Input files for PIP**

<u>E38 constraints definition</u>
We used the following energetic constraints file for PIP Version 1 and Version 2:

```
CoordinateConstraint CG  38 CA 1 12.159  64.031 -28.170 HARMONIC 0.0 1.0
CoordinateConstraint OE1 38 CA 1 10.881  63.345 -30.090 HARMONIC 0.0 1.0
CoordinateConstraint OE2 38 CA 1 11.759  65.409 -30.106 HARMONIC 0.0 1.0
```

<u>Additional constraint on F54 for PIP version 2</u>
For PIP Version 2, we included the following additional constraint to prevent a conformational change in the sidechain of F54 that was frequently observed in PIP Version 1:

```
CoordinateConstraint CZ  54 CA 1 15.196  64.334 -29.952 HARMONIC 0.0 1.0
```

<u>Loop definitions for design V1D8r</u>
We used the following loop definition for PIP Version 1 models with a single deletion in the loop region:

```
LOOP 34 45 45 0 1
```

The loop definition for input models with no deletions included residue 46.

<u>Loop definitions for design V2D9r</u>
For PIP Version 2, we used the following loop definitions:

```
LOOP  26  51  40 0 0
```

```
LOOP 198 203 200 0 0
```

<u>Design step resfile for PIP Version 1</u>

We used the following resfile for PIP Version 1 (for input models with a single deletion):

```
NATRO
START
# Design residues in the loop itself. Don't move the catalytic residue,
# because we want to find designs which stabilize that rotamer.
34 - 37 A NOTAA HC
38 A NATRO
39 - 45 A NOTAA HC
# Design any residue that has a sidechain atom within 4A of any loop atom in
# any input model. Phe53, Ala113, and Phe115 are excluded because they are
# known to be important for positioning the catalytic residue.
29 A NOTAA HC
30 A NOTAA HC
31 A NOTAA HC
32 A NOTAA HC
33 A NOTAA HC
46 A NOTAA HC
48 A NOTAA HC
49 A NOTAA HC
50 A NOTAA HC
52 A NOTAA HC
54 A NOTAA HC
56 A NOTAA HC
57 A NOTAA HC
108 A NOTAA HC
109 A NOTAA HC
110 A NOTAA HC
111 A NOTAA HC
112 A NOTAA HC
114 A NOTAA HC
116 A NOTAA HC
117 A NOTAA HC
120 A NOTAA HC
198 B NOTAA HC
199 B NOTAA HC
200 B NOTAA HC
201 B NOTAA HC
# Repack any residue that has a sidechain atom within 6A of any loop atom in
# any input model.
10 A NATAA
11 A NATAA
13 A NATAA
14 A NATAA
15 A NATAA
16 A NATAA
17 A NATAA
```

18 A NATAA
23 A NATAA
26 A NATAA
27 A NATAA
28 A NATAA
47 A NATAA
51 A NATAA
53 A NATAA
55 A NATAA
58 A NATAA
59 A NATAA
60 A NATAA
62 A NATAA

For inputs with no deletions, we included the additional loop residue with the tag NOTAA HC and adjusted the residue numbers of post-loop positions accordingly.

Design step resfile for PIP Version 2
We used the following resfile for PIP version 2:

```
NATRO
START
34     A NOTAA CH
35     A NOTAA CH
36     A NOTAA CH
37     A NOTAA CH
38     A PIKAA E
39     A NOTAA CH
40     A NOTAA CH
41     A NOTAA CH
42     A NOTAA CH
43     A NOTAA CH
44     A NOTAA CH
45     A NOTAA CH
46     A NOTAA CH
199    B NOTAA CH
200    B NOTAA CH
201    B NOTAA CH
202    B NOTAA CH

# Repack positions
# ================
# The following repack positions were chosen by the clash-based repack
# shell creator (excluding the ligand).

14     A NATAA
```

```
30      A NATAA
50      A NATAA
51      A NATAA
54      A NATAA
55      A NATAA
95      A NATAA
109     A NATAA
111     A NATAA
112     A NATAA
113     A NATAA
114     A NATAA
115     A NATAA
116     A NATAA
121     A NATAA
127     B NATAA
204     B NATAA
225     B NATAA
227     B NATAA

# The following repack positions were added after visual inspection of
# clash-based repack shell.

10      A NATAA
13      A NATAA
17      A NATAA
25      A NATAA
52      A NATAA
53      A NATAA
56      A NATAA
57      A NATAA
58      A NATAA
108     A NATAA
110     A NATAA
117     A NATAA
118     A NATAA
126     B NATAA
128     B NATAA
228     B NATAA
```

**PIP Step 1: Build Models**

We used the following command line for the Build Models step in PIP Version 1. Let the "main" directory in Rosetta be */path/to/rosetta/main/*; the (relaxed) input PDB be $INPUT_PDB; the (unrelaxed) native PDB be $NATIVE_PDB; the full-atom and centroid scorefunction parameters for the talaris2013 scorefunction be "EQU.fa.params" and "EQU.cen.params", respectively; the resfile be $RESFILE; the constriants file be $CONSTRAINTS and the loops file be $LOOP.

```
/path/to/rosetta/main/source/bin/loopmodel.linuxgccrelease \
-in:file:s $INPUT_PDB \
-in:file:native $NATIVE_PDB \
-in:file:extra_res_fa "EQU.fa.params" \
-in:file:extra_res_cen "EQU.cen.params" \
-in:file:fullatom \
-out:overwrite \
-out:pdb_gz \
-packing:ex1 \
-packing:ex2 \
-packing:extrachi_cutoff 0 \
-packing:resfile $RESFILE \
-constraints:cst_fa_weight 1.0 \
-constraints:cst_fa_file $CONSTRAINTS \
-loops:loop_file $LOOP \
-loops:remodel "perturb_kic" \
-loops:refine "refine_kic" \
-loops:kic_rama2b \
-loops:kic_omega_sampling \
-loops:allow_omega_move "true" \
-loops:ramp_fa_rep \
-loops:ramp_rama \
```

For PIP Version 2, we used the following Rosetta script to build new backbone geometries. Let the loops file be $LOOPS_PATH. Definitions common to all steps are found in "shared_defs.xml", described below. For all Rosetta scripts, variables are filled in by the PIP package.

```
<ROSETTASCRIPTS>

  {% include "shared_defs.xml" %}

  <TASKOPERATIONS>
    <RestrictToRepacking name="repackonly"/>
  </TASKOPERATIONS>

  <MOVERS>
    <LoopModeler name="modeler"
```

```
    config="loophash_kic"
    scorefxn_fa="scorefxn_cst"
    task_operations="resfile,repackonly,ex,aro,curr"
    loops_file="$LOOPS_PATH"
    loophash_perturb_sequence="yes"
    loophash_seqposes_no_mutate="38"
    fast="no"
  />
 </MOVERS>

 <PROTOCOLS>
  <!-- Constraints read from command line -->
  <Add mover_name="modeler"/>
  <Add mover_name="writer"/>
 </PROTOCOLS>

 <OUTPUT scorefxn="scorefxn"/>

</ROSETTASCRIPTS>
```

The PIP package also builds the command line, but a representative example is shown below. Let the path to the “main” folder in Rosetta be */path/to/rosetta/main/*; the (relaxed) input PDB be $INPUT_PDB; the (unrelaxed) native PDB be $NATIVE_PDB; the folder where models are to be saved be $OUPUT_FOLDER; the name of the particular design be $OUTPUT_NAME; the “build models” Rosetta script be *build_models.xml*; the resfile path be $RESFILE_PATH; the path to the constraints file be $CONSTRAINTS, and the path to the loophash database be *path_to_loophash_db*.

```
/path/to/rosetta/main/source/bin/rosetta_scripts.linuxgccrelease \
-database /path/to/rosetta/main/database/ \
-in:file:s $INPUT_PDB \
-in:file:native $NATIVE_PDB \
-out:prefix $OUTPUT_FOLDER \
-out:suffix $OUTPUT_NAME \
-out:no_nstruct_label -out:overwrite -out:pdb_gz \
-out:mute protocols.loops.loops_main \
-parser:protocol build_models.xml \
-packing:resfile $RESFILE_PATH \
-constraints:cst_fa_file $CONSTRAINTS \
-lh:loopsizes 6 7 8 9 10 11 12 13 14 \
-lh:db_path path_to_loophash_db
```

**PIP Step 2: Design Models**

We used the following command line to design models in step 2 of PIP Version 1. Let the “main” directory in Rosetta be */path/to/rosetta/main/*; the (relaxed) input PDB be $INPUT_PDB; the full-atom and centroid scorefunction parameters be “EQU.fa.params” and “EQU.cen.params”, respectively, and the resfile be $RESFILE.

```
/path/to/rosetta/main/source/bin/fixbb.linuxgccrelease \
-in:file:s $INPUT_PDB \
-in:file:extra_res_fa "EQU.fa.params" \
-in:file:extra_res_cen "EQU.cen.params" \
-out:overwrite \
-out:pdb_gz \
-packing:ex1 \
-packing:ex2 \
-packing:extrachi_cutoff 0 \
-packing:use_input_sc \
-packing:resfile $RESFILE \
```

For PIP Version 2, we defined a custom fold tree for the design step:

```
FOLD_TREE
EDGE   1  39 -1
EDGE 100   1  3
EDGE 100  40 -1
EDGE 100 125 -1
EDGE 100 225  1
EDGE 100 251  2
EDGE 126 200 -1
EDGE 225 126  4
EDGE 225 201 -1
EDGE 225 250 -1
```

We then used the following Rosetta script to design sequences for the new backbone geometries. Let the path to the fold tree file be $FOLDTREE. Definitions common to all steps are found in “shared_defs.xml”, described below.

```
<ROSETTASCRIPTS>

  {% include "shared_defs.xml" %}

  <RESIDUE_SELECTORS>
    <Index name="turn" resnums="200-201"/>
  </RESIDUE_SELECTORS>
```

```
  <TASKOPERATIONS>
    <LayerDesign name="layer"
        ignore_pikaa_natro="yes"/>
    <ConsensusLoopDesign name="abego"
        residue_selector="turn"
        include_adjacent_residues="no"/>
  </TASKOPERATIONS>

  <MOVERS>
    <AtomTree name="foldtree" fold_tree_file="$FOLDTREE"/>
    <AtomTree name="unfoldtree" simple_ft="yes"/>
    <AddChainBreak name="break_loop" resnum="39" change_foldtree="no"/>
    <AddChainBreak name="break_turn" resnum="200" change_foldtree="no"/>
    <FastDesign name="fastdesign"
        task_operations="resfile,layer,abego,ex,aro,curr"
        scorefxn="scorefxn_cst" >
      <MoveMap bb="no" chi="yes" jump="no">
        <Span begin="26"  end="51"  chi="yes" bb="yes"/>
        <Span begin="198" end="203" chi="yes" bb="yes"/>
      </MoveMap>
    </FastDesign>
  </MOVERS>

  <PROTOCOLS>
    <Add mover_name="nativebonus"/>
    <Add mover_name="cst"/> <!-- Added via mover b/c command-line ignored. -->
    <Add mover_name="foldtree"/>
    <Add mover_name="break_loop"/>
    <Add mover_name="break_turn"/>
    <Add mover_name="fastdesign"/>
    <Add mover_name="unfoldtree"/> <!-- Otherwise Foldability segfaults. -->
    <Add mover_name="writer"/>
  </PROTOCOLS>

  <OUTPUT scorefxn="scorefxn"/>

</ROSETTASCRIPTS>
```

A representative command line is shown below. Let the path to the “main” folder in Rosetta be */path/to/rosetta/main/*; the (relaxed) input PDB be $INPUT_PDB; the (unrelaxed) native PDB be $NATIVE_PDB; the folder where models are to be saved be $OUPUT_FOLDER; the name of the particular design be $OUTPUT_NAME; the “design models” Rosetta script be *design_models.xml*, and the resfile path be $RESFILE_PATH.

```
/path/to/rosetta/main/source/bin/rosetta_scripts.linuxgccrelease \
-database /path/to/rosetta/main/database/ \
-in:file:s $INPUT_PDB \
-in:file:native $NATIVE_PDB \
-out:prefix $OUTPUT_FOLDER \
-out:suffix $OUTPUT_NAME \
-out:no_nstruct_label -out:overwrite -out:pdb_gz \
-out:mute protocols.loops.loops_main \
-parser:protocol design_models.xml \
-packing:resfile $RESFILE_PATH \
```

**PIP Step 3: Structure Prediction**

We used the following command line to design models in step 2 of PIP Version 1. Let the “main” directory in Rosetta be */path/to/rosetta/main/*; the (relaxed) input PDB be $INPUT_PDB; the full-atom and centroid scorefunction parameters be “EQU.fa.params” and “EQU.cen.params”, respectively, and the loops file be $LOOPS.

```
/path/to/rosetta/main/source/bin/loopmodel.linuxgccrelease \
-in:file:s $INPUT_PDB \
-in:file:native $NATIVE_PDB \
-in:file:extra_res_fa "EQU.fa.params" \
-in:file:extra_res_cen "EQU.cen.params" \
-in:file:fullatom \
-out:pdb_gz \
-out:overwrite \
-packing:ex1 \
-packing:ex2 \
-packing:extrachi_cutoff 0 \
-loops:loop_file $LOOPS \
-loops:remodel "perturb_kic" \
-loops:refine "refine_kic" \
-loops:kic_rama2b \
-loops:kic_omega_sampling \
-loops:ramp_fa_rep \
-loops:ramp_rama \
```

For PIP Version 2, we used the following Rosetta script to predict the structures of picked designs. Let the loops file be $LOOPS_PATH. Definitions common to all steps are found in “shared_defs.xml”, described below.

```
<ROSETTASCRIPTS>
```

```
  {% include "shared_defs.xml" %}

  <MOVERS>
    <LoopModeler name="modeler"
      config="kic_with_frags"
      scorefxn_fa="scorefxn"
      loops_file="$LOOPS_PATH"
      fast="no">
        <Build skip="yes"/>
    </LoopModeler>
  </MOVERS>

  <PROTOCOLS>
    <Add mover_name="modeler"/>
    <Add mover_name="writer"/>
  </PROTOCOLS>

  <OUTPUT scorefxn="scorefxn"/>

</ROSETTASCRIPTS>
```

A representative command line for PIP Version 2 is shown below. Let the path to the “main” folder in Rosetta be */path/to/rosetta/main/*; the (relaxed) input PDB be $INPUT_PDB; the (unrelaxed) native PDB be $NATIVE_PDB; the folder where models are to be saved be $OUPUT_FOLDER; the name of the particular design be $OUTPUT_NAME; the “predict models” Rosetta script be *predict_models.xml*; the paths to the 9-mer fragments for the first and second loop be *path_to_9mers_A* and *path_to_9mers_B*, respectively, and the paths to 3-mer fragments for the first and second loop be *path_to_3mers_A* and *path_to_3mers_B*, respectively.

```
/path/to/rosetta/main/source/bin/rosetta_scripts.linuxgccrelease \
-database /path/to/rosetta/main/database/ \
-in:file:s $INPUT_PDB \
-in:file:native $NATIVE_PDB \
-out:prefix $OUTPUT_FOLDER \
-out:suffix $OUTPUT_NAME \
-out:no_nstruct_label -out:overwrite -out:pdb_gz \
-out:mute protocols.loops.loops_main \
-parser:protocol predict_models.xml \
-loops:frag_sizes 9 9 3 3 \
-loops:frag_files path_to_9mers_A path_to_9mers_B path_to_3mers_A path_to_3mers_B
```

**PIP Version 2 Shared Definitions**

The following Rosetta script was included in all steps for PIP Version 2. Let the path to the scorefunction weights file be $SCOREFXN_WEIGHTS and the path to the constraints file be $CONSTRAINTS. The weights file was identical to the default weights for the ref2015 scorefunction.

```
{% include "filters.xml" %}

<SCOREFXNS>
  <ScoreFunction name="scorefxn" weights="$SCOREFXN_WEIGHTS"/>
  <ScoreFunction name="scorefxn_cst" weights="$SCOREFXN_WEIGHTS">
    <Reweight scoretype="coordinate_constraint" weight="1.0"/>
    <Reweight scoretype="atom_pair_constraint" weight="1.0"/>
    <Reweight scoretype="angle_constraint" weight="1.0"/>
    <Reweight scoretype="dihedral_constraint" weight="1.0"/>
    <Reweight scoretype="res_type_constraint" weight="1.0"/>
    <Reweight scoretype="chainbreak" weight="100.0"/>
  </ScoreFunction>
</SCOREFXNS>

<RESIDUE_SELECTORS>
  <Chain name="chA" chains="A"/>
  <Index name="E38" resnums="38"/>
</RESIDUE_SELECTORS>

<TASKOPERATIONS>
  <ReadResfile name="resfile"/>
  <ExtraRotamersGeneric name="ex" ex1="yes" ex2="yes" extrachi_cutoff="0"/>
  <LimitAromaChi2 name="aro" include_trp="yes"/>
  <IncludeCurrent name="curr"/>
</TASKOPERATIONS>

<MOVERS>
  <FavorNativeResidue name="nativebonus" />
  <ConstraintSetMover name="cst" cst_fa_file="$CONSTRAINTS"/>
  <WriteFiltersToPose name="writer" prefix="EXTRA_METRIC "/>
</MOVERS>
```

**Filters for PIP Version 2**

We used the following Rosetta script to run filters for all three steps in PIP Version 2, with the exceptions of the fragment quality metric (FragmentScoreFilter) and the Foldability metric, which were not included in the Structure Prediction step. For fragment picking, several variables are defined. Let the folder where fragment picking files are stored be $OUTPUT_DIR; the job-specific name of these files be $OUTPUT_NAME; the path to the required CSBLAST, BLAST, PSIPRED, and SPARKS-X programs be */path/to/csblast-2.2.3_linux64*, */path/to/blast-2.2.26/bin/blastpgp*, */path/to/psipred/runpsipred_single*, and */path/to/sparks-x*, respectively; the path to the BLAST database consisting of FASTA-formatted sequence information for proteins in the PDB be */path/to/BLAST/sequences*; the weights file for scoring fragments be $FRAMGNET_WEIGHTS, and the path to the vall database */path/to/Rosetta/database/sampling/vall.jul19.2011.torsions.gz*.

```
<FILTERS>
  <PackStat
    name="PackStat Score [+]"
    threshold="0"
    chain="0"
    repeats="1"
  />
  <ResidueIE
    name="E38 Interaction Energy [-]"
    scorefxn="scorefxn_cst"
    score_type="total_score"
    energy_cutoff="-10"
    restype3="GLU"
    interface="0"
    whole_pose="0"
    selector="E38"
    jump_number="1"
    interface_distance_cutoff="8.0"
    max_penalty="1000.0"
    penalty_factor="1.0"
  />
  <PreProline
    name="Pre-Proline Potential [-]"
    use_statistical_potential="true"
  />
  <TotalSasa
    name="Total SASA [-]"
    threshold="0"
    upper_threshold="1000000000000000"
    hydrophobic="0"
    polar="0"
  />
  <ExposedHydrophobics
    name="Exposed Hydrophobic Residue SASA [-]"
    sasa_cutoff="20"
    threshold="-1"
```

```
/>
<HbondsToResidue
  name="H-bonds to E38 [+]"
  scorefxn="scorefxn_cst"
  partners="0"
  energy_cutoff="-0.5"
  backbone="true"
  bb_bb="true"
  sidechain="true"
  residue="38"
  from_other_chains="true"
  from_same_chain="true"
/>
<HbondsToResidue
  name="H-bonds to E38 (Backbone) [+]"
  scorefxn="scorefxn_cst"
  partners="0"
  energy_cutoff="-0.5"
  backbone="true"
  bb_bb="true"
  sidechain="false"
  residue="38"
  from_other_chains="true"
  from_same_chain="true"
/>
<HbondsToResidue
  name="H-bonds to E38 (Sidechain) [+]"
  scorefxn="scorefxn_cst"
  partners="0"
  energy_cutoff="-0.5"
  backbone="false"
  bb_bb="false"
  sidechain="true"
  residue="38"
  from_other_chains="true"
  from_same_chain="true"
/>
<BuriedUnsatHbonds
  name="Buried Unsatisfied H-Bonds [-]"
  scorefxn="scorefxn"
  print_out_info_to_pdb="true"
  task_operations="resfile"
/>
<OversaturatedHbondAcceptorFilter
```

```
  name="Oversaturated H-bonds [-]"
  scorefxn="scorefxn_cst"
  max_allowed_oversaturated="0"
  hbond_energy_cutoff="-0.5"
  consider_mainchain_only="false"
/>
<RepackWithoutLigand
  name="Repack Without Ligand (delta REU) [-]"
  scorefxn="scorefxn_cst"
  target_res="all_repacked"
  rms_threshold="100"
/>
{% if w.focus_name != 'validate_designs' %}
<Foldability
  name="Foldability (35-41)"
  tries="60"
  start_res="35" {# Unaffected by loop length. #}
  end_res="41"   {# Unaffected by loop length. #}
/>
<Foldability
  name="Foldability (37-44)"
  tries="60"
  start_res="37" {# Unaffected by loop length. #}
  end_res="44"   {# Unaffected by loop length. #}
/>
<FragmentScoreFilter
  name="Max 9-Residue Fragment RMSD (C alpha) [-]"
  scoretype="FragmentCrmsd"
  sort_by="FragmentCrmsd"
  threshold="9999"
  direction="-"
  start_res="26 "
  end_res="51"
  compute="maximum"
  outputs_folder="$OUTPUT_DIR"
  outputs_name="$OUTPUT_NAME"
  csblast="/path/to/csblast-2.2.3_linux64"
  blast_pgp="/path/to/blast-2.2.26/bin/blastpgp"
  placeholder_seqs="/path/to/BLAST/sequences"
  psipred="/path/to/psipred/runpsipred_single"
  sparks-x="/path/to/sparks-x"
  sparks-x_query="/path/to/sparks-x/bin/buildinp_query.sh"
  frags_scoring_config="$FRAGMENTS_WEIGHTS"
  n_frags="200"
```

```
    n_candidates="1000"
    fragment_size="9"
    vall_path="/path/to/Rosetta/main/database/sampling/vall.jul19.2011.torsions.gz"
    print_to_pdb="true"
  />
  {% endif %}
</FILTERS>
```

We used the following weights file to pick fragments for the fragment quality metric during the structure prediction step:

```
# score name          priority  wght   min_allowed  extras
ProfileScoreL1           700     1.0     -
ProfileScoreStructL1     100     4.0     -
SolventAccessibility     500     1.5     -
Phi                      300     1.0     -
Psi                      200     0.6     -
SecondarySimilarity      600     1.0     -    predA
RamaScore                400     0.8     -    predA
FragmentCrmsd              0     0.0     -
```

### SUPPLEMENTARY NOTE 1

#### Analysis of failure cases

Despite the performance improvements with FKIC, in particular for the challenging *Mixed Segment* and *Multiple Segments* datasets, FKIC failed to accurately model 12 of the 45 segments in the *Standard* 12-residue benchmark dataset. In four cases (1cs6, 1msc, 2tgi and 4i1b) no sub-Å model was generated. In the other eight cases (1arb, 1bhe, 1cyo, 1m3s, 1onc, 1qlw, 1t1d and 1thg), sub-Å models were generated but could not be identified by energy (the RMSDs of lowest energy structures were larger than 1.1Å). These failures could result from deficiencies in sampling near-native conformations, from inaccuracies in the energy function and/or from problems with the crystal structure conformation such as effects of crystal packing. Sampling and energy function errors are often coupled, as the energy function guides sampling during the simulations. To gain insights into potential reasons for the failures we observed, we ran simulations of the failed proteins starting from their native structures as inputs. In these simulations, we skipped the first build stage (yellow in **Supplementary Fig. 1**) so that the native bond angles and bond lengths were kept.

The results of these simulations allowed us to classify failure cases into 4 categories (**Supplementary Fig. 2**):

(1) Only the native-input simulations generate sub-Å models, which are correctly identified by energy. This occurred in two of the 12 failure cases (1cs6 and 2tgi). As the energies of native-like models are much lower than the non-native decoys (**Supplementary Fig. 2a**), failures in these cases are most likely due to the insufficient sampling.

(2) Both standard and native-input FKIC generate sub-Å models, but these models are only correctly identified by lowest energy in the native-input simulations (three of the 12 failure cases: 1bhe, 1onc and 1t1d). As the native-input simulations generated a larger number of correct models with lower energies (**Supplementary Fig. 2b**), the failures are likely caused by the failure of Rosetta to efficiently sample near-native energy minima. One of the possible explanations is that the standard simulation idealizes bond lengths and bond angles in standard FKIC. Because of the rugged energy landscape, small conformational changes can result in significant energy differences.

(3) As in (2), both simulations generate sub-Å models and native-input simulations correctly identify these models by energy, but standard FKIC generates incorrect models with lower energies (**Supplementary Fig. 2c,** two of the 12 failure cases: 1arb and 1qlw). This behavior indicates linked scoring and sampling deficiencies.

(4) Neither standard nor native input simulations generate sub-Å models (**Supplementary Fig. 2d**; five of 12 failure cases: 1cyo, 1m3s, 1msc, 1thg and 4i1b). While these simulations start from the native backbones, they do not include crystal contacts. Because crystal packing affecting loop conformations is well known[1,2], there is a formal possibility that the failures of 1cyo, 1m3s, 1msc and 4i1b are due to

crystal packing. For the lowest energy models for 1cyo and 4i1b, the incorrectly modeled loops would have unfavorable contacts in the crystal lattice. For 1m3s and 1msc, the native loop conformations make contacts with another monomer in the crystal. For 1thg, the RMSD of the lowest energy structure improved from 1.86Å to 1.12Å when including native bond lengths and angles, so there might be both sampling and energy function problems.

In sum, in particular for categories (1) and (2), it could be beneficial to incorporate sampling of bond lengths and bond angles, which we kept to their idealized values to reduce the conformational space to be sampled. Category (3) is indicative of energy function failures although we note that sampling and scoring are coupled in our simulations that accept or reject models based on their energies. Category (4) identifies a number of cases where crystal packing may influence the conformation of the modeled segment in the experimentally determined structure.

### SUPPLEMENTARY NOTE 2

#### Benchmark LHKIC on structure prediction

The LHKIC method was developed for loop design, but it can also be used for structure prediction. We benchmarked the prediction performance of LHKIC on the *Standard* 12 residue dataset (**Supplementary Table 1**). The performance of LHKIC is comparable to NGK and FKIC in terms of median RMSD of lowest scoring models. The median sub-A fraction of LHKIC is 24.35%, better than NGK but worse than FKIC. This result indicates that sequence independent fragments (as in LHKIC) can improve sampling in structure predictions over non fragment-based methods such as NGK, but the improvement is smaller than when using fragments picked with sequence information as in FKIC. Note that the absolute energies for LHKIC cannot be directly compared to the other methods since LHKIC was developed in a newer version of Rosetta (revision 60022, see **Methods**).

### SUPPLEMENTARY NOTE 3

#### 8-Residue and 12-residue *Multiple Segments* datasets

To further benchmark the performance of FKIC on multiple interacting segments, we constructed a dataset of 8-residue interacting segments and a dataset of 12-residue interacting segments in a similar manner to the construction of the 10-residue *Multiple Segments* dataset (see **Methods**). On the 8-residue interacting segment dataset, FKIC has 0.65Å median accuracy and 59.9% median fraction of sub-Å prediction; NGK has 0.79Å median accuracy and 35.6% median fraction of sub-Å prediction (median accuracy and median fraction of sub-Å prediction are described in the main text). On the 12-residue interacting dataset, FKIC has 1.53Å median accuracy and 0.21% median fraction of sub-Å prediction; NGK has 1.94Å median accuracy and 0% median fraction of sub-Å prediction. Thus, FKIC improves the prediction accuracy for multiple interacting segments consistently on different segment lengths and is able to find correct solutions for large conformational search problems, such as the set with two interacting 12-residue segments where previous methods such as NGK and CCD frequently fail (**Supplementary Table 6** and **Supplementary Table 7**).

### SUPPLEMENTARY NOTE 4

#### Loop modeling on template-based models and perturbed datasets

To test the performance of FKIC in contexts where the environment of the remodeled loop is non-native, we benchmarked FKIC on several datasets from ref[3]: a benchmark set from template-based modeling, and three sidechain/backbone perturbed loop datasets (**Supplementary Table 11**). On the side chain perturbed datasets, FKIC performs similarly to NGK, and outperforms the other methods. This behavior is likely due to the fact that surrounding residues are repacked during FKIC or NGK simulations, indicating that these methods can account for slight imperfections in the environment that can be resolved by altering side chain conformations. On the backbone-perturbed dataset, the performance of FKIC is comparable to the reported results of GalaxyLoop-PS2[3], with a median RMSD of lowest-scoring models of 1.68Å and 1.65Å for FKIC and Galaxy-PS2, respectively. On the template-based model dataset, none of the methods performs well, with the median RMSD of lowest scoring models above 3 Å for all methods.

It should be noted that the prediction and evaluation approach described here is not suited to appropriately assess the accuracy of loop modeling methods in environments where the surrounding backbone is perturbed, since the backbone in the environment is not allowed to change during the simulation. In our study, we compare a predicted loop structure to the native loop structure after aligning the surrounding environment of the loop. The native loop is the correct answer when the surrounding environment is unperturbed. However, the native loop conformation may not be compatible with the perturbed backbone and can therefore not be identified as the lowest scoring model in simulations where the backbone of the environment remains in its (unchanged) perturbed conformation. Our analysis on the dataset containing backbone perturbations after MD simulations from ref[3] supports this argument. For each protein in this dataset, we defined the residues within 10 Å from the native loop as surrounding residues. We then superimposed the native structure and the perturbed structure by the backbone heavy atoms of the surrounding residues and calculated heavy atom steric clashes between the native loop and the perturbed surrounding backbone atoms. Two heavy atoms were defined as clashing if their interatomic distance was within 2.5 Å. This analysis revealed that twelve out of the twenty native loops in the dataset contained clashes with their perturbed environments that cannot be resolved with simulations that do not relax the surrounding backbone. When excluding these cases, the FKIC and GalaxyLoop-PS2 median RMSDs improved to 1.3 Å and 1.4 Å, respectively (**Supplementary Table 11**).

These considerations highlight that modeling loops in perturbed environments such as homology models remains an important unsolved problem. To adapt the design-centric loop modeling methods presented here to the problem of homology modeling should include simultaneous or iterative refinement of both loop structures and the environment.

### SUPPLEMENTARY TABLES

#### Supplementary Table 1. Datasets and performance summary

#### a) Comparison of methods

| Dataset | Sampling method | Rosetta energy function | Median RMSD of lowest energy model (Å) | Median RMSD of lowest RMSD model (Å) | Median RMSD all models (Å) | Median sub-A fraction | Median lowest energy (REU) | Median time (s) |
|---|---|---|---|---|---|---|---|---|
| ***Standard*** | KIC* | score12 | 1.05 | NA | NA | 4.30% | NA | NA |
| ***Standard*** | CCD | ref2015 | 1.26 | 0.47 | 3.27 | 2.00% | -709.18 | 2299 |
| ***Standard*** | NGK | ref2015 | 0.64 | 0.37 | 2.70 | 13.00% | -712.65 | 3642 |
| ***Standard*** | FKIC | ref2015 | 0.62 | **0.32** | **1.16** | **47.80%** | **-716.78** | 3456 |
| ***Standard*** | LHKIC | ref2015 | **0.55**** | 0.34 | 2.66 | 24.35% | -655.18*** | 3057 |
| ***Mixed*** | CCD | ref2015 | 1.29 | 0.67 | 3.46 | 0.50% | -719.42 | 4309 |
| ***Mixed*** | NGK | ref2015 | 1.07 | 0.45 | 4.65 | 1.15% | -728.18 | 7341 |
| ***Mixed*** | FKIC | ref2015 | **0.53** | **0.34** | **1.46** | **52.30%** | **-739.38** | 7196 |
| ***Multiple*** | CCD | ref2015 | 1.97 | 0.90 | 2.95 | 0.20% | -557.06 | 5204 |
| ***Multiple*** | NGK | ref2015 | 1.29 | 0.52 | 2.35 | 5.50% | -573.60 | 9472 |
| ***Multiple*** | FKIC | ref2015 | **1.00** | **0.41** | **1.82** | **28.50%** | **-581.42** | 8834 |

REU, Rosetta energy units
* taken from ref.[4]; all other simulations were run using the Rosetta energy function "ref2015" as described in ref.[5]
** bold numbers denote best performance for given dataset
*** REU value not directly comparable to other methods as LHKIC was benchmarked using a more recent Rosetta version (see **Methods**).

#### b) Inclusion of fragments from homologous structures

| Dataset | Sampling method | Rosetta energy function | Median RMSD of lowest scoring model (Å) | Median RMSD of lowest RMSD model (Å) | Median RMSD all models (Å) | Median sub-A fraction | Median lowest energy (REU) | Median time (s) |
|---|---|---|---|---|---|---|---|---|
| ***Standard*** | FKIC | talaris2013 | 0.70 | 0.36 | 1.19 | 44.89% | -274.50 | 1646 |
| ***Standard*** | FKIC (+ homologs) | talaris2013 | **0.59** | **0.33** | **0.85** | **66.80%** | -273.29 | 1919 |

### Supplementary Table 2. *Mixed Segment* dataset detailed performance

| | | CCD | | | | | NGK | | | | | FKIC | | | | |
|---|---|---|---|---|---|---|---|---|---|---|---|---|---|---|---|---|
| **PDB name** | Target segment residues | RMSD of lowest energy model (Å) | Lowest energy (REU) | Lowest RMSD (Å) | Energy of lowest RMSD model (REU) | Fraction sub-Å models | RMSD of lowest energy model (Å) | Lowest energy (REU) | Lowest RMSD (Å) | Energy of lowest RMSD model (REU) | Fraction sub-Å models | RMSD of lowest energy model (Å) | Lowest energy (REU) | Lowest RMSD (Å) | Energy of lowest RMSD model (REU) | Fraction sub-Å models |
| **1a8d** | 275-290 | 3.04 | -984.24 | 1.94 | -944.26 | 0.00 | 2.94 | -992.55 | 0.77 | -984.31 | 0.01 | 0.42 | -1000.57 | 0.38 | -974.06 | 0.04 |
| **1arb** | 165-180 | 4.80 | -510.37 | 3.21 | -493.67 | 0.00 | 6.23 | -528.00 | 1.17 | -507.84 | 0.00 | 6.48 | -531.35 | 2.31 | -509.84 | 0.00 |
| **1bhe** | 340-355 | 2.10 | -820.58 | 0.72 | -815.20 | 0.00 | 0.66 | -843.95 | 0.53 | -824.24 | 0.36 | 0.76 | -835.54 | 0.65 | -825.45 | 0.07 |
| **1bn8** | 38-53 | 3.08 | -937.27 | 1.37 | -928.40 | 0.00 | 0.55 | -946.28 | 0.44 | -941.02 | 0.03 | 0.52 | -948.32 | 0.45 | -943.16 | 0.03 |
| **1c5e** | 57-72 | 2.77 | -691.08 | 1.33 | -524.80 | 0.00 | 3.57 | -706.51 | 0.50 | -699.10 | 0.03 | 0.29 | -709.42 | 0.29 | -709.42 | 0.67 |
| **1cb0** | 91-106 | 0.63 | -662.32 | 0.47 | -651.31 | 0.28 | 0.68 | -668.11 | 0.30 | -663.53 | 0.97 | 0.61 | -683.38 | 0.34 | -663.29 | 0.96 |
| **1cs6** | 46-61 | 5.27 | -779.57 | 0.87 | -766.75 | 0.00 | 1.34 | -789.74 | 0.74 | -781.50 | 0.00 | 1.27 | -796.08 | 0.51 | -770.26 | 0.16 |
| **1dqz** | 103-118 | 0.50 | -1243.04 | 0.36 | -1207.68 | 0.33 | 2.40 | -1255.72 | 0.45 | -1222.43 | 0.01 | 0.35 | -1261.68 | 0.24 | -1198.37 | 0.84 |
| **1ede** | 76-91 | 0.38 | -694.76 | 0.38 | -694.76 | 0.03 | 0.26 | -710.85 | 0.19 | -689.15 | 0.37 | 0.28 | -719.71 | 0.21 | -706.81 | 0.72 |
| **1exm** | 252-267 | 1.45 | -977.38 | 0.66 | -969.81 | 0.01 | 4.62 | -983.36 | 1.38 | -962.42 | 0.00 | 4.38 | -991.09 | 0.38 | -983.50 | 0.21 |
| **1ezm** | 24-39 | 6.88 | -654.25 | 3.31 | -556.10 | 0.00 | 0.50 | -684.52 | 0.40 | -668.73 | 0.00 | 6.00 | -681.88 | 1.37 | -611.59 | 0.00 |
| **1f46** | 45-60 | 5.48 | -705.82 | 1.07 | -701.81 | 0.00 | 7.10 | -723.09 | 0.99 | -708.18 | 0.00 | 1.51 | -735.38 | 1.08 | -704.28 | 0.00 |
| **1i7p** | 155-170 | 0.68 | -647.33 | 0.68 | -647.33 | 0.00 | 1.59 | -654.72 | 1.51 | -648.48 | 0.00 | 0.56 | -657.64 | 0.37 | -652.65 | 0.46 |
| **1ms9** | 431-446 | 10.98 | -2721.15 | 4.21 | -2392.51 | 0.00 | 11.83 | -2744.95 | 3.10 | -2427.70 | 0.00 | 6.42 | -2749.58 | 0.70 | -2079.87 | 0.01 |
| **1oth** | 272-287 | 1.31 | -612.54 | 0.74 | -510.97 | 0.01 | 2.24 | -685.23 | 0.45 | -555.42 | 0.02 | 0.71 | -673.30 | 0.38 | -559.34 | 0.80 |
| **1oyc** | 330-345 | 1.35 | -557.96 | 0.48 | -137.82 | 0.08 | 5.66 | -536.60 | 0.59 | 126.42 | 0.00 | 0.54 | -593.99 | 0.39 | -471.11 | 0.65 |
| **1pbe** | 200-215 | 0.35 | -803.31 | 0.35 | -803.31 | 0.17 | 0.59 | -814.05 | 0.31 | -807.25 | 0.25 | 0.57 | -816.86 | 0.25 | -813.62 | 0.58 |
| **1qlw** | 284-299 | 0.46 | -1557.59 | 0.39 | -1547.37 | 0.09 | 0.52 | -1566.76 | 0.34 | -1561.36 | 0.06 | 0.35 | -1573.51 | 0.18 | -1569.30 | 0.73 |
| **1srp** | 418-433 | 0.53 | -733.70 | 0.53 | -733.70 | 0.00 | 0.67 | -733.27 | 0.65 | -729.35 | 0.01 | 0.42 | -743.39 | 0.29 | -739.06 | 0.32 |
| **1tca** | 163-178 | 2.33 | -829.93 | 1.05 | -815.78 | 0.00 | 0.28 | -850.81 | 0.28 | -850.81 | 0.08 | 0.36 | -852.46 | 0.27 | -850.07 | 0.17 |
| **1thg** | 359-374 | 3.78 | -1290.91 | 0.98 | -798.31 | 0.00 | 7.31 | -836.82 | 2.20 | -782.91 | 0.00 | 0.87 | -820.01 | 0.76 | -793.95 | 0.28 |
| **1thw** | 97-112 | 0.50 | -365.34 | 0.50 | -365.34 | 0.00 | 0.33 | -404.36 | 0.28 | -361.20 | 0.35 | 0.34 | -406.53 | 0.27 | -361.89 | 0.61 |
| **1tib** | 166-181 | 0.49 | -469.26 | 0.49 | -469.26 | 0.07 | 0.30 | -513.50 | 0.25 | -488.46 | 0.71 | 0.29 | -512.87 | 0.25 | -499.93 | 0.81 |
| **1tml** | 26-41 | 0.83 | -733.03 | 0.69 | -729.62 | 0.02 | 0.35 | -756.99 | 0.34 | -745.83 | 0.04 | 0.35 | -764.53 | 0.26 | -756.61 | 0.80 |
| **1xif** | 50-65 | 1.28 | -815.74 | 0.47 | -813.47 | 0.02 | 0.45 | -820.65 | 0.37 | -819.84 | 0.01 | 0.59 | -829.72 | 0.32 | -820.15 | 0.35 |
| **2ebn** | 11-26 | 0.50 | -686.47 | 0.50 | -686.47 | 0.03 | 0.31 | -715.52 | 0.25 | -708.04 | 0.09 | 0.29 | -719.70 | 0.24 | -711.08 | 0.74 |
| **2exo** | 48-63 | 3.51 | -615.43 | 1.17 | -602.03 | 0.00 | 4.79 | -654.47 | 2.30 | -602.47 | 0.00 | 4.88 | -680.12 | 0.43 | -626.40 | 0.07 |
| **2pia** | 43-58 | 1.08 | -666.90 | 0.37 | -594.66 | 0.33 | 1.52 | -683.56 | 0.39 | -581.39 | 0.43 | 0.45 | -682.60 | 0.34 | -600.69 | 0.96 |
| **2sil** | 209-224 | 0.65 | -440.99 | 0.35 | -366.67 | 0.13 | 6.82 | -523.38 | 0.37 | -370.74 | 0.01 | 0.48 | -551.19 | 0.29 | -365.12 | 0.67 |
| **3hsc** | 125-140 | 0.33 | -874.75 | 0.31 | -869.11 | 0.99 | 0.81 | -868.57 | 0.81 | -868.57 | 0.01 | 0.30 | -878.17 | 0.27 | -873.40 | 0.80 |

**Supplementary Table 3.** ***Multiple Segments*** **dataset detailed performance**

| | | **CCD** | | | | | **NGK** | | | | | **FKIC** | | | | |
|---|---|---|---|---|---|---|---|---|---|---|---|---|---|---|---|---|
| **PDB name** | Target segment residues | RMSD of lowest energy model (Å) | Lowest energy (REU) | Lowest RMSD (Å) | Energy of lowest RMSD model (REU) | Fraction sub-Å models | RMSD of lowest energy model (Å) | Lowest energy (REU) | Lowest RMSD (Å) | Energy of lowest RMSD model (REU) | Fraction sub-Å models | RMSD of lowest energy model (Å) | Lowest energy (REU) | Lowest RMSD (Å) | Energy of lowest RMSD model (REU) | Fraction sub-Å models |
| **1a3a** | 11-20<br>87-96<br>170-179 | 5.65 | -340.13 | 0.83 | -284.99 | 0.00 | 0.84 | -316.80 | 0.45 | -246.79 | 0.13 | 0.84 | -325.15 | 0.48 | -254.18 | 0.58 |
| **1deu** | 225-234<br>148-157 | 6.49 | 16511.10 | 1.81 | 16591.50 | 0.00 | 2.21 | 16114.00 | 1.56 | 16135.00 | 0.00 | 2.57 | 16115.50 | 0.97 | 16143.30 | 0.00 |
| **1dqz** | 252-261<br>529-538 | 2.62 | -623.02 | 1.45 | -555.00 | 0.00 | 1.93 | -671.27 | 0.41 | -637.28 | 0.12 | 2.07 | -667.08 | 0.55 | -626.29 | 0.06 |
| **1euv** | 570-579<br>239-248 | 2.17 | -500.01 | 1.56 | -474.99 | 0.00 | 1.88 | -521.07 | 0.96 | -513.85 | 0.00 | 1.10 | -523.17 | 0.93 | -517.57 | 0.01 |
| **1fo9** | 261-270<br>211-220 | 1.09 | -774.02 | 1.09 | -774.02 | 0.00 | 0.84 | -804.34 | 0.43 | -795.79 | 0.67 | 0.81 | -811.44 | 0.40 | -797.06 | 0.83 |
| **1ftr** | 278-287<br>116-125 | 9.14 | -700.66 | 2.23 | -670.62 | 0.00 | 8.51 | -719.42 | 2.41 | -696.58 | 0.00 | 3.02 | -714.95 | 0.77 | -712.83 | 0.00 |
| **1h1n** | 153-162<br>170-179 | 0.49 | -651.04 | 0.43 | -637.42 | 0.17 | 1.29 | -670.91 | 0.25 | -633.85 | 0.36 | 0.36 | -671.26 | 0.23 | -668.07 | 0.60 |
| **1h6u** | 202-211<br>85-94 | 0.37 | -728.01 | 0.36 | -724.78 | 0.45 | 0.37 | -732.42 | 0.32 | -729.53 | 0.28 | 0.42 | -734.81 | 0.28 | -730.74 | 0.38 |
| **1i7k** | 118-127<br>71-80 | 3.33 | -308.26 | 1.79 | -295.82 | 0.00 | 3.11 | -326.13 | 1.51 | -317.13 | 0.00 | 1.95 | -325.51 | 1.06 | -308.24 | 0.00 |
| **1idp** | 111-120<br>251-260 | 0.65 | -289.74 | 0.44 | -286.54 | 0.11 | 4.43 | -302.14 | 0.60 | -296.58 | 0.02 | 0.77 | -306.48 | 0.28 | -296.60 | 0.80 |
| **1inl** | 266-275<br>35-44 | 2.65 | -520.07 | 0.67 | -432.87 | 0.00 | 0.52 | -579.68 | 0.43 | -562.67 | 0.07 | 0.57 | -586.58 | 0.40 | -558.11 | 0.55 |
| **1j7d** | 69-78<br>44-53 | 0.61 | -332.51 | 0.47 | -324.83 | 0.04 | 1.30 | -337.14 | 1.01 | -328.33 | 0.00 | 2.01 | -341.38 | 0.93 | -329.21 | 0.02 |
| **1jfr** | 15-24<br>34-43 | 3.61 | -554.55 | 0.71 | -434.64 | 0.01 | 1.28 | -567.51 | 0.38 | -469.28 | 0.42 | 7.05 | -576.27 | 0.36 | -494.83 | 0.17 |
| **1jfu** | 136-145<br>628-637 | 1.38 | -407.70 | 0.91 | -398.11 | 0.00 | 5.58 | -414.98 | 1.04 | -403.42 | 0.00 | 1.19 | -415.17 | 0.91 | -397.13 | 0.02 |
| **1ku1** | 645-654<br>44-53 | 0.67 | 550.86 | 0.58 | 551.87 | 0.22 | 0.35 | 445.64 | 0.35 | 542.16 | 0.22 | 0.40 | 445.59 | 0.27 | 455.27 | 0.83 |
| **1kzq** | 83-92<br>71-80 | 3.58 | -444.31 | 1.87 | -407.29 | 0.00 | 2.61 | -466.18 | 1.72 | -458.54 | 0.00 | 1.99 | -463.25 | 0.85 | -450.18 | 0.00 |
| **1m0z** | 49-58<br>12-21 | 0.72 | -559.57 | 0.57 | -550.88 | 0.15 | 0.73 | -564.57 | 0.51 | -558.06 | 0.05 | 0.66 | -571.10 | 0.40 | -560.56 | 0.76 |
| **1nxm** | 98-107<br>18-27 | 0.40 | -465.63 | 0.24 | -453.77 | 0.47 | 0.32 | -482.36 | 0.23 | -468.24 | 0.92 | 0.24 | -476.32 | 0.22 | -468.21 | 0.98 |
| **1qwd** | 93-102<br>140-149 | 2.97 | -376.26 | 1.84 | -368.08 | 0.00 | 1.17 | -393.42 | 0.78 | -388.26 | 0.00 | 0.90 | -389.35 | 0.36 | -383.74 | 0.01 |
| **1t6g** | 197-206<br>240-249 | 1.00 | -753.66 | 0.81 | -737.17 | 0.01 | 1.17 | -777.29 | 0.58 | -753.35 | 0.17 | 1.10 | -777.69 | 0.56 | -763.90 | 0.44 |
| **1u09** | 293-302<br>275-284 | 1.85 | -916.14 | 1.23 | -893.76 | 0.00 | 2.13 | -1110.23 | 1.00 | -902.43 | 0.00 | 3.31 | -933.51 | 0.39 | -926.61 | 0.01 |
| **1w0d** | 315-324 | 1.84 | -663.35 | 0.86 | -662.27 | 0.00 | 1.61 | -670.13 | 0.54 | -664.68 | 0.09 | 0.75 | -669.27 | 0.41 | -662.14 | 0.66 |

| | | CCD | | | | | NGK | | | | | FKIC | | | | |
|---|---|---|---|---|---|---|---|---|---|---|---|---|---|---|---|---|
| **PDB name** | Target segment residues | RMSD of lowest energy model (Å) | Lowest energy (REU) | Lowest RMSD (Å) | Energy of lowest RMSD model (REU) | Fraction sub-Å models | RMSD of lowest energy model (Å) | Lowest energy (REU) | Lowest RMSD (Å) | Energy of lowest RMSD model (REU) | Fraction sub-Å models | RMSD of lowest energy model (Å) | Lowest energy (REU) | Lowest RMSD (Å) | Energy of lowest RMSD model (REU) | Fraction sub-Å models |
| **1xdw** | 49-58<br>27-36 | 0.89 | -783.62 | 0.89 | -783.62 | 0.00 | 0.68 | -805.64 | 0.59 | -797.40 | 0.42 | 0.68 | -811.82 | 0.42 | -802.75 | 0.62 |
| **1xg2** | 184-193<br>218-227 | 3.59 | -594.97 | 0.93 | -554.27 | 0.00 | 2.33 | -637.06 | 0.49 | -614.76 | 0.01 | 0.52 | -638.36 | 0.41 | -621.65 | 0.56 |
| **1xsz** | 77-86<br>119-128 | 0.91 | -815.59 | 0.41 | -774.90 | 0.57 | 0.49 | -816.42 | 0.32 | -793.45 | 0.35 | 0.58 | -823.76 | 0.32 | -795.46 | 0.95 |
| **1xwt** | 333-342<br>369-378 | 2.10 | -941.99 | 0.92 | -923.93 | 0.00 | 1.57 | -976.37 | 0.35 | -953.47 | 0.44 | 1.82 | -977.69 | 0.40 | -950.40 | 0.34 |
| **1yif** | 176-185<br>144-153 | 3.06 | -748.46 | 2.18 | -682.37 | 0.00 | 2.50 | -761.17 | 1.81 | -759.79 | 0.00 | 2.92 | -765.53 | 1.88 | -755.31 | 0.00 |
| **1zvt** | 659-668<br>716-725 | 3.72 | -508.86 | 2.52 | -505.57 | 0.00 | 0.32 | -542.65 | 0.32 | -542.65 | 0.07 | 1.49 | -535.43 | 1.28 | -531.60 | 0.00 |
| **2a4a** | 37-46<br>18-27 | 2.44 | -598.61 | 1.60 | -450.13 | 0.00 | 2.01 | -620.40 | 0.71 | -594.80 | 0.00 | 1.84 | -631.78 | 0.93 | -610.00 | 0.00 |
| **2b0a** | 102-111<br>123-132 | 0.45 | -395.53 | 0.45 | -395.53 | 0.00 | 0.42 | -404.93 | 0.42 | -404.93 | 0.04 | 0.38 | -408.45 | 0.31 | -395.35 | 0.24 |

### Supplementary Table 4. *Standard* dataset detailed performance

| | | CCD | | | | | NGK | | | | | FKIC | | | | |
|---|---|---|---|---|---|---|---|---|---|---|---|---|---|---|---|---|
| **PDB name** | Target segment residues | RMSD of lowest energy model (Å) | Lowest energy (REU) | Lowest RMSD (Å) | Energy of lowest RMSD model (REU) | Fraction sub-Å models | RMSD of lowest energy model (Å) | Lowest energy (REU) | Lowest RMSD (Å) | Energy of lowest RMSD model (REU) | Fraction sub-Å models | RMSD of lowest energy model (Å) | Lowest energy (REU) | Lowest RMSD (Å) | Energy of lowest RMSD model (REU) | Fraction sub-Å models |
| **1a8d** | 155-166 | 2.83 | -1008.38 | 2.14 | -973.52 | 0.00 | 0.38 | -1024.27 | 0.38 | -1024.27 | 0.03 | 0.42 | -1022.05 | 0.31 | -1019.72 | 0.01 |
| **1arb** | 182-193 | 2.37 | -554.35 | 0.23 | -530.49 | 0.01 | 0.54 | -562.94 | 0.37 | -539.98 | 0.38 | 2.06 | -563.70 | 0.39 | -526.90 | 0.17 |
| **1bhe** | 121-132 | 2.09 | -923.69 | 1.84 | -913.87 | 0.00 | 0.68 | -908.66 | 0.30 | -903.45 | 0.18 | 1.91 | -915.76 | 0.31 | -906.99 | 0.13 |
| **1bn8** | 298-309 | 0.70 | -987.69 | 0.33 | -982.93 | 0.02 | 0.75 | -993.21 | 0.43 | -986.78 | 0.09 | 0.48 | -1000.64 | 0.30 | -991.29 | 0.44 |
| **1c5e** | 82-93 | 0.40 | -742.69 | 0.30 | -742.50 | 0.09 | 0.36 | -745.93 | 0.28 | -742.69 | 0.41 | 0.38 | -748.17 | 0.31 | -744.34 | 0.90 |
| **1cb0** | 33-44 | 0.41 | -679.12 | 0.30 | -671.31 | 0.10 | 0.25 | -679.95 | 0.16 | -651.78 | 0.36 | 0.36 | -680.65 | 0.14 | -675.44 | 0.84 |
| **1cnv** | 188-199 | 3.09 | -691.00 | 1.24 | -665.03 | 0.00 | 1.06 | -701.43 | 0.47 | -685.34 | 0.01 | 1.02 | -700.87 | 0.39 | -699.85 | 0.03 |
| **1cs6** | 145-156 | 3.60 | -731.51 | 1.71 | -704.86 | 0.00 | 4.26 | -777.82 | 0.99 | -760.81 | 0.00 | 4.24 | -773.94 | 1.55 | -745.58 | 0.00 |
| **1cyo** | 12-23 | 0.91 | -192.93 | 0.58 | -191.35 | 0.03 | 4.98 | -197.51 | 0.53 | -189.01 | 0.02 | 5.09 | -196.90 | 0.51 | -191.38 | 0.01 |
| **1dqz** | 209-220 | 0.40 | -1471.41 | 0.40 | -1471.41 | 0.00 | 0.49 | -1476.24 | 0.30 | -1456.86 | 0.33 | 0.46 | -1480.22 | 0.32 | -1325.32 | 0.32 |
| **1dts** | 41-52 | 7.66 | -445.87 | 0.95 | -436.78 | 0.00 | 6.63 | -470.55 | 0.94 | -436.96 | 0.01 | 1.07 | -472.01 | 0.75 | -417.87 | 0.16 |
| **1eco** | 35-46 | 0.36 | -283.70 | 0.29 | -267.09 | 0.63 | 0.40 | -284.48 | 0.32 | -277.13 | 0.49 | 0.37 | -285.75 | 0.30 | -269.98 | 0.99 |
| **1ede** | 150-161 | 1.26 | -598.32 | 0.76 | -592.01 | 0.01 | 0.72 | -615.96 | 0.59 | -612.93 | 0.05 | 0.77 | -631.71 | 0.54 | -613.56 | 0.21 |
| **1exm** | 291-302 | 0.52 | -1062.06 | 0.47 | -1051.67 | 0.35 | 0.47 | -1048.50 | 0.27 | -1040.46 | 0.62 | 0.63 | -1048.74 | 0.27 | -1039.80 | 0.89 |
| **1ezm** | 122-133 | 2.53 | -733.12 | 1.25 | -719.78 | 0.00 | 0.37 | -758.95 | 0.37 | -758.95 | 0.04 | 0.62 | -753.62 | 0.44 | -747.20 | 0.03 |
| **1f46** | 64-75 | 2.27 | -709.18 | 0.80 | -697.38 | 0.03 | 3.09 | -712.65 | 0.78 | -705.47 | 0.14 | 0.44 | -716.78 | 0.32 | -710.87 | 0.30 |
| **1i7p** | 63-74 | 0.51 | -669.97 | 0.36 | -661.74 | 0.11 | 0.42 | -674.13 | 0.31 | -651.77 | 0.53 | 0.38 | -672.19 | 0.33 | -669.48 | 0.90 |
| **1m3s** | 68-79 | 5.40 | -672.68 | 0.68 | -666.48 | 0.01 | 5.05 | -674.66 | 0.44 | -664.02 | 0.31 | 5.66 | -673.18 | 0.42 | -664.04 | 0.51 |
| **1ms9** | 529-540 | 2.48 | -3018.90 | 0.77 | -2998.79 | 0.04 | 0.27 | -3041.97 | 0.22 | -3041.75 | 0.91 | 0.35 | -3044.20 | 0.23 | -3030.97 | 0.69 |
| **1msc** | 9-20 | 7.68 | 151.93 | 2.93 | 188.30 | 0.00 | 6.54 | 148.08 | 1.27 | 162.88 | 0.00 | 7.85 | 147.53 | 1.08 | 155.56 | 0.00 |
| **1my7** | 254-265 | 0.49 | -504.92 | 0.35 | -496.96 | 0.11 | 0.61 | -511.55 | 0.40 | -507.90 | 0.75 | 0.53 | -512.36 | 0.45 | -507.57 | 0.81 |
| **1onc** | 23-34 | 2.88 | -200.66 | 1.70 | -190.59 | 0.00 | 3.63 | -209.76 | 0.46 | -208.03 | 0.04 | 3.62 | -208.46 | 0.93 | -200.42 | 0.00 |
| **1oth** | 69-80 | 0.52 | -837.57 | 0.38 | -836.26 | 0.18 | 0.45 | -837.27 | 0.31 | -834.46 | 0.34 | 0.50 | -843.18 | 0.27 | -833.69 | 0.97 |
| **1oyc** | 203-214 | 2.99 | -776.66 | 0.89 | -691.71 | 0.00 | 0.30 | -818.69 | 0.28 | -719.38 | 0.13 | 0.32 | -819.70 | 0.25 | -727.24 | 0.54 |
| **1pbe** | 129-140 | 2.60 | -833.07 | 0.43 | -790.79 | 0.06 | 0.45 | -820.65 | 0.32 | -801.51 | 0.32 | 0.38 | -824.99 | 0.33 | -804.46 | 0.72 |
| **1qlw** | 31-42 | 4.34 | -1611.33 | 2.07 | -1604.34 | 0.00 | 4.87 | -1624.91 | 0.30 | -1605.29 | 0.13 | 2.83 | -1626.69 | 0.33 | -1620.54 | 0.06 |
| **1rro** | 17-28 | 0.44 | -272.26 | 0.34 | -270.03 | 0.08 | 0.64 | -277.16 | 0.39 | -270.25 | 0.10 | 0.63 | -279.08 | 0.32 | -269.39 | 0.48 |
| **1srp** | 311-322 | 0.48 | -782.41 | 0.31 | -773.57 | 0.05 | 0.31 | -877.44 | 0.25 | -772.31 | 0.99 | 0.37 | -879.03 | 0.25 | -759.90 | 0.98 |
| **1t1d** | 127-138 | 0.42 | -206.93 | 0.34 | -204.69 | 0.14 | 2.79 | -217.61 | 0.32 | -207.60 | 0.09 | 2.94 | -215.11 | 0.30 | -209.60 | 0.39 |
| **1tca** | 305-316 | 9.52 | -846.00 | 2.37 | -806.46 | 0.00 | 0.66 | -859.34 | 0.26 | -854.54 | 0.12 | 0.29 | -864.98 | 0.24 | -848.65 | 0.24 |
| **1thg** | 127-138 | 2.88 | -1419.28 | 1.64 | -1385.22 | 0.00 | 1.05 | -1434.24 | 0.71 | -1411.26 | 0.07 | 1.86 | -1434.72 | 0.28 | -1423.84 | 0.07 |
| **1thw** | 178-189 | 1.06 | -394.74 | 1.06 | -394.74 | 0.00 | 0.65 | -413.28 | 0.49 | -411.02 | 0.05 | 0.61 | -414.92 | 0.46 | -409.31 | 0.23 |
| **1tib** | 99-110 | 0.64 | -527.93 | 0.39 | -524.55 | 0.10 | 0.75 | -529.58 | 0.40 | -526.29 | 0.07 | 0.63 | -535.99 | 0.39 | -529.46 | 0.57 |
| **1tml** | 243-254 | 0.46 | -766.55 | 0.38 | -765.86 | 0.05 | 0.50 | -776.42 | 0.25 | -773.23 | 0.51 | 0.52 | -777.92 | 0.28 | -774.05 | 0.79 |
| **1xif** | 203-214 | 1.90 | -821.11 | 1.26 | -814.10 | 0.00 | 0.41 | -838.28 | 0.18 | -828.66 | 0.94 | 0.37 | -839.04 | 0.22 | -833.59 | 0.85 |
| **2cpl** | 145-156 | 0.60 | -457.29 | 0.26 | -453.37 | 0.26 | 0.32 | -459.00 | 0.21 | -451.22 | 0.26 | 0.30 | -458.11 | 0.24 | -454.07 | 0.99 |
| **2ebn** | 136-147 | 0.35 | -722.21 | 0.35 | -722.21 | 0.02 | 0.44 | -719.92 | 0.43 | -719.89 | 0.01 | 0.60 | -721.73 | 0.43 | -718.80 | 0.09 |
| **2exo** | 293-304 | 0.61 | -737.11 | 0.41 | -728.13 | 0.12 | 0.49 | -731.90 | 0.42 | -722.45 | 0.18 | 0.47 | -734.53 | 0.35 | -725.14 | 0.99 |
| **2pia** | 30-41 | 1.00 | -686.94 | 0.38 | -662.42 | 0.55 | 0.82 | -688.54 | 0.48 | -680.08 | 0.99 | 0.86 | -688.99 | 0.56 | -673.23 | 1.00 |
| **2rn2** | 90-101 | 1.41 | -283.79 | 1.41 | -283.79 | 0.00 | 0.64 | -297.34 | 0.32 | -289.54 | 0.40 | 0.68 | -298.25 | 0.29 | -290.57 | 0.69 |
| **2sil** | 255-266 | 0.69 | -744.26 | 0.41 | -732.81 | 0.55 | 1.26 | -745.15 | 0.60 | -734.86 | 0.00 | 0.93 | -747.04 | 0.46 | -740.32 | 0.57 |
| **2tgi** | 48-59 | 2.69 | -140.59 | 1.73 | -121.99 | 0.00 | 3.57 | -162.15 | 0.36 | -89.29 | 0.00 | 3.14 | -163.39 | 1.36 | -87.48 | 0.00 |
| **3cla** | 176-187 | 1.32 | -475.15 | 0.41 | -472.78 | 0.21 | 1.21 | -483.29 | 0.39 | -478.16 | 0.04 | 0.64 | -484.91 | 0.28 | -477.79 | 0.63 |
| **3hsc** | 72-83 | 0.51 | -894.79 | 0.46 | -891.28 | 0.04 | 0.32 | -896.76 | 0.25 | -890.33 | 0.72 | 0.56 | -899.54 | 0.25 | -893.37 | 0.94 |
| **4i1b** | 46-57 | 3.63 | -134.68 | 2.14 | -130.51 | 0.00 | 6.69 | -140.23 | 0.82 | -135.44 | 0.00 | 6.63 | -140.98 | 1.04 | -97.29 | 0.00 |

### Supplementary Table 5. Failure cases, *Standard* dataset

| | | FKIC | | | | FKIC native input | | | | Reason for failure |
|---|---|---|---|---|---|---|---|---|---|---|
| **PDB name** | Target segment residues | RMSD of lowest energy model (Å) | Lowest energy (REU) | Lowest RMSD (Å) | Energy of lowest RMSD model (REU) | RMSD of lowest energy model (Å) | Lowest energy (REU) | Lowest RMSD (Å) | Energy of lowest RMSD model (REU) | |
| **1arb** | 182-193 | 2.06 | -563.70 | 0.39 | -526.90 | 0.40 | -557.64 | 0.21 | -537.67 | insufficient sampling |
| **1bhe** | 121-132 | 1.91 | -915.76 | 0.31 | -906.99 | 0.24 | -940.55 | 0.24 | -928.05 | insufficient sampling |
| **1cs6** | 145-156 | 4.24 | -773.94 | 1.55 | -745.58 | 0.39 | -809.13 | 0.25 | -804.74 | insufficient sampling |
| **1cyo** | 12-23 | 5.09 | -196.90 | 0.51 | -191.38 | **1.06*** | -194.88 | 0.32 | -188.95 | crystal packing |
| **1m3s** | 68-79 | 5.66 | -673.18 | 0.42 | -664.04 | **5.39** | -680.83 | 0.43 | -669.76 | crystal packing |
| **1msc** | 9-20 | 7.85 | 147.53 | 1.08 | 155.56 | **8.74** | 146.72 | 0.86 | 163.39 | crystal packing |
| **1onc** | 23-34 | 3.62 | -208.46 | 0.93 | -200.42 | 0.51 | -208.74 | 0.39 | -200.49 | insufficient sampling |
| **1qlw** | 31-42 | 2.83 | -1626.69 | 0.33 | -1620.54 | 0.48 | -1623.97 | 0.29 | -1595.85 | insufficient sampling / energy function deficiency |
| **1t1d** | 127-138 | 2.94 | -215.11 | 0.30 | -209.60 | 0.33 | -220.36 | 0.22 | -217.79 | insufficient sampling |
| **1thg** | 127-138 | 1.86 | -1434.72 | 0.28 | -1423.84 | **1.12** | -1451.32 | 0.23 | -1434.68 | insufficient sampling / energy function deficiency |
| **2tgi** | 48-59 | 3.14 | -163.39 | 1.36 | -87.48 | 0.41 | -189.26 | 0.31 | -182.44 | insufficient sampling |
| **4i1b** | 46-57 | 6.63 | -140.98 | 1.04 | -97.29 | **6.64** | -140.95 | 0.81 | -137.17 | crystal packing |

* bold numbers indicate cases where FKIC native input simulations also failed to correctly identify sub-Å models

### Supplementary Table 6. *Multiple Segments* (12 residues) dataset detailed performance

| | | NGK | | | | | FKIC | | | | |
|---|---|---|---|---|---|---|---|---|---|---|---|
| **PDB name** | Target segment residues | RMSD of lowest energy model (Å) | Lowest energy (REU) | Lowest RMSD (Å) | Energy of lowest RMSD model (REU) | Fraction sub-Å models | RMSD of lowest energy Model (Å) | Lowest energy (REU) | Lowest RMSD (Å) | Energy of lowest RMSD model (REU) | Fraction sub-Å models |
| **1ceo** | 10-21, 53-64 | 3.74 | -786.28 | 2.57 | -763.87 | 0.00 | 2.47 | -801.47 | 1.60 | -774.43 | 0.00 |
| **1deu** | 47-58, 117-128 | 2.07 | 16572.20 | 1.27 | 16605.60 | 0.00 | 1.05 | 16537.10 | 0.53 | 16568.70 | 0.06 |
| **1dqz** | 146-157, 250-261 | 2.78 | -657.72 | 0.89 | -627.68 | 0.00 | 0.80 | -656.04 | 0.80 | -656.04 | 0.00 |
| **1euv** | 553-564, 527-538 | 1.35 | -497.87 | 1.16 | -480.99 | 0.00 | 2.59 | -502.24 | 1.58 | -495.34 | 0.00 |
| **1ftr** | 209-220, 276-287 | 9.25 | -717.41 | 2.57 | -698.25 | 0.00 | 7.33 | -723.89 | 2.19 | -708.61 | 0.00 |
| **1h6u** | 168-179, 190-201 | 1.01 | -715.67 | 1.01 | -715.67 | 0.00 | 0.40 | -745.35 | 0.26 | -733.33 | 0.64 |
| **1i7k** | 138-149, 66-77 | 1.80 | -322.00 | 0.78 | -312.85 | 0.00 | 0.47 | -327.72 | 0.36 | -321.18 | 0.30 |
| **1j7d** | 33-44, 67-78 | 2.30 | -329.13 | 1.35 | -323.97 | 0.00 | 1.98 | -343.43 | 0.92 | -333.10 | 0.01 |
| **1jfu** | 32-43, 134-145 | 5.49 | -409.27 | 1.49 | -391.57 | 0.00 | 1.29 | -415.60 | 0.54 | -400.58 | 0.05 |
| **1ku1** | 677-688, 729-740 | 4.37 | 567.65 | 1.65 | 576.01 | 0.00 | 1.11 | 560.84 | 0.67 | 563.89 | 0.02 |
| **1m0z** | 199-210, 244-255 | 3.35 | -576.11 | 1.60 | -126.88 | 0.00 | 2.17 | -583.89 | 1.12 | 27.73 | 0.00 |
| **1qs1** | 264-275, 333-344 | 1.98 | -957.12 | 1.77 | -923.44 | 0.00 | 1.67 | -970.92 | 0.90 | -957.65 | 0.00 |
| **1ryl** | 54-65, 118-129 | 2.19 | -363.80 | 1.73 | -334.16 | 0.00 | 2.23 | -362.49 | 1.75 | -348.55 | 0.00 |
| **1suu** | 741-752, 780-791 | 6.21 | -711.24 | 2.66 | -698.90 | 0.00 | 3.11 | -708.22 | 2.53 | -692.72 | 0.00 |
| **1t6g** | 195-206, 220-231 | 1.10 | -765.81 | 0.85 | -747.86 | 0.01 | 1.87 | -771.86 | 0.62 | -756.32 | 0.17 |
| **1u09** | 238-249, 291-302 | 2.59 | -945.68 | 1.25 | -916.78 | 0.00 | 2.91 | -952.12 | 0.60 | -929.50 | 0.00 |
| **1v5d** | 282-293, 308-319 | 6.08 | -938.87 | 4.75 | -895.23 | 0.00 | 7.20 | -945.93 | 5.15 | -920.17 | 0.00 |
| **1w0d** | 65-76, 274-285 | 5.47 | -661.63 | 1.25 | -534.61 | 0.00 | 0.65 | -676.38 | 0.31 | -650.39 | 0.44 |
| **1wko** | 38-49, 109-120 | 1.52 | -353.30 | 0.97 | -321.36 | 0.00 | 1.19 | -357.76 | 0.46 | -321.97 | 0.08 |
| **1xwt** | 367-378, 38-49 | 5.47 | -974.75 | 1.52 | -954.86 | 0.00 | 2.62 | -995.65 | 1.07 | -950.23 | 0.00 |
| **1xyz** | 792-803, 813-824 | 7.55 | -786.12 | 3.18 | -737.98 | 0.00 | 1.99 | -800.04 | 0.65 | -767.73 | 0.00 |
| **1yif** | 142-153, 174-185 | 5.15 | -773.41 | 2.34 | -712.48 | 0.00 | 3.84 | -781.24 | 1.91 | -702.07 | 0.00 |
| **1zvt** | 657-668, 714-725 | 0.35 | -545.67 | 0.32 | -545.44 | 0.03 | 1.44 | -542.72 | 1.41 | -526.26 | 0.00 |
| **2ahf** | 182-193, 208-219 | 6.52 | -707.08 | 3.03 | -269.19 | 0.00 | 4.51 | -775.93 | 0.80 | -371.41 | 0.00 |
| **2b49** | 674-685, 652-663 | 0.55 | -574.78 | 0.55 | -574.78 | 0.04 | 2.88 | -579.10 | 0.81 | -572.12 | 0.00 |
| **2c61** | 410-421, 164-175 | 3.31 | -981.07 | 0.74 | -973.54 | 0.00 | 5.14 | -987.68 | 1.29 | -970.39 | 0.00 |
| **2cyg** | 248-259, 294-305 | 0.72 | -728.09 | 0.72 | -728.09 | 0.02 | 1.22 | -753.92 | 0.58 | -723.71 | 0.55 |
| **2e01** | 87-98, 256-267 | 2.02 | -909.67 | 1.61 | -894.78 | 0.00 | 0.44 | -946.66 | 0.31 | -922.67 | 0.75 |
| **2g30** | 790-801, 822-833 | 1.45 | -500.35 | 1.18 | -486.01 | 0.00 | 3.35 | -486.53 | 1.61 | -483.88 | 0.00 |
| **2hjv** | 299-310, 326-337 | 1.78 | -390.05 | 1.08 | -377.63 | 0.00 | 1.44 | -393.43 | 1.35 | -385.84 | 0.00 |

### Supplementary Table 7. *Multiple Segments* (8 residues) dataset detailed performance

| | | NGK | | | | | FKIC | | | | |
|---|---|---|---|---|---|---|---|---|---|---|---|
| **PDB name** | Target segment residues | RMSD of lowest energy model (Å) | Lowest energy (REU) | Lowest RMSD (Å) | Energy of lowest RMSD model (REU) | Fraction sub-Å models | RMSD of lowest energy model (Å) | Lowest energy (REU) | Lowest RMSD (Å) | Energy of lowest RMSD model (REU) | Fraction sub-Å models |
| **1a8d** | 278-285,303-310 | 1.15 | -998.03 | 0.35 | -993.09 | 0.10 | 0.94 | -983.98 | 0.73 | -978.66 | 0.01 |
| **1a8u** | 225-232,251-258 | 0.94 | -1073.51 | 0.94 | -1073.51 | 0.00 | 0.65 | -1081.86 | 0.58 | -1036.74 | 0.42 |
| **1ako** | 150-157,173-180 | 0.45 | -668.54 | 0.26 | -666.92 | 0.66 | 0.43 | -670.50 | 0.29 | -662.28 | 0.68 |
| **1bhe** | 282-289,344-351 | 0.51 | -877.73 | 0.28 | -837.52 | 0.93 | 0.68 | -875.49 | 0.38 | -865.22 | 0.24 |
| **1bn8** | 250-257,338-345 | 1.98 | -963.94 | 0.62 | -959.38 | 0.05 | 2.45 | -967.40 | 0.68 | -949.27 | 0.00 |
| **1brt** | 28-35,205-212 | 0.76 | -510.31 | 0.73 | -500.12 | 0.21 | 0.81 | -609.85 | 0.33 | -488.96 | 0.94 |
| **1c5e** | 49-56,97-104 | 0.45 | -702.21 | 0.29 | -695.39 | 0.66 | 0.44 | -703.36 | 0.26 | -697.43 | 0.67 |
| **1cil** | 39-46,80-87 | 0.57 | 178.15 | 0.46 | 199.82 | 0.96 | 0.64 | 187.24 | 0.38 | 217.96 | 1.00 |
| **1cs6** | 126-133,158-165 | 1.10 | -741.75 | 0.33 | -719.69 | 0.57 | 0.72 | -742.82 | 0.43 | -741.61 | 0.73 |
| **1dqz** | 135-142,111-118 | 0.23 | -1264.84 | 0.21 | -1221.96 | 0.98 | 0.23 | -1262.80 | 0.21 | -1149.68 | 0.99 |
| **1ede** | 52-59,283-290 | 0.79 | -689.00 | 0.69 | -680.74 | 0.46 | 0.47 | -694.65 | 0.32 | -684.81 | 0.84 |
| **1exm** | 305-312,255-262 | 0.95 | -998.75 | 0.30 | -980.32 | 0.22 | 0.38 | -994.71 | 0.31 | -970.07 | 0.57 |
| **1gai** | 282-289,401-408 | 0.65 | -306.61 | 0.26 | -292.27 | 0.25 | 0.33 | -294.19 | 0.28 | -288.39 | 0.63 |
| **1gof** | 7-14,31-38 | 0.80 | -1253.43 | 0.72 | -1216.17 | 0.22 | 0.87 | -1228.70 | 0.66 | -1174.50 | 0.41 |
| **1jev** | 451-458,147-154 | 0.79 | -1258.90 | 0.43 | -1242.82 | 0.86 | 0.88 | -1247.44 | 0.37 | -1239.03 | 0.97 |
| **1ms9** | 433-440,468-475 | 2.27 | -2629.19 | 0.32 | -2392.50 | 0.14 | 2.74 | -2411.81 | 0.29 | -2332.46 | 0.02 |
| **1nif** | 64-71,221-228 | 1.49 | -722.30 | 0.93 | -643.85 | 0.00 | 1.50 | -726.32 | 0.51 | -651.66 | 0.14 |
| **1oth** | 219-226,280-287 | 0.30 | -671.55 | 0.22 | -640.91 | 1.00 | 0.27 | -654.61 | 0.22 | -649.22 | 1.00 |
| **1qlw** | 130-137,67-74 | 0.40 | -1600.69 | 0.25 | -1560.94 | 1.00 | 0.37 | -1596.93 | 0.26 | -1592.87 | 1.00 |
| **1srp** | 260-267,294-301 | 0.26 | -760.01 | 0.19 | -758.04 | 0.55 | 0.25 | -760.73 | 0.19 | -758.14 | 0.45 |
| **1tad** | 109-116,159-166 | 0.65 | -1951.28 | 0.36 | -1945.22 | 0.98 | 0.64 | -1964.78 | 0.33 | -1947.35 | 0.76 |
| **1thg** | 351-358,307-314 | 2.63 | -979.60 | 2.07 | -928.81 | 0.00 | 3.48 | -977.31 | 1.80 | -935.77 | 0.00 |
| **1tib** | 171-178,211-218 | 1.08 | -466.52 | 0.67 | -440.39 | 0.37 | 0.54 | -471.50 | 0.29 | -460.77 | 0.53 |
| **1udc** | 32-39,78-85 | 0.62 | -668.79 | 0.54 | -660.04 | 0.06 | 0.59 | -667.57 | 0.46 | -660.73 | 0.19 |
| **3bto** | 256-263,280-287 | 1.15 | -3377.16 | 0.29 | -3110.83 | 0.19 | 1.07 | -3405.40 | 0.34 | -3274.16 | 0.06 |
| **3grs** | 131-138,292-299 | 1.06 | -998.99 | 0.29 | -916.54 | 0.34 | 1.52 | -1000.24 | 0.30 | -929.86 | 0.67 |
| **4pga** | 112-119,216-223 | 0.60 | -1315.25 | 0.24 | -1309.43 | 0.97 | 0.57 | -1380.06 | 0.26 | -1313.87 | 1.00 |
| **6cel** | 132-139,367-374 | 1.36 | -588.32 | 0.97 | -568.66 | 0.00 | 1.70 | -584.39 | 0.81 | -573.68 | 0.00 |

**Supplementary Table 8. Parameters of designs selected for experimental testing.**

| | *Design Name* | *Sequence Cluster* | *Struct Cluster* | *Largest Restraint Dist (Å) in lowest scoring model* | *Loop RMSD (Å)* | *Score Gap (REU)* | *% Sub-Å Restraints* | *% Sub-Å Loops* |
|---|---|---|---|---|---|---|---|---|
| **Full-length** | V1D1r | 1 | 1 | 0.83 | 0.20 | 3.16 | 7.60 | 13.00 |
| | V1D2r | 1 | 1 | 1.05 | 0.38 | 2.05 | 0.20 | 12.20 |
| | V1D3r | 2 | 1 | 0.55 | 0.20 | 6.50 | 7.20 | 4.60 |
| | V1D4 | 2 | 1 | 0.68 | 0.24 | 6.56 | 6.60 | 5.20 |
| | V1D5r | 2 | 1 | 0.92 | 0.27 | 11.88 | 2.20 | 2.00 |
| | V1D6 | 3 | 2 | 2.37 | 0.33 | 0.00 | 0.00 | 17.20 |
| | V1D7 | 3 | 3 | 2.63 | 0.25 | 0.00 | 0.00 | 3.20 |
| **Del1** | V1D8r | 1 | 2 | 0.57 | 0.09 | 4.72 | 11.20 | 31.80 |
| | V1D9r | 1 | 2 | 0.80 | 0.08 | 5.09 | 11.40 | 20.00 |
| | V1D10r | 1 | 2 | 0.87 | 0.18 | 4.51 | 13.20 | 19.40 |
| | V1D11 | 2 | 1 | 2.25 | 0.21 | 0.00 | 0.60 | 10.60 |
| | V1D12r | 2 | 1 | 2.35 | 0.32 | 0.00 | 3.20 | 20.40 |
| | V1D13 | 2 | 1 | 2.36 | 1.22 | 0.00 | 1.40 | 6.80 |
| | V1D14 | 3 | 3 | 2.31 | 0.23 | 0.00 | 0.00 | 4.60 |
| **Full-length** | V2D1 | 3 | 2 | 0.67 | 0.32 | 8.16 | 5.80 | 7.25 |
| | V2D2 | 3 | 2 | 0.83 | 0.73 | 8.97 | 4.23 | 7.04 |
| | V2D3 | 3 | 2 | 0.91 | 0.38 | 5.54 | 8.45 | 14.08 |
| | V2D4 | 3 | 2 | 0.71 | 0.23 | 5.35 | 5.88 | 5.88 |
| | V2D5 | 3 | 2 | 0.72 | 0.24 | 10.31 | 1.41 | 1.41 |
| | V2D6 | 1 | 4 | 0.95 | 0.16 | 0.90 | 19.12 | 63.24 |
| | V2D7 | 5 | 1 | 1.03 | 0.61 | 5.52 | 5.71 | 32.86 |
| | V2D8 | 5 | 1 | 1.05 | 0.81 | 2.92 | 5.88 | 35.29 |
| | V2D9 | 2 | 3 | 1.15 | 0.13 | 0.22 | 4.42 | 71.46 |
| | V2D10 | 1 | 1 | 0.86 | 0.71 | 0.09 | 8.57 | 61.10 |
| | V2D11 | 1 | 1 | 0.87 | 0.66 | 12.02 | 26.70 | 54.92 |

Boxes on the left side indicate whether designs were based off of a full-length input structure or contained a 1-residue deletion (Del1) in the catalytic loop. Sequence cluster: Designs were clustered hierarchically (see **Methods**) according to sequence distance, determined using the BLOSUM80 substitution matrix. Struct Cluster: Designs were clustered (see **Methods**) according to the C/Cα/N/O RMSD for the positions where the backbone was remodeled. Largest Restraint Dist (Å) in lowest scoring model: The furthest distance between any of the atoms in the E38 carboxylate (Cδ, Oε1, or Oε2) and their target positions, for the lowest scoring loop modeling decoy. Loop RMSD (Å): The average RMSD in Å between loop modeling decoys and the input design structure. Score Gap: The difference in fa_attr score between the lowest scoring decoy that puts all of the atoms of the E38 carboxylate less that 1Å from their target positions, and the lowest scoring decoy that puts at least one atom of the E38 carboxylate more than 2Å from its target position. A score gap of 0 indicates that the lowest-scoring decoy was more than 2Å from its target position. % Sub-Å Restraints: The fraction of the loop modeling decoys that are predicted to position the all atoms of the Glu carboxylate less than 1Å from their target positions. % Sub-Å Loops: The fraction of loop modeling decoys that are predicted to position all backbone atoms (C/Cα/N/O) within 1Å RMSD of the input design structure.

**Supplementary Table 9a. List of mutations for PIP version 1 designs.**

| Design Name | Mutations |
|---|---|
| **V1D1r** | T35K, E37T, D38E, P39D, V40A, S42L, E43G, P44G, R45Y, S46Q, A50W, V74N, L115T |
| **V1D2r** | D33N, T35Y, D38E, P39S, V40A, S42Q, E43P, P44K, R45Y, S46W, A49D, E53K, S58Q, V74A, M112A, L115I, E118D |
| **V1D3r** | T35K, E37I, D38E, P39T, V40Q, G41Y, S42P, S46K, V74A, M112A |
| **V1D4** | F30Y, D32P, D33N, T35K, E37I, D38E, P39T, V40Q, G41Y, S42P, S46K, T48R, A49D, A50N, N57E, S58A, K60R, V74A, A75N, N76G, V109I, V110N, S111Y, M112A, R113Q, L115V, E118P |
| **V1D5r** | D33N, T35R, D38E, P39T, V40K, G41Y, S42P, P44D, S46K, S58Q, K60A, V74A, M112A, R113Q |
| **V1D6** | F30Y, A31D, D33T, T35R, V36R, E37N, D38E, P39I, V40G, S42P, E43P, R45L, S46P, T48R, A49D, A50N, E53K, N57E, S58D, K60A, V74T, A75N, N76G, V109I, V110S, S111Y, M112A, R113Q, L115V, E118D |
| **V1D7** | F30Y, A31S, D32P, T35I, V36R, E37R, D38E, P39R, V40Y, G41A, S42K, E43A, P44N, R45P, S46R, T48R, A49D, E53Q, N57E, S58D, K60A, V74T, A75N, N76G, V109I, V110A, S111E, M112A, R113Q, L115I, E118D |
| **V1D8r** | F30Y, D32S, E37T, D38E, P39S, V40F, G41-, S42R, E43P, R45F, S46T, A49E, V74A, A75N, N76G, V109I, S111Y, M112A, R113Q |
| **V1D9r** | E37R, D38E, P39S, V40F, G41-, S42R, E43P, R45F, S46T, S58N, V74S, A75N, S111E, M112A, R113Q |
| **V1D10r** | T35S, E37T, D38E, P39S, V40F, G41-, S42R, E43P, R45F, S46T, S58N, V74S, A75N, M112A |
| **V1D11** | F30Y, D32P, D33N, T35R, V36Y, E37D, D38E, P39I, V40G, G41-, S42F, E43P, P44D, R45T, S46G, A49D, A50N, E53A, N57K, S58Q, V74T, A75N, N76G, V109I, V110A, S111E, M112A, L115I, E118D |
| **V1D12r** | T35Q, V36Y, E37N, D38E, P39I, V40G, G41-, S42F, E43R, P44G, R45D, S58Q, A75N, N76G, S111Y, M112A |
| **V1D13** | F30Y, A31D, D32S, D33T, T35Q, V36Y, E37D, D38E, P39I, V40G, G41-, S42F, E43D, P44G, R45G, A49E, A50N, E53R, N57K, S58Q, V74T, A75N, N76G, V109I, V110A, S111E, M112A, L115I, E118D |
| **V1D14** | F30Y, A31D, D32S, D33T, T35I, V36R, E37Y, D38E, P39Q, V40Y, G41-, S42Y, P44G, R45G, S46K, A49D, A50N, E53K, N57M, S58D, V74T, A75N, N76G, V109I, V110A, S111E, M112A, L115I, E118D |

### Supplementary Table 9b. List of mutations for PIP V2 designs.

| ***Design Name*** | ***Mutations*** |
|---|---|
| **V2D1** | V27A, A28G, L29F, F30L, D33G, A34I, T35K, V36I, E37D, D38E, P39D, V40Q, G41N, S42R, E43K, P44Q, R45V, S46T, G47D, T48A, A50Q, I51K, A73S, A75S |
| **V2D1r** | D33G, A34I, V36I, E37D, D38E, P39D, G41N, S42R, E43K, P44Q, R45V, G47D, A50Q, A73S, A75S |
| **V2D2** | V27A, A28G, L29F, F30L, D33G, A34V, T35K, E37D, D38E, P39D, V40Q, G41K, S42K, E43T, P44T, R45V, S46T, G47D, T48A, A50Q, I51K, A73S, A75D |
| **V2D2r** | D33G, A34V, E37D, D38E, P39D, V40Q, G41K, S42K, E43T, P44T, R45V, G47D, A50Q, A75D |
| **V2D3** | V27A, A28G, L29F, F30L, D33G, A34V, T35Q, V36I, E37D, D38E, P39D, V40Q, G41N, S42K, E43T, P44T, R45V, S46T, G47D, T48A, A50Q, I51K, A73S, A75D |
| **V2D3r** | D33G, A34V, V36I, E37D, D38E, P39D, V40Q, G41N, S42K, E43T, P44T, R45V, G47D, A50Q, A75D |
| **V2D4** | V27A, A28G, L29F, F30L, D33G, A34V, T35Q, V36I, E37D, D38E, P39D, V40Q, G41Q, S42T, E43S, P44T, R45V, S46T, G47D, T48A, A50Q, I51K, A73S, A75K |
| **V2D4r** | D33G, A34V, V36I, E37D, D38E, P39D, V40Q, G41Q, S42T, E43S, P44T, R45V, G47D, A50Q, A75K |
| **V2D5** | V27A, A28G, L29F, F30L, D33G, A34V, V36I, E37L, D38E, P39D, V40Q, G41Q, S42K, E43S, P44T, R45V, S46T, G47D, T48A, A50Q, I51K, A73S, A75D |
| **V2D5r** | D33G, A34V, V36I, E37D, D38E, P39D, V40Q, G41Q, S42K, E43S, P44T, R45V, G47D, A50Q, A75D |
| **V2D6** | F30L, D32S, A34V, V36L, E37W, D38E, P39T, V40S, G41Q, S42D, E43R, P44T, R45Y, S46T, T48N, A49S, A73S, V74N, A75R |
| **V2D6r** | V36L, E37W, D38E, P39T, V40S, G41Q, S42D, E43R, P44T, R45Y, V74N, A75R |
| **V2D7** | A28G, L29Q, A31G, D32P, D33Q, A34V, V36I, D38E, P39S, V40K, G41F, S42P, E43P, P44A, R45D, S46P, G47D, T48L, A49S, A73V, V74Y, A75N, N76Y, E77T, A78T |
| **V2D7r** | A31G, D32P, A34V, V36I, D38E, P39S, V40K, G41F, S42P, E43P, P44A, R45D, S46P, G47D, V74Y, A75N |
| **V2D8** | A28G, L29Q, A31G, D32P, D33Q, A34V, T35V, V36I, D38E, P39S, V40K, G41Q, S42P, E43P, P44T, R45D, S46P, G47D, T48L, A49S, A73V, V74Y, A75N, E77T, A78T |
| **V2D8r** | A31G, D32P, V36I, D38E, P39S, V40K, G41Q, S42P, E43P, P44T, R45D, S46P, G47D, V74Y, A75N |
| **V2D9** | F30L, D32S, A34V, V36L, E37Y, D38E, P39T, V40S, G41Q, S42D, E43R, P44T, R45Y, S46T, T48N, A49S, A73S, V74N, A75R, E77T |
| **V2D9r** | E37Y, D38E, P39T, V40S, G41Q, S42D, E43R, P44T, R45Y, S46T, V74N, A75R |
| **V2D10** | A28G, L29Q, A31G, D32P, D33Q, A34V, V36L, E37V, D38E, P39S, V40K, A(G)41S, S42P, E43P, P44A, R45D, S46P, A(G)47D, T48L, A49S, A73V, V74S, A75Q, E77T, A78T |
| **V2D11** | A28G, L29Q, A31G, D32P, D33Q, A34V, V36L, E37W, D38E, P39S, V40K, G41Y, S42P, E43P, P44A, R45D, S46P, G47D, T48L, A49S, A73V, V74S, A75G, A78T |
| **V2D11r(1)** | A31G, D32P, A34V, V36L, E37W, D38E, P39S, V40K, G41Y, S42P, E43P, P44A, R45D, S46P, G47D, T448L, A49S, V74S, A75G |
| **V2D11r(2)** | A31G, D32P, A34V, E37W, D38E, P39S, V40K, G41Y, S42P, E43P, P44A, R45D, S46P, G47D, T448L, A49S, V74S, A75G |

**Supplemental Table 10: Experimental characterization of designs from PIP version 1.**

| Design | Mutations | Expression | Purified | Solubility | Activity |
|---|---|---|---|---|---|
| **V1D1r** | 14 | Insoluble | Yes | Soluble | - |
| **V1D2r** | 18 | Insoluble | Yes | Insoluble | N/A |
| **V1D3r** | 11 | Insoluble | Yes | Soluble | - |
| **V1D4** | 28 | Insoluble | Yes | Insoluble | N/A |
| **V1D5r** | 15 | Insoluble | Yes | Insoluble | N/A |
| **V1D6** | 31 | None | No | N/A | N/A |
| **V1D7** | 32 | Insoluble | No | N/A | N/A |
| **V1D8r** | 19 | Insoluble | Yes | Soluble | + |
| **V1D9r** | 15 | Insoluble | Yes | Soluble | + |
| **V1D10r** | 14 | Insoluble | Yes | Soluble | + |
| **V1D11** | 29 | Insoluble | Yes | Insoluble | N/A |
| **V1D12r** | 16 | None | No | N/A | N/A |
| **V1D13** | 29 | Insoluble | Yes | Insoluble | N/A |
| **V1D14** | 29 | None | No | N/A | N/A |

Mutations: Number of mutations from wild-type KSI, excluding deletions. Expression: Whether the design expressed in inclusion bodies ("Insoluble") or not at all ("None"). Purified: Whether or not the design was successfully purified. Solubility: Whether the purified design was soluble after re-folding from inclusion bodies. Activity: Whether the design had any observable enzymatic activity ("+") or not ("-"); N/A: not applicable as protein could not be purified or was not soluble after purification.

**Supplementary Table 11. Comparison of median RMSD of lowest energy models on perturbed structures.** References for the different methods are indicated. Bold numbers denote best performance for given dataset (excluding FKIC with homologous fragments).

| Method | 8 residue side chain perturbed (Å) | 12 residue side chain perturbed (Å) | template based models (Å) | 12 residue backbone perturbed (Å) | 12 residue backbone perturbed no unavoidable clashes (Å) | 12 residue backbone perturbed unavoidable clashes (Å) |
|---|---|---|---|---|---|---|
| **HLP[2] *** | 2.2 | 2.25 | - | - | - | - |
| **HLP-SS[6] *** | 0.85 | 1.15 | - | - | - | - |
| **NGK[7] *** | **0.4** | 0.75 | 3.9 | - | - | - |
| **Galaxy-PS1[8] *** | 1.45 | 3.05 | 3.5 | - | - | - |
| **Galaxy-PS2[3] *** | 1.05 | 1.55 | **3.3** | **1.65** | 1.4 | **1.8** |
| **FKIC** | 0.45 | **0.64** | 3.9 | 1.68 | **1.28** | 2.59 |
| **FKIC with homologous fragments** | 0.42 | 0.54 | - | - | - | - |

* Values reported by ref[3].

**Supplementary Table 12.**

| *Structure* | *V1D8r (6UAD)* | *V2D9r (6UAE)* |
|---|---|---|
| *Wavelength* | 1.116Å | 1.116Å |
| *Resolution Range* | 46.03-1.75 (1.80-1.75) | 105.00-1.93 (1.96-1.93) |
| *Unit Cell* | a=b=53.15Å , c=178.03Å **α**=**β**=90° ***γ***=120° | a=73.01Å , b=210.00Å , c=39.64Å **α**=**β**=***γ***=90° |
| *Space Group* | $P6_522$ | $P2_12_12$ |
| *Unique Reflections* | 15833 (1048) | 46239 (2182) |
| *Multiplicity* | 9.7 (3.9) | 19.1 (17.7) |
| *Completeness* | 99.2% (92.0%) | 98.1% (94.2%) |
| *<I/σI>* | 34.2 (4.0) | 9.3 (1.0) |
| $CC_{1/2}$[9] | 1.000 (0.939) | 0.995 (0.652) |
| $R_{pim}$[10] | 0.013 (0.159) | 0.059 (0.818) |
| $R_{work}$[11] | 0.1742 (0.2083) | 0.1752 (0.3041) |
| $R_{free}$[11] | 0.2095 (0.2852) | 0.2122 (0.3916) |
| *Total Refined Atoms* | 1244 | 4662 |
| *Protein Residues* | 121 | 495 |
| *Solvent Molecules* | 163 | 372 |
| *Refined Ligand Atoms* | 66 | 204 |
| *Average B-factor* | 24.8Å$^2$ | 37.7Å$^2$ |
| $RMSD_{bonds}$ | 0.014Å | 0.006Å |
| $RMSD_{angles}$ | 1.23° | 0.91° |
| *Rama. Plot:* | | |
| *Favored* | 99.2% | 99.0% |
| *Allowed* | 0.8% | 1.0% |
| *Outliers* | 0.0% | 0.0% |
| *Molprobity Clashscore*[12] | 0.93 | 4.64 |
| *PDB ID* | 6UAD | 6UAE |

### Supplementary Table 13. Summary of energy function comparisons

| Dataset | Sampling method | Rosetta energy function | Median RMSD of lowest energy model (Å) | Median RMSD of lowest RMSD model (Å) | Median RMSD all models (Å) | Median sub-Å fraction | Median time (s) |
|---|---|---|---|---|---|---|---|
| ***Standard*** | NGK | talaris2013 | 0.74 | 0.37 | 2.71 | 11.40% | 2281 |
| ***Standard*** | NGK | ref2015 | 0.64 | 0.37 | 2.70 | 13.00% | 3642 |
| ***Standard*** | FKIC | talaris2013 | 0.70 | 0.36 | 1.19 | 44.89% | 1646 |
| ***Standard*** | FKIC | talaris2014 | 0.64 | 0.35 | 1.27 | 46.20% | 1813 |
| ***Standard*** | FKIC | ref2015 | 0.62 | 0.32 | 1.16 | 47.80% | 3456 |
| ***Mixed*** | NGK | talaris2013 | 1.94 | 0.45 | 4.66 | 1.90% | 3788 |
| ***Mixed*** | NGK | ref2015 | 1.07 | 0.45 | 4.65 | 1.15% | 7341 |
| ***Mixed*** | FKIC | talaris2013 | 0.61 | 0.34 | 1.74 | 34.60% | 3654 |
| ***Mixed*** | FKIC | ref2015 | 0.53 | 0.34 | 1.46 | 52.30% | 7196 |

### SUPPLEMENTARY FIGURES

#### Supplementary Figure 1. Detailed FKIC protocol.

The FKIC / LHKIC modeling protocol has a build stage (yellow), a centroid sampling stage (light red) and a full atom sampling stage (red). Both the centroid stage and the full atom stage perform simulated annealing which ramp the *rama* and *fa_rep* terms of the Rosetta energy function[5,13] in outer cycles and ramp the temperature in inner cycles.

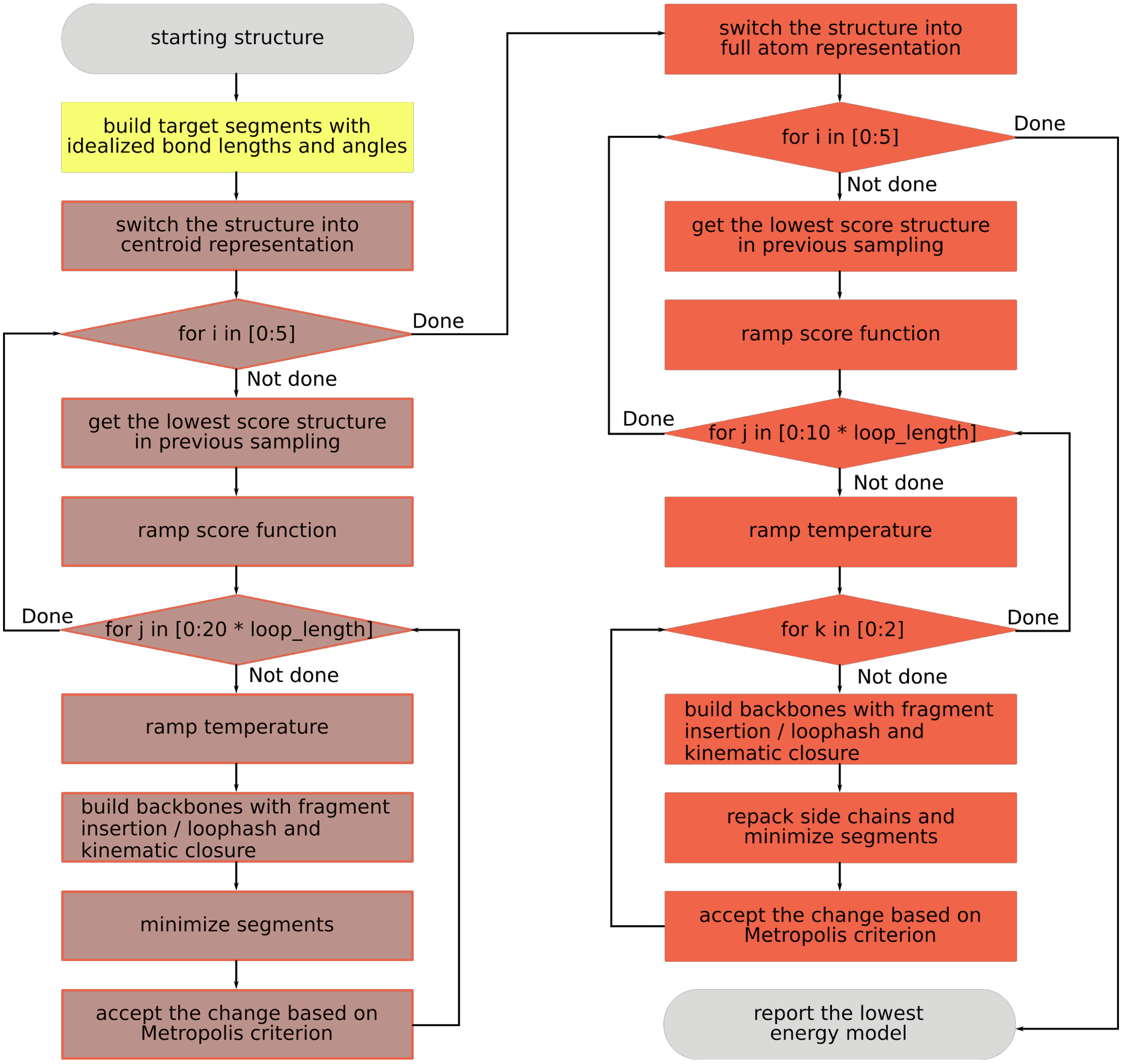

**Supplementary Figure 2. Examples of failures of FKIC.**

Results of standard FKIC are shown in red (right in each panel) and results of FKIC with native input information and native bond lengths and angles are shown in green (left in each panel). Each point represents a Rosetta generated model. REU, Rosetta energy units. (**a**) Sub-Å models are generated only with native inputs. (**b**) Standard FKIC generates a few sub-Å models but they are not identified by energy. Using native inputs generates a larger number of near-native solutions that can be correctly identified by energy. (**c**) The simulation with native inputs correctly identifies native-like models, but a model generated by standard FKIC has lower energy. (**d**) Neither standard nor native-input simulations correctly identify sub-Å models.

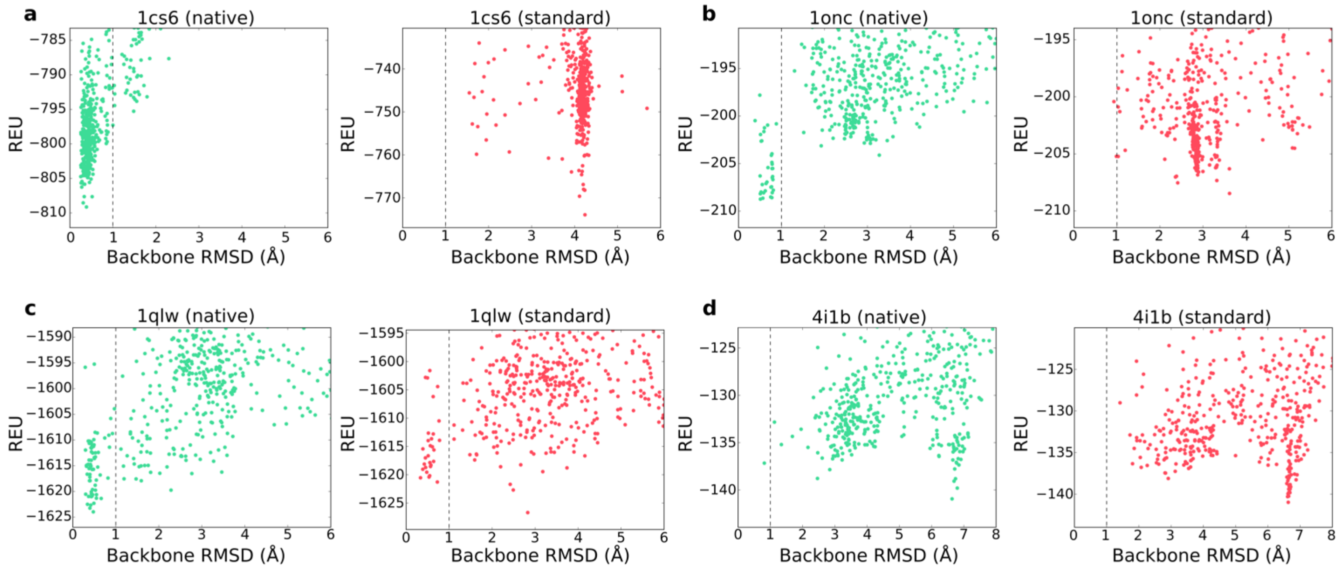

**Supplementary Figure 3. Selection of designs for via Pareto fronts in PIP version 2.**

Designs picked for computational structure prediction. Plots of design Lennard-Jones attractive (fa_attr) Rosetta score, in REU, vs. restraint satisfaction (longest distance of any restrained atom to its ideal position, in Å (**Methods**)) for the first (**a**), second (**b**), and third (**c**) iterations of design for PIP version 2. Designs chosen via Pareto fronts for structure prediction are shown in red, other designs in blue.

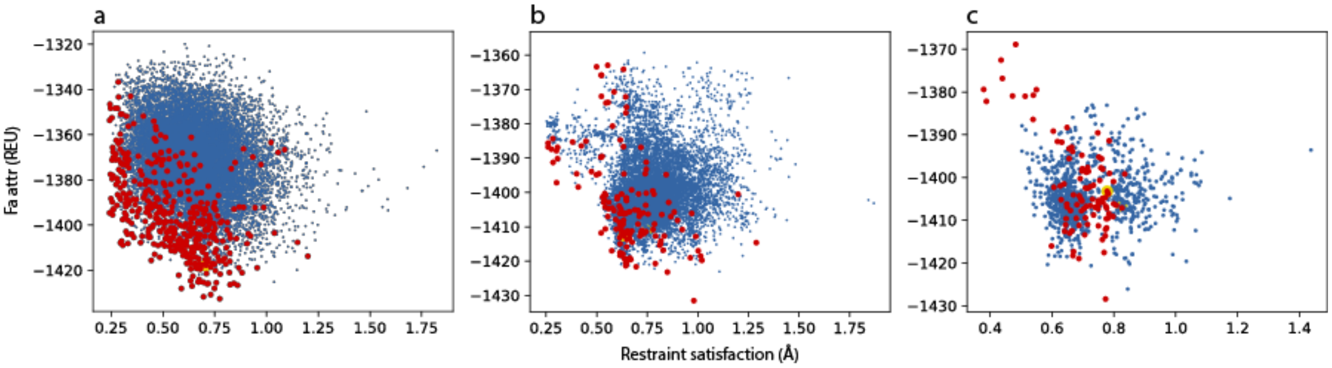

**Supplementary Figure 4 Computational structure prediction of designs from PIP V1.**

Rosetta total score (in REU) versus loop backbone RMSD (in Å) for designs selected for experimental testing from PIP version 1. Vertical dashed lines indicate 1Å loop RMSD. The experimentally characterized V1D8r design is highlighted in yellow. Structure prediction was performed on both the initial designs and the reversion mutants.

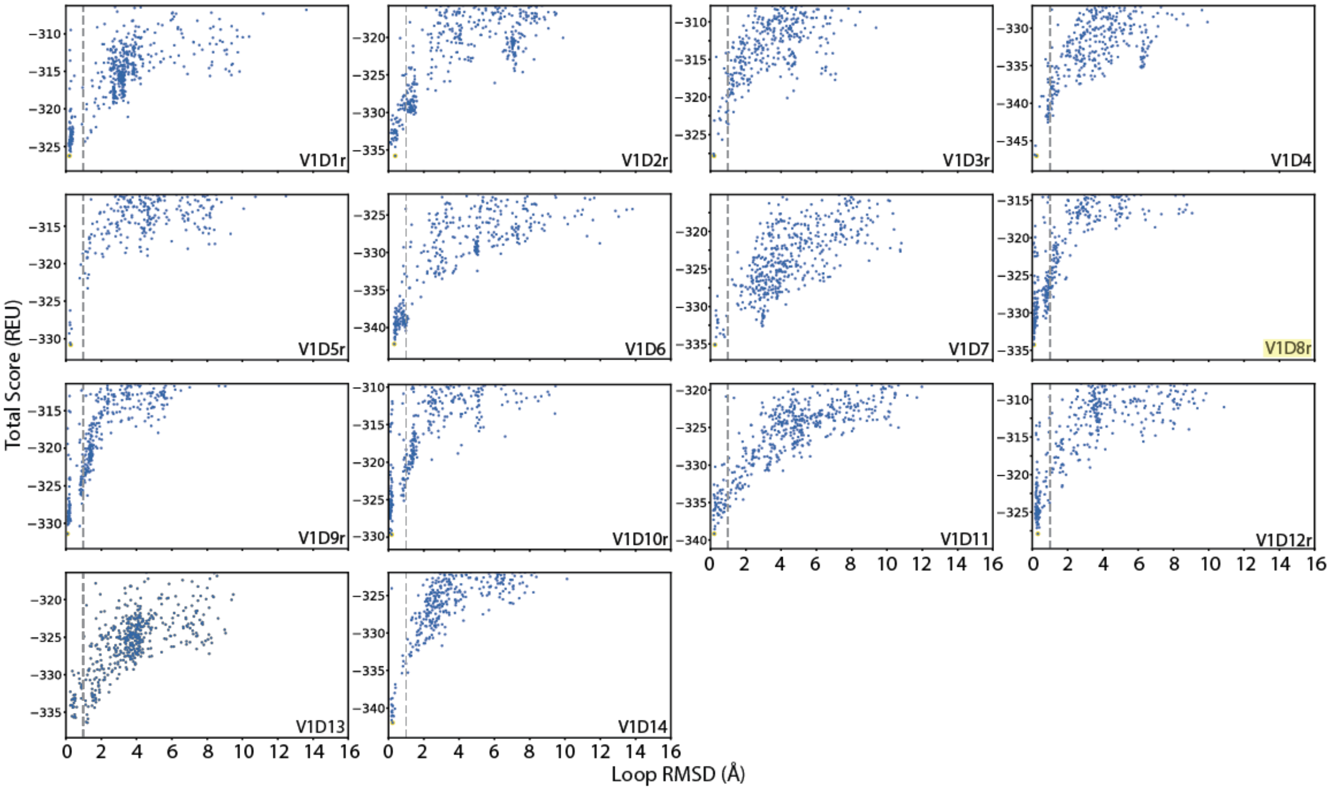

**Supplementary Figure 5. Computational structure predictions of designs from PIP V2.**
Rosetta total score (in REU) versus loop backbone RMSD (in Å) for designs selected for experimental testing from PIP version 2. Plots are shown for the designs excluding the reversion mutants. Vertical dashed lines indicate 1Å loop RMSD. V2D9, the design corresponding to the experimentally characterized V2D9r reversion mutant, is highlighted in yellow.

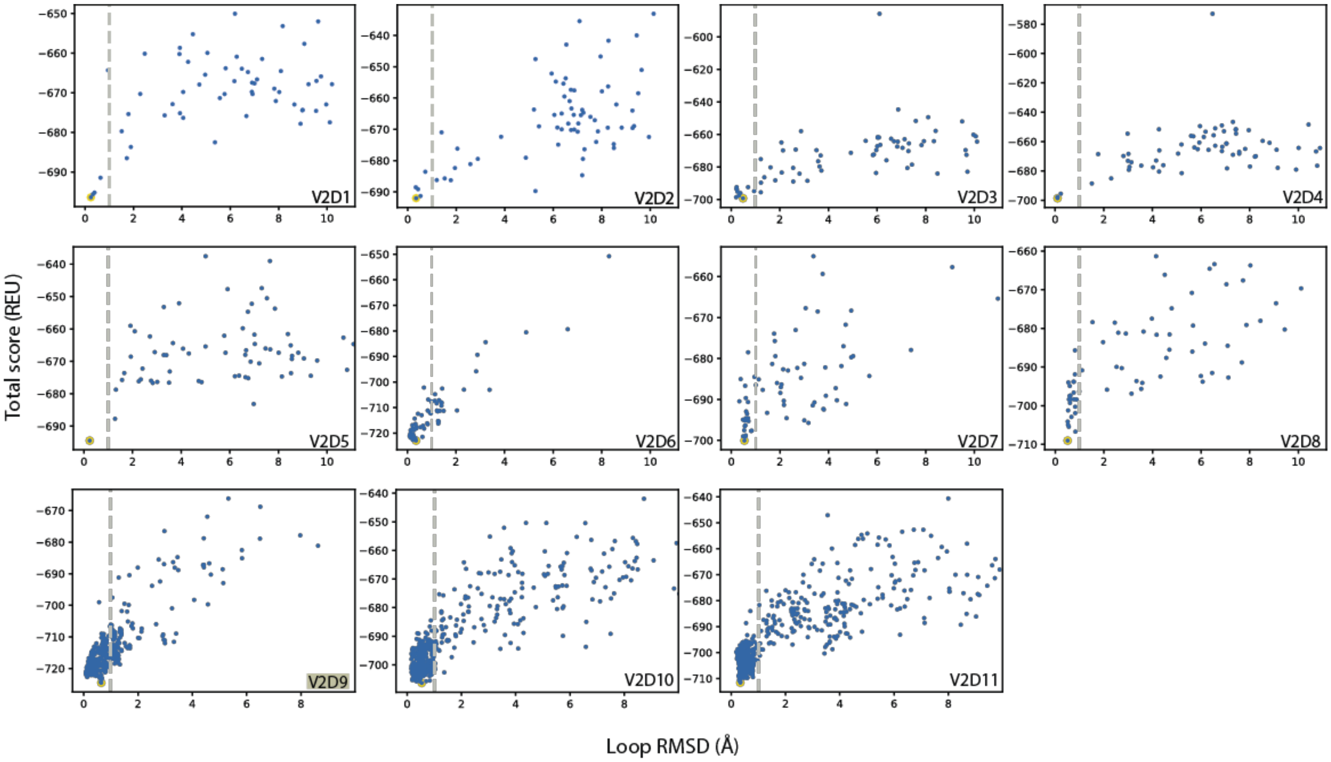

**Supplementary Figure 6. Biophysical and structural characterization of designs.**

Circular dichroism (CD) spectra for wild-type (**a**), V1D8r (**b**), or V2D9r (**c**). (**d**) Normalized temperature melting curves, measured via CD at 222 nm. V1D8r Tm: 43.5 C, V2D9r: 67.3 C. (**e**) Electron density of possible alternate conformations of E38 observed in design 2. Density is contoured at 0.5 sigma for residues 37-39 in chain B. Residue E38 (teal) and equilenin (purple) are shown as sticks.

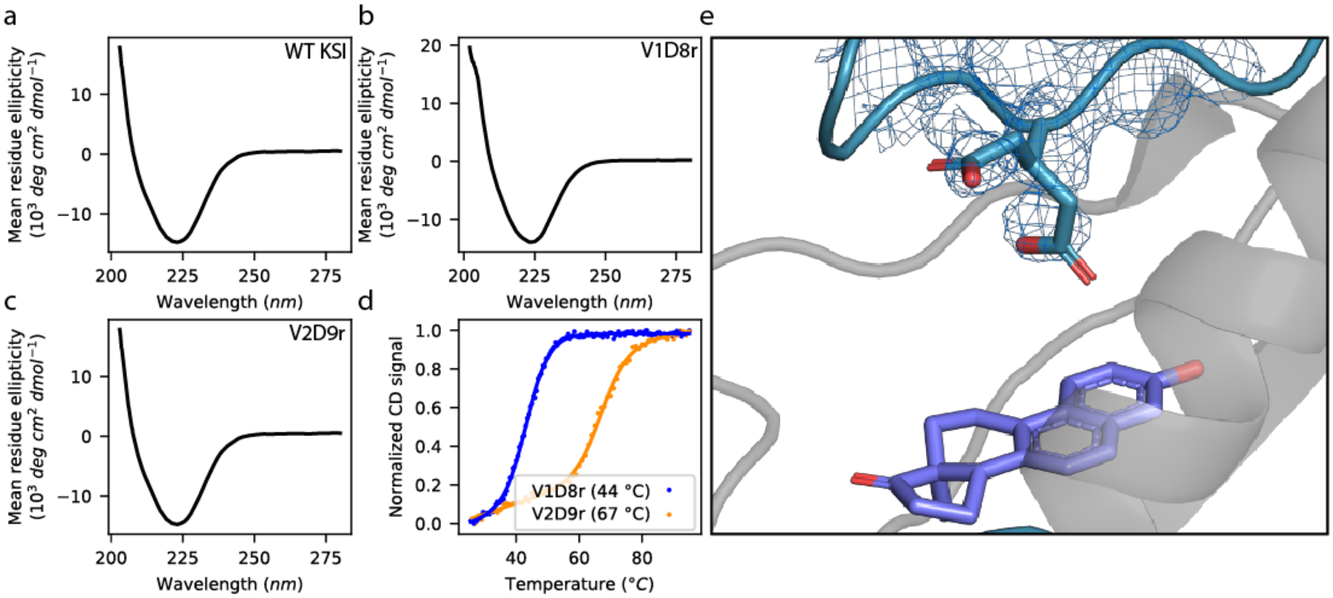

**Supplementary Figure 7. Analytical size exclusion chromatography (SEC) of KSI designs.**

Normalized absorbance (280 nm) from analytical SEC of V1D8r, V2D9r, wild-type KSI or a standards mixture with molecular weights as labeled.

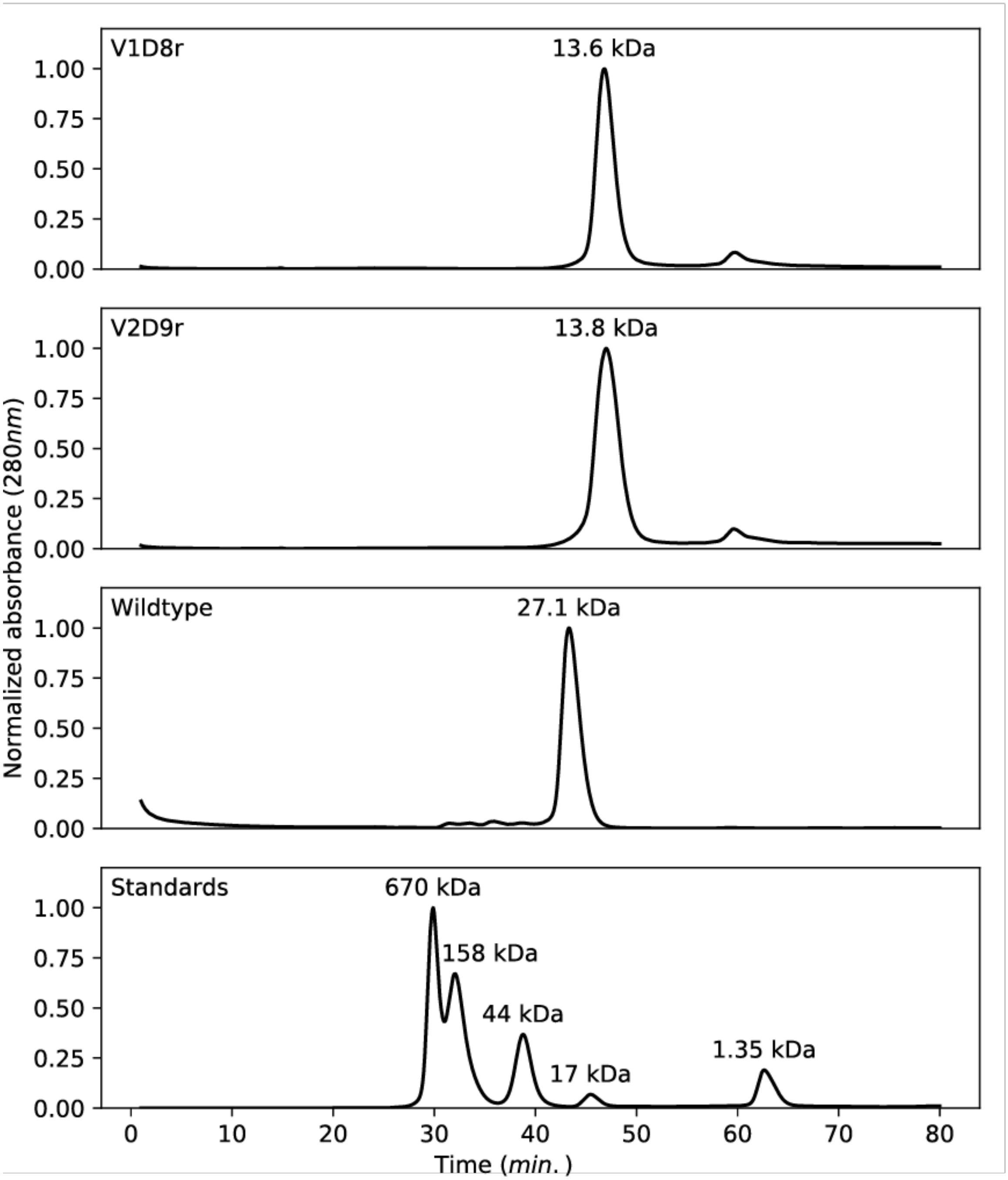
